# PMRIPrep: automated preprocessing and quantification of DSC and DCE MRI

**DOI:** 10.64898/2026.11.17.767132

**Authors:** Aaron M. Rulseh, Archith Rajan, Brandon Curl, Suyash Mohan, Josef Vymazal, Sanjeev Chawla

## Abstract

While dynamic susceptibility contrast (DSC) MRI measures hemodynamics and dynamic contrast-enhanced (DCE) MRI characterizes vascular permeability, quantitative values from both modalities remain processing-dependent. We present PMRIPrep, a Brain Imaging Data Structure (BIDS) App for automated DSC and DCE preprocessing and quantification. PMRIPrep adapts to available acquisitions and metadata, integrates both modalities within a session, and supports four DSC deconvolution methods and four DCE pharmacokinetic models. In DSC simulations with 4% cerebral blood volume and signal-to-noise ratio 50, pooled median absolute cerebral blood flow error was 13–19% across methods. Median K^trans^ bias remained ≤2% across four model-matched DCE configurations over 0–10% noise. End-to-end OSIPI analyses yielded DCE accuracy and clinical test-retest repeatability scores of 0.85 and 0.85, respectively. Outputs follow BIDS-Derivatives conventions and include machine-readable provenance and visual quality-control reports. PMRIPrep is freely available as a containerized application.

## Introduction

The most commonly used perfusion-weighted imaging techniques for characterizing brain tumor vasculature include dynamic susceptibility contrast (DSC) and dynamic contrast-enhanced (DCE) MRI. DSC MRI measures hemodynamic parameters such as cerebral blood flow (CBF), cerebral blood volume (CBV), and mean transit time (MTT) by tracking the first-pass bolus of gadolinium-based contrast agents through the microvasculature [1]. DCE MRI characterizes blood-brain barrier permeability by modeling the extravasation of gadolinium-based contrast agents from intravascular space into the extravascular extracellular space, yielding metrics such as the volume transfer constant K^trans^, extravascular extracellular volume fraction (v_e_), and plasma volume fraction (v_p_) [2,3]. Quantitative and semi-quantitative parameters derived from DSC and DCE are used widely for tumor characterization, grading and treatment-response assessment [4–6]; however, processing-dependent variation limits comparison across studies and software platforms.

Quantitative values from DSC and DCE remain highly sensitive to processing implementation. In a controlled multisite comparison using identical inputs, inter-software coefficient of variation for normalized CBV averaged 30% in permeable tissue [7]; normalized-CBV test-retest variation reached up to 67% depending on postprocessing method [8]. Arterial input function (AIF) selection, deconvolution, tracer-kinetic model, leakage correction, and signal-to-concentration assumptions are key contributing factors to variability [3,7,9]. The Quantitative Imaging Biomarkers Alliance (QIBA) has developed acquisition and quantitative-analysis profiles for DSC and DCE MRI; moreover, the International Society for Magnetic Resonance in Medicine (ISMRM) Open Science Initiative for Perfusion Imaging (OSIPI) has developed standardized terminology and public benchmarking challenges for perfusion analysis [10–13]. However, an integrated open-source perfusion imaging framework spanning both DSC and DCE protocols is still lacking.

fMRIPrep established reproducible, auditable preprocessing for functional MRI within the NiPreps (NeuroImaging PREProcessing toolS) ecosystem [14]. This model was subsequently extended to arterial spin labeling with ASLPrep [15] and diffusion MRI with QSIPrep [16]. There is a critical need to develop similar pipelines for processing DSC and DCE MRI data as currently available tools are commonly vendor-specific, limited to one perfusion modality, dependent on proprietary platforms, or reliant on manual intervention.

We present PMRIPrep, a Brain Imaging Data Structure (BIDS) App designed for the automated preprocessing and quantification of DSC and DCE MRI. PMRIPrep reads acquisition metadata from BIDS sidecars to configure appropriate processing workflows without manual parameter specification. It supports four DSC deconvolution methods and four DCE pharmacokinetic models and generates visual reports for quality control. PMRIPrep is open-source under the Apache License, Version 2.0, and distributed as Docker and Apptainer containers.

## Results

### Automated multimodal perfusion workflow

PMRIPrep constructs each workflow from the available BIDS modalities, acquisition metadata, and user configuration (Fig. 1). Preflight checks identify missing or inconsistent metadata required for quantitative processing before execution. Anatomical preprocessing supplies a common subject reference and brain mask, with tissue segmentation masks generated when an eligible precontrast T1-weighted (T1w) image is available. DSC and DCE data are quantified in their respective native perfusion spaces without resampling to standard space. DSC processing uses a measured AIF when detection succeeds; it otherwise retains AIF-independent outputs and withholds AIF-dependent maps. Boxerman-Schmainda-Weisskoff (BSW) leakage correction can be enabled or explicitly disabled. DCE uses measured transmit-field (B1+) and T1 information when present and records the population assumptions used otherwise; a population AIF is used when no suitable subject-specific AIF is available.

**Figure 1.**
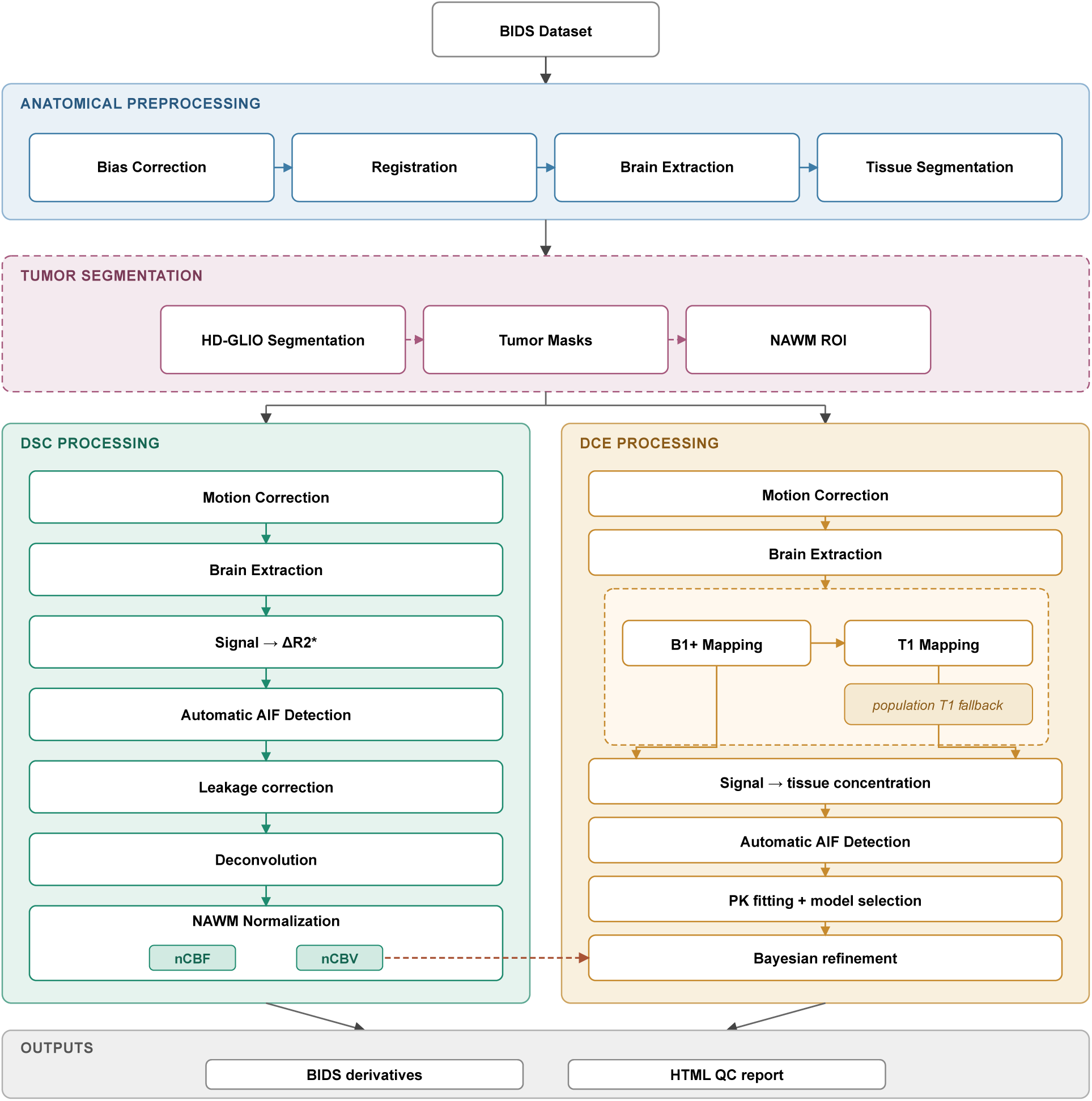
PMRIPrep pipeline overview. BIDS inputs undergo anatomical preprocessing to generate a common anatomical reference and brain mask, with tissue segmentation performed when an acceptable precontrast T1w image is available. Glioma segmentation and NAWM construction are added when the required structural images are available. The DSC pathway performs motion correction, signal-to-ΔR2* conversion, AIF detection, optional leakage correction, AUC-based rCBV calculation, and deconvolution. The DCE pathway performs motion correction, B1+ and T1 mapping when available, signal-to-concentration conversion, AIF detection, and pharmacokinetic modeling. Same-session DSC nCBV can supply a spatially varying v_p_ prior during Bayesian DCE refinement. Dashed enclosures denote modality-dependent extensions, and the dashed red arrow denotes cross-modal transfer. Outputs comprise BIDS-compatible derivatives, JSON provenance, and a per-session HTML QC report.

When the required anatomical images are present, PMRIPrep generates tumor and normal-appearing white matter (NAWM) masks. User-supplied lesion, NAWM, and region-of-interest (ROI) masks are also accepted. These masks provide regional summaries and allow separate NAWM normalization for DSC-DCE concordance. The DSC-derived NAWM-normalized CBV (nCBV) map can additionally define a spatially varying v_p_ prior during optional Bayesian DCE refinement. Derivatives record the relevant processing choices and assumptions, and the session report distinguishes completed, skipped, fallback, and failed operations. Representative quality-control (QC) report figures are presented in Supplementary Figure S2.

### DSC quantitative performance

Standard singular value decomposition (sSVD) [17], oscillation-index SVD (oSVD) [18], Tikhonov regularization [19], and control point interpolation (CPI) [20] were evaluated on identical simulated DSC observations generated using an independently implemented forward model. Following the ranges of Mouridsen et al. [21], full-integral CBV was fixed at 4% and CBF varied from 10 to 70 mL/100 g/min (MTT 24–3.43 s). Exponential, triangular, and box residues were evaluated without delay under noiseless conditions and at signal-to-noise ratios (SNR) of 100, 50, and 20 (Fig. 2a-c; Supplementary Section A).

**Figure 2.**
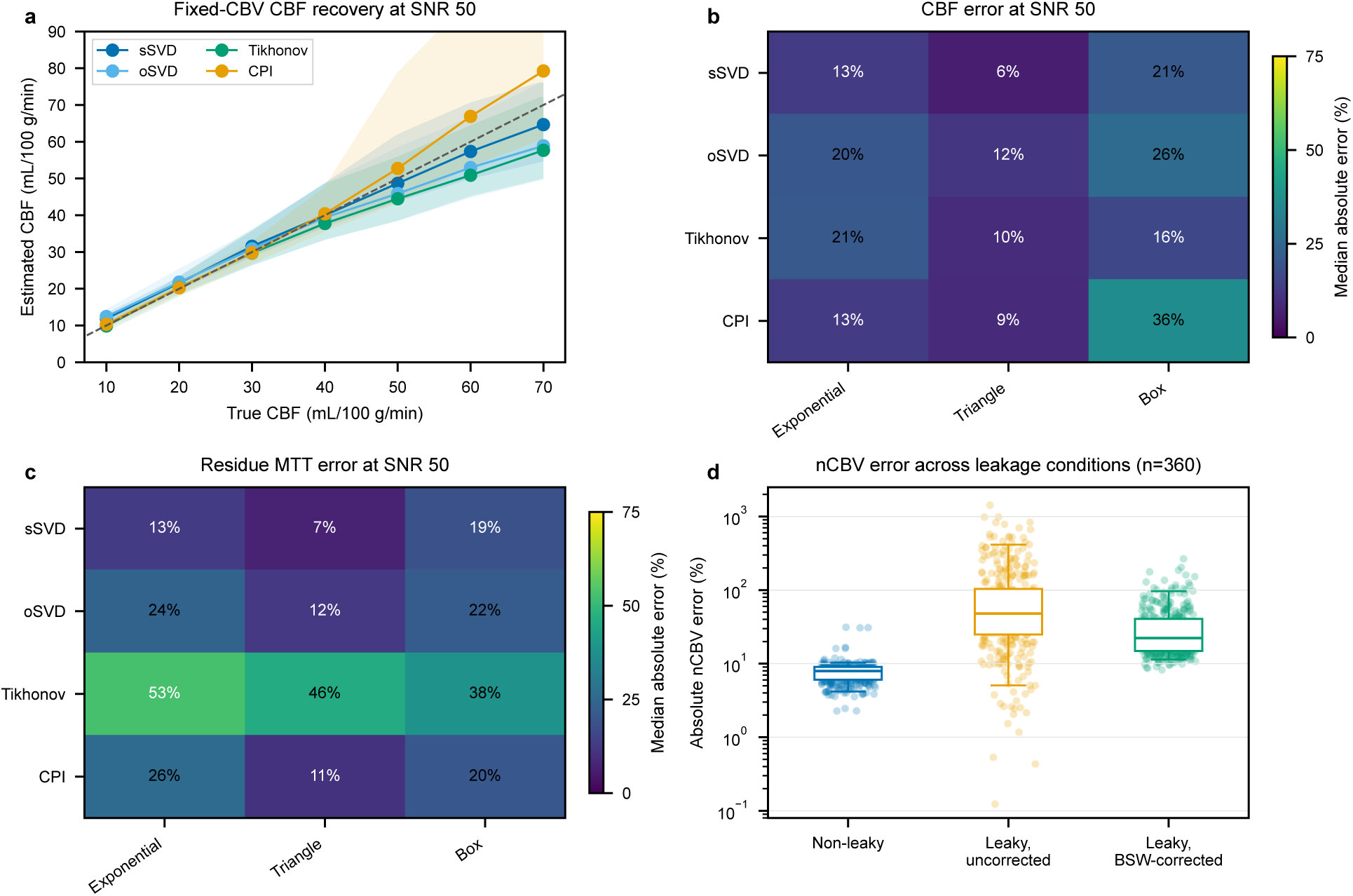
DSC-MRI deconvolution and leakage-correction validation. **a**, Estimated versus simulated CBF for sSVD, oSVD, Tikhonov and CPI. Lines and shading show medians and interquartile ranges across exponential, triangular and box residues; the dashed line indicates identity. **b,c,** Median absolute percentage errors in CBF (b) and residue-integral MTT (c), pooled across flow values within each residue family. Panels a-c use CBV of 4%, baseline-signal SNR 50 and 200 realizations per flow-residue combination. **d,** Absolute percentage error in nCBV from the GBM-DSC-MRI-DRO voxelwise concentration-curve analysis, comparing non-leaky tumor, uncorrected leaky tumor and BSW-corrected leaky tumor (360 acquisition configurations per condition). Boxes show medians and interquartile ranges, whiskers the 5th-95th percentiles, and points individual configurations.

At baseline-signal SNR 50, pooled median absolute percentage error in CBF was 13% for sSVD, 19% for oSVD, 15% for Tikhonov, and 13% for CPI. Median absolute residue-MTT error among finite estimates was 13%, 19%, 46%, and 17%, respectively. Tikhonov residue MTT was unavailable in 621 of 4,200 observations; the other methods returned finite residue MTT throughout. The corresponding reported-MTT errors were 13%, 19%, 15%, and 17%; for the SVD methods MTT is calculated as area under the curve (AUC)-based CBV divided by CBF, whereas CPI reports fitted-residue MTT.

Recovery depended on flow, residue family, and SNR. At SNR 50, CPI’s median CBF error was 9% for triangular residues, 13% for exponential residues, and 36% for box residues. Its joint leakage fit recovered the K2 sign for leakage generated by the BSW model, but gave the wrong sign for T2*-dominant exchange leakage when tissue transit was prolonged (Supplementary Section B).

On the external GBM-DSC-MRI digital reference object (DRO) [22,23], voxelwise concentration-curve analysis across 360 acquisition configurations gave median non-leaky tumor nCBV bias of −8% (interquartile range, −9% to −6%). For leaky tumor, uncorrected bias was −10% (−104% to +33%). BSW correction narrowed this distribution but retained a negative bias and tail (median −22%; interquartile range, −41% to −15%), without restoring ground-truth nCBV (Fig. 2d). In a separate analysis that averaged the magnitude signal within each region before conversion, BSW-corrected non-leaky tumor nCBV bias was +7.0% (interquartile range, +2.9% to +10.9%). In this ROI-averaged analysis, CPI median NAWM-normalized CBF (nCBF) bias was −1.1% (interquartile range, −19.3% to +19.7%), with tumor and NAWM MTT biases of −7.8% and −11.7%, respectively (Supplementary Tables S5a-1 and S5b).

### DCE quantitative performance

Tofts, Extended Tofts, Patlak, and the two-compartment exchange model (2CXM) were assessed on data matching their respective generative assumptions, separating estimator recovery from model selection and cross-model behavior. On a synthetic grid spanning the QIBA v9 K^trans^ × v_e_ values [24], most Tofts cells were recovered within ±0.5%; error increased with undersampled high efflux rate constant k_ep_ and reached +35% (Fig. 3a). On the noise-free QIBA v9 DRO at 6-s sampling, median K^trans^ and v_e_ biases were within 2% in the higher-signal configurations. At 10-s sampling, median K^trans^ bias ranged from approximately −4% to +6%, depending on sample timing relative to the AIF peak. The lower-signal-scale and low signal-to-noise ratio configurations had substantially larger errors (Supplementary Section F). Extended Tofts tracked true v_p_ from 0.01 to 0.08 with K^trans^ and v_e_ within approximately 1% in the tested configurations (Fig. 3b); on the shared noise-free QIBA v4 DRO, mean absolute errors were 0.0018 min⁻¹ for K^trans^, 0.032 for v_e_, and 0.00071 for v_p_ (Supplementary Section G).

**Figure 3.**
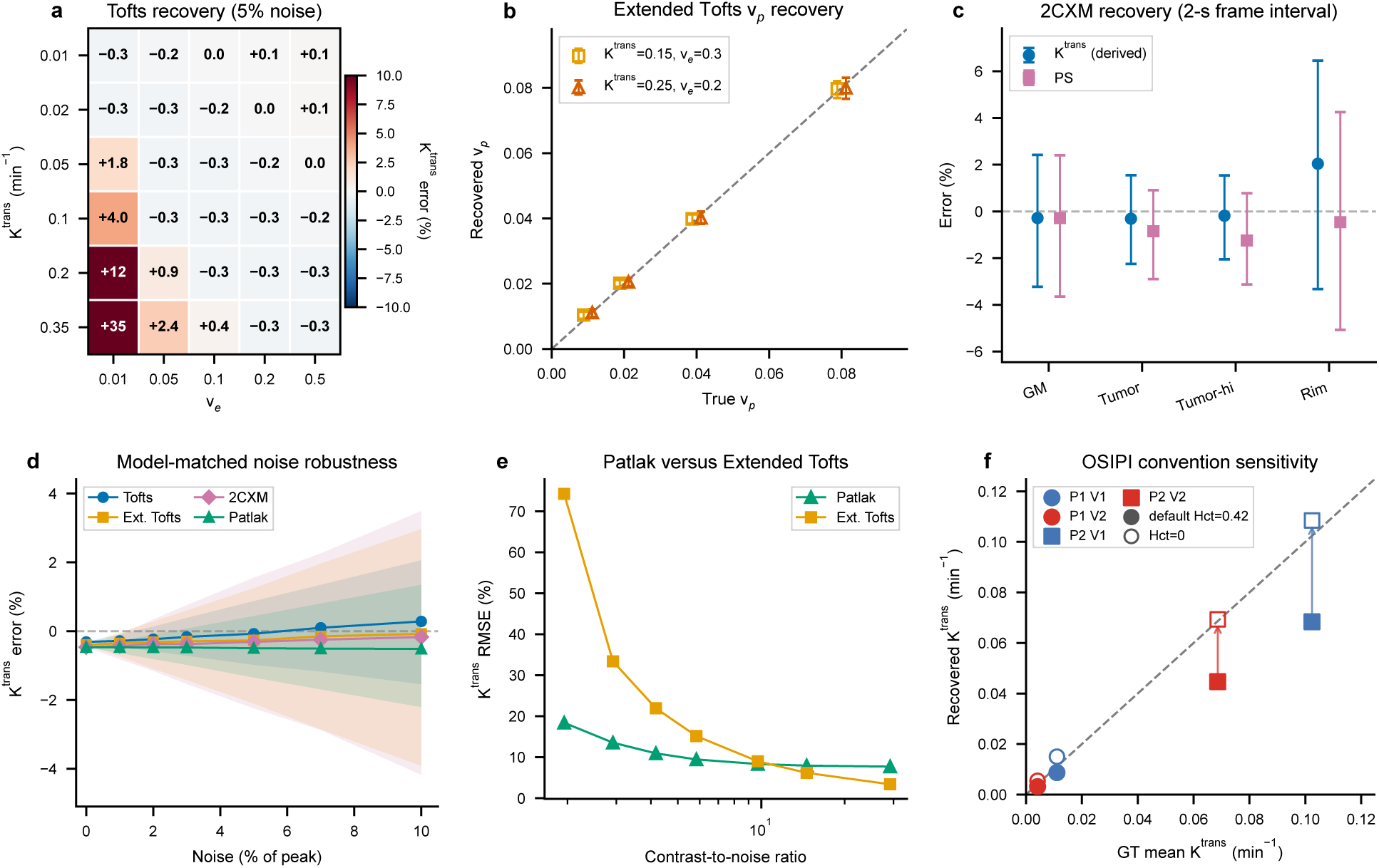
DCE-MRI pharmacokinetic model performance. **a**, Tofts K^trans^ error across a synthetic grid spanning the QIBA v9 K^trans^ and v_e_ values at 5% noise. Delay estimation increased bias in the two fastest-washout cells; with delay fixed at its true value of zero, median bias was within 2%. **b,** Extended-Tofts recovery of v_p_ in two model-matched configurations at 5% noise. **c,** Error in derived K^trans^ and permeability-surface-area product across four 2CXM configurations at a temporal resolution of 2 s and 5% noise. **d,** K^trans^ error across 0–10% noise for model-matched Tofts, Extended-Tofts, Patlak, and 2CXM simulations. **e,** K^trans^ root-mean-square error for Patlak and Extended Tofts in NAWM-like reversible tissue across contrast-to-noise ratio. **f,** Challenge-ROI mean K^trans^ versus ground truth for four OSIPI synthetic sessions under the default analysis (filled markers) and hematocrit-zero sensitivity analysis (open markers), evaluated in finite voxels. Circles and squares denote the two synthetic subjects (P1 and P2), colors denote visits, and arrows connect paired results. Panels a-e use 200 Monte Carlo realizations per condition and effective-delay estimation. Heat-map cells show median values; in line plots, lines connect medians and shaded bands or error bars show interquartile ranges. Dashed lines indicate identity in panels b and f and zero error in panels c and d.

At a temporal resolution of 2 s, 2CXM recovered median derived K^trans^ and permeability-surface-area product within approximately 2% and v_e_ and v_p_ within 0.004 at 5% noise (Fig. 3c). Across 0–10% noise, median K^trans^ bias remained within approximately 2% in all four model-matched configurations (Fig. 3d). In NAWM-like tissue with low contrast-to-noise ratio (CNR), the Patlak model’s bias due to assumed negligible contrast-agent backflux was offset by lower variance than Extended Tofts, with the root-mean-square error (RMSE) curves crossing near CNR 10 (Fig. 3e). Effective-delay estimation reduced Extended-Tofts K^trans^ mean absolute error for a 5-s tissue delay from 0.021 to 0.002 min⁻¹ at 4-s temporal resolution, but could fit a nonzero delay where none was present, particularly at coarse temporal resolution (Supplementary Section K). Cross-model recovery is reported in Supplementary Section H; explicit-2CXM plasma-flow recovery across temporal resolutions and AIF-tissue delays is reported in Supplementary Section J.

End-to-end validation used the four OSIPI DCE synthetic sessions, processed from BIDS input through quantitative maps with automatic AIF detection and Bayesian information criterion (BIC)-based model selection. The primary analysis used all finite values within the challenge tumor ROIs, consistent with the OSIPI challenge convention [13], and achieved an aggregate accuracy score of 0.85. A hematocrit-zero sensitivity analysis matching the source phantom’s plasma-AIF convention reduced underestimation in P2, with a higher aggregate score of 0.92 (Fig. 3f). In a paired analysis restricted to voxels passing the DCE QC mask under both conventions, the corresponding scores were 0.86 and 0.90 (Supplementary Section I). The QC-filtered scores are presented as secondary results, as voxel filtering differs from the published challenge methodology.

### DCE test-retest reproducibility

Reproducibility was assessed using the OSIPI clinical DCE test-retest dataset: eight subjects with recurrent glioblastoma selected from the RIDER Neuro MRI collection and scanned twice on the same 1.5 T scanner, two days apart [13]. Each session was processed with default settings. The AIF source was consistent between visits for all eight subjects: six used a detected AIF at both visits, while two fell back to a population AIF at both visits.

Following the OSIPI-DCE challenge methodology, mean parameters were computed within the supplied tumor masks at both visits and each two-visit pair used its population standard deviation. Root-mean-square (RMS) within-subject coefficients of variation were 18.2% for K^trans^, 16.6% for v_e_, and 13.0% for v_p_ (Fig. 4b; Supplementary Table S9d). The corresponding K^trans^ repeatability coefficient was 50.4% (95% bootstrap interval, 36.0–62.3%). Bland-Altman analysis of tumor-mean K^trans^ showed a mean visit difference of −0.00274 min⁻¹ and 95% limits of agreement from −0.0252 to 0.0197 min⁻¹ (Fig. 4a).

**Figure 4.**
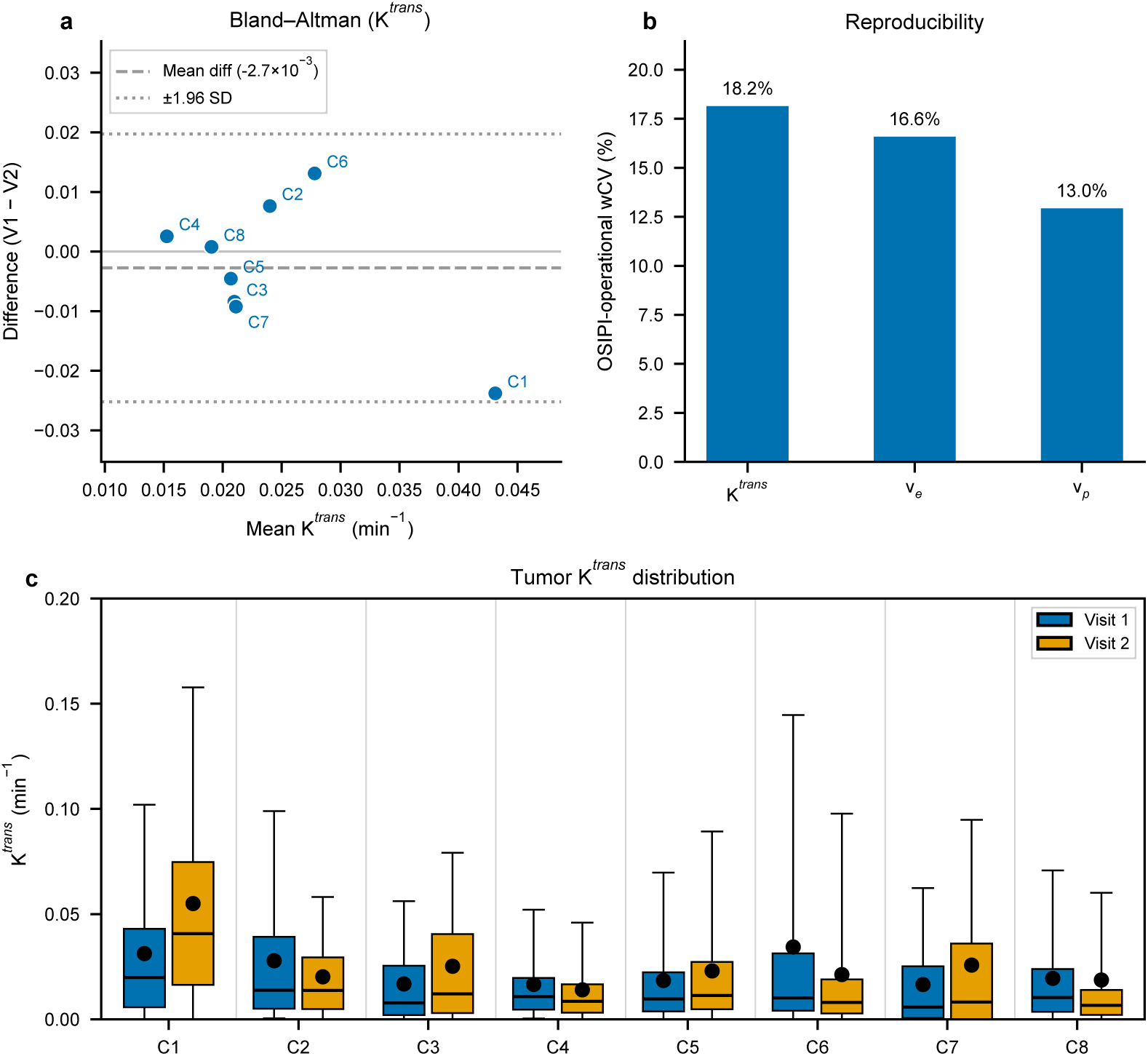
DCE-MRI test-retest reproducibility. **a**, Bland-Altman plot of tumor-mean K^trans^ across eight subjects with recurrent glioblastoma scanned at two visits. The dashed line is the mean visit difference (−0.00274 min⁻¹) and dotted lines are the 95% limits of agreement (−0.0252 to 0.0197 min⁻¹). **b,** RMS within-subject coefficients of variation (using the OSIPI challenge convention) were 18.2% for K^trans^, 16.6% for v_e_, and 13.0% for v_p_. **c,** Tumor K^trans^ distributions at visits 1 and 2; boxes show the interquartile range, center lines the median, whiskers the 5th-95th percentiles, and points the mean. The AIF source was concordant between visits for all eight subjects: six used a detected AIF at both visits and two used a fallback population AIF at both visits.

Using the OSIPI scoring methodology [13], the primary analysis achieved an accuracy score of 0.85, K^trans^ repeatability score of 0.85, and combined silver score of 72. A QC-filtered secondary analysis yielded 0.86, 0.89, and 77, respectively, and reduced the challenge-convention K^trans^ repeatability coefficient to 34.4% (95% bootstrap interval, 24.7–42.4%; Fig. 4; Supplementary Section I).

### Clinical demonstration and cross-modality DSC-DCE integration

An external-site dataset of 11 post-treatment glioblastoma cases with DSC and DCE acquisitions was used during development to evaluate generalization and generate representative outputs (Fig. 5); it was not treated as an independent accuracy-validation cohort. No variable-flip-angle (VFA) or B1+ maps were available, and therefore DCE processing used the uniform 3 T population T1 of 1200 ms. PMRIPrep completed automatic AIF detection, BIC-based model selection, lesion statistics, and same-session DSC-DCE registration in all cases.

**Figure 5.**
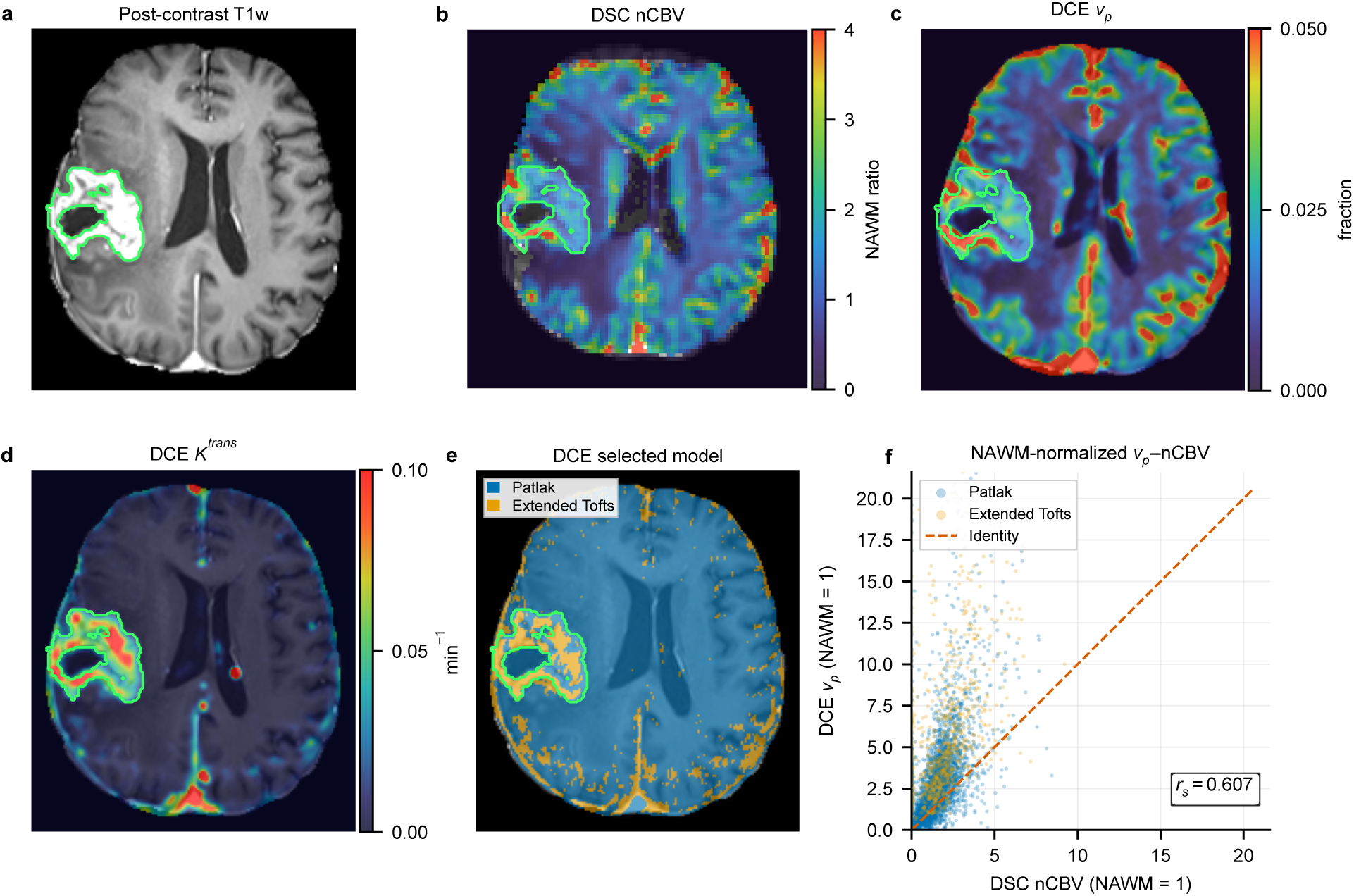
Representative PMRIPrep output in glioblastoma. **a**, Post-contrast T1-weighted image with the enhancing-tumor contour in green. **b,** NAWM-normalized DSC CBV after BSW leakage correction. **c,d,** DCE v_p_ and K^trans^ from automatic least-squares model selection. **e,** Retained DCE model; 78.2% of whole-brain voxels selected Patlak and 21.8% selected Extended Tofts. **f,** Independent descriptive concordance between separately NAWM-normalized DCE v_p_ and DSC CBV after matched Gaussian smoothing (σ = 1 mm). Whole-brain Spearman ρ was 0.607 across 426,854 jointly valid voxels; a fixed random sample of 5,000 voxels is shown and colored by retained model. The dashed line is identity. Panels a and c-e are in DCE space; panel b is in native DSC space.

The default least-squares estimator was used in all 11 cases; DCE fitting therefore did not apply the DSC-informed v_p_ prior. Descriptive concordance was independent of fitting: after separate within-subject NAWM normalization and matched Gaussian smoothing (σ = 1 mm), whole-brain Spearman ρ ranged from 0.43 to 0.61 and lesion-restricted ρ from 0.46 to 0.75. In one case, PMRIPrep was compared with Siemens Spectra (Tissue4D) inline maps as a cross-software illustration. Within enhancing tumor, Pearson r/Spearman ρ/Lin concordance correlation coefficient values were 0.85/0.91/0.84 for K^trans^, 0.87/0.83/0.70 for v_p_, 0.82/0.84/0.79 for v_e_, and 0.99/0.99/0.99 for NAWM-normalized CBV (Supplementary Fig. S1). v_p_ had a higher Pearson correlation but lower rank correlation and quantitative agreement than K^trans^. The undocumented Spectra fitting and masking settings preclude attributing differences to specific processing steps.

### DSC-informed DCE vp prior

In simulations, variational Bayesian inference (VI) with each model’s default priors reduced 2CXM RMSE relative to least-squares fitting by 54–79% across all fits and by 5–36% for K^trans^ and v_p_ in well-constrained fits; Extended-Tofts changes were small except for v_e_ (up to 39% lower), and Patlak error increased by up to 21%. Relative to default-prior VI, a DSC-informed prior with unbiased centers reduced v_p_ RMSE by up to 4.9% for Extended Tofts and 8.5% for Patlak, and reduced v_p_ and K^trans^ RMSE in well-constrained 2CXM fits by up to 26.6% and 19.9%, respectively. With priors derived from simulated DSC signal, Extended-Tofts v_p_ RMSE decreased by 1.4–1.8%, compared with 1.9% with priors set to true v_p_; with incorrect NAWM calibration, the benefit was smaller and no increase in error was detected. For 2CXM, no difference in v_p_ RMSE was detected between signal-derived priors and priors set to true v_p_ in well-constrained fits (Supplementary Section L, Tables S12a-h).

The 11 clinical cases were additionally processed in paired DCE VI analyses with the DSC-informed v_p_ prior enabled or skipped. The prior was used in a median of 65.8% of retained fits across subjects (range, 53.5–72.1%). VI reverted to least-squares estimates in a median of 3.9% of retained voxels with the prior enabled (range, 2.2– 4.9%), compared with 3.2% with the prior skipped (1.3–4.3%). Median enhancing-tumor values changed by −1.6% for v_p_, +0.5% for K^trans^, and −0.2% for v_e_, whereas v_p_ decreased by 16.2% in NAWM and 17.9% in non-enhancing tumor; median R² changed by −0.0015 (range, −0.0026 to +0.0016; Supplementary Section L).

## Discussion

PMRIPrep processes DSC and DCE MRI within the same BIDS-compatible workflow and records the inputs, methods, and assumptions used for each derivative. Validation showed that quantitative performance varied by both acquisition conditions and model assumptions, revealing that no estimator performed best across all tested settings.

PMRIPrep extends CPI by estimating leakage jointly with baseline signal and perfusion. Accuracy remains dependent on residue shape and the assumed leakage model. Optional dispersion correction improved recovery in some simulated conditions, while spatial regularization provided only small gains. The leakage extension broadens CPI’s applicability, although CPI did not show a general accuracy advantage over the SVD methods. For AUC-based CBV, BSW correction remains sensitive to differences between tissue and reference transit and to departures from the assumed leakage model.

Accurate DCE parameter recovery under matched model assumptions does not by itself establish that the generating model can be identified from noisy data. BIC balances fit quality against model complexity, favoring simpler models when additional parameters provide insufficient improvement. The Patlak model reduced variance at low CNR by assuming negligible backflux, at the cost of bias when that assumption was violated. Plasma-flow estimation with 2CXM was sensitive to temporal sampling and AIF width. PMRIPrep therefore combines BIC-based selection with a temporal-resolution restriction on 2CXM eligibility (Supplementary Sections H and J).

PMRIPrep uses automatic AIF detection by default, reducing reliance on manual AIF placement, a major source of inter-operator variability [9]. The atlas-to-whole-brain cascades record the selected tier and fallback path. While the AIF source was stable between OSIPI visits, the two cases requiring population fallback exhibited the largest K^trans^ coefficients of variation. Although this observation does not establish causality, the AIF source is recorded in QC outputs so its potential influence can be considered when interpreting quantitative results.

In simulations, the DSC-informed prior had model-dependent effects: small v_p_ improvements for Extended Tofts and Patlak, and larger improvements for explicit 2CXM, with small v_e_ increases at 10% noise in well-constrained fits. Benefits depended on temporal resolution, noise and prior calibration, and priors derived from simulated DSC signal gave benefits close to those of priors set to true v_p_. In the paired clinical VI analysis, the prior was used in most retained fits but showed tissue-dependent effects: enhancing-tumor estimates changed little, whereas v_p_ changed more in weakly enhancing tissue. Changes in DCE fit R² were negligible. The larger v_p_ changes in weakly enhancing tissue are consistent with greater prior influence when the DCE signal is less informative, but the absence of ground truth for the clinical data prevents assessing these changes as more accurate.

DSC-DCE concordance is independent under default least-squares fitting, which does not use DSC information. However, this independence is lost when DSC nCBV supplies the Bayesian v_p_ prior. PMRIPrep labels the latter comparison prior-informed and does not present it as independent validation. Finally, combining these hemodynamic and permeability measures can provide complementary information about tumor biology and the microenvironment [25].

Some aspects of PMRIPrep’s design merit discussion. DSC derived parameters (CBF and CBV) remain relative rather than absolutely calibrated. DCE quantification uses VFA-derived or precomputed T1 maps when available; otherwise uniform field-strength-dependent T1 is used. As concentration and pharmacokinetic estimates depend nonlinearly on T1 [2,26], this fallback may introduce tissue- and enhancement-dependent bias and is not equivalent to measured T1. Automated tumor segmentation uses HD-GLIO [27] and is limited to glioma, although user-supplied masks permit NAWM normalization and regional statistics for other pathologies. DSC currently assumes single-echo gradient-echo echo-planar imaging, and DCE assumes spoiled gradient-echo signal behavior. Validation included controlled internal simulations, external DSC and DCE digital reference objects, and OSIPI clinical test-retest data, but prospective multisite evaluation across scanners and protocols remains necessary.

PMRIPrep implements DSC and DCE processing in a single BIDS-compatible workflow. It retains processing choices in machine-readable provenance and exposes major decisions and fallbacks in visual QC reports. Validation across simulations, reference datasets, and clinical data supports its use in research settings while identifying where acquisition and modeling choices limit interpretation.

## Methods

### Pipeline architecture

PMRIPrep is implemented in Python 3.11–3.12 and orchestrated by Nipype [28], which manages inter-stage data dependencies and enables parallel execution. The pipeline is distributed as Docker and Apptainer container images that bundle its runtime dependencies for reproducible execution. PMRIPrep reads input organized according to the Brain Imaging Data Structure (BIDS) [29]. Its outputs follow BIDS-Derivatives organization and include JSON sidecar files for provenance.

Five stages are conditionally executed according to data availability: anatomical preprocessing, tumor segmentation, DSC quantification, DCE quantification, and QC reporting (Fig. 1). Before workflow construction, modality-specific preflight checks validate the acquisition metadata needed for quantitative processing. Errors halt processing, while warnings and remediation guidance are retained in the report.

### Spatial handling and motion

DSC and DCE time series data are motion-corrected within their respective acquisition grids, and quantitative fitting and map generation remain in native perfusion space. The anatomical reference images, atlas, and masks (lesion, tissue, and NAWM) are transformed to the corresponding perfusion space. Categorical masks use nearest-neighbor interpolation, whereas continuous maps use continuous interpolation with explicit validity-mask handling where required. This avoids additional anatomical-or standard-space interpolation of dynamic data and quantitative maps. Measured T1 and B1+ maps are used in DCE processing only when the required registrations pass QC; otherwise PMRIPrep uses the documented fallback. The NAWM-normalized DSC derived cerebral blood volume map (nCBV) is similarly used as a spatially varying plasma volume fraction (v_p_) prior only when the full registration chain passes QC.

Framewise displacement is retained after rigid-body motion correction, and motion parameters are recorded in QC.

#### (1) Anatomical preprocessing

The selected T1w reference is reoriented to RAS standard orientation, field-of-view cropped, and bias-field corrected with N4BiasFieldCorrection [30]. Brain extraction is performed by default with HD-BET [31]. When an eligible precontrast T1w image is available, three-class tissue segmentation (white matter, gray matter, cerebrospinal fluid) is performed using FMRIB’s Automated Segmentation Tool (FAST) [32].

Secondary structural modalities are coregistered to the T1-weighted reference image using a six-degree-of-freedom rigid-body transformation utilizing FMRIB’s Linear Image Registration Tool (FLIRT) [33] with modality-appropriate cost functions.

#### (2) Tumor segmentation

Automated glioma segmentation uses HD-GLIO (a well-validated, deep-learning-based segmentation tool) [27] to generate enhancing-, non-enhancing-, and whole-tumor masks when pre- and post-contrast T1w, T2-weighted (T2w), and T2-weighted fluid-attenuated inversion recovery (T2-FLAIR) images are available; user-supplied masks are also supported. The automatic NAWM reference is derived from eroded supratentorial white matter after tumor exclusion. The NAWM reference is restricted to the contralateral hemisphere for lateralized tumors; for bilateral tumors, it remains unilateral and is selected from the hemisphere with lower tumor volume. Infratentorial tumors use both supratentorial hemispheres. The resulting mask is used for perfusion normalization and same-session DSC-DCE comparison without manual NAWM placement.

#### (3) DSC Quantification

The DSC processing pipeline comprises motion correction using rigid ANTs registration [34,35], conversion of signal to change in transverse relaxation rate (ΔR2*) using the standard logarithmic transformation [1], automated AIF detection, optional per-voxel Boxerman-Schmainda-Weisskoff (BSW) leakage correction [36], AUC-based relative CBV (rCBV) calculation, and finally perfusion estimation using either an SVD-family method or CPI.

##### (3a) Baseline and AIF detection

A configured pre-bolus interval (five frames by default) supplies S0 for an initial ΔR2* conversion. Bolus arrival is then estimated from a weighted consensus of high-percentile and mean whole-brain concentration curves, and this preliminary estimate can be refined using the selected or externally supplied AIF. The baseline interval preceding the final arrival estimate is used for baseline centering. The arrival estimate, its source and its confidence are recorded in QC provenance.

An automatic cascade searches arterial atlas masks [37] transformed to native DSC space before broadening to a whole-brain search region. Candidates are filtered and ranked by bolus timing, amplitude, shape, baseline behavior, and partial-volume indicators; peri-lesional candidates are excluded when a lesion mask is available. The final AIF is an inverse-cost-weighted average of the best candidates from the first acceptable tier. Complete baseline-refinement rules, AIF tier definitions and fallback behaviors are reported in Supplementary Methods. These rigorous criteria were used to minimize user bias and improve reproducibility across varying image qualities and clinical settings.

##### (3b) Leakage correction

BSW leakage correction [36] estimates K1 and K2 independently in each voxel by unconstrained ordinary least-squares regression of the voxel concentration-time curve against the mean concentration-time curve of an eligible whole-brain reference pool and the cumulative integral of that reference curve. The reference pool excludes voxels whose late raw signal exceeds the pre-bolus mean by more than one baseline sample standard deviation, screening persistent T1-dominant enhancement without rectifying the fitted K2 distribution. Correction can be disabled explicitly; when disabled, no K2 derivative is produced and any DSC-informed DCE prior is skipped.

##### (3c) Relative cerebral blood volume (rCBV)

The primary rCBV output is the ratio of the tissue concentration integral (after BSW correction, when enabled) to the baseline-centered AIF integral from bolus arrival through the end of acquisition [36,38]. When no acceptable AIF is detected, AIF-dependent outputs are not computed, and nCBV is obtained directly from the tissue integral, as the AIF integral cancels on NAWM normalization. For singular value decomposition (SVD)-family methods, mean transit time (MTT) is calculated as area under the curve (AUC)-based rCBV divided by deconvolved relative cerebral blood flow (rCBF) according to the central-volume theorem [17,38]; control point interpolation (CPI) instead derives MTT from the fitted residue. Residue-integral rCBV is retained as a diagnostic. Primary rCBV is normalized to its NAWM median to produce dimensionless nCBV [10,38]. rCBF is normalized independently to its NAWM median to produce dimensionless nCBF.

##### (3d) Deconvolution

Four DSC deconvolution methods are implemented. Standard SVD (sSVD) [17] uses singular-value truncation; oscillation-index SVD (oSVD) [18] uses a block-circulant formulation and data-driven truncation; and Tikhonov regularization [19] uses a modified first-difference penalty and L-curve selection over 100 log-spaced regularization parameters [39]. These three methods operate on the tissue concentration-time curve. Non-parametric CPI [20] instead fits the measured DSC magnitude signal using a continuous AIF-residue convolution model. Before inference, all tissue signals are scaled by one factor so that the median pre-bolus signal within the brain mask matches the prior mean for the baseline signal. Fitted signals are then returned to the original signal units. The unit residue is represented by monotone, non-negative piecewise cubic Hermite interpolation through moving control points; the control-point count is fixed for each acquisition and selected automatically by default. Flow, residue shape, AIF-relative delay, and baseline signal are estimated jointly. When BSW leakage handling is enabled, CPI jointly estimates signed K2 using a voxel-specific Gaussian prior centered on the BSW regression estimate [36]. Voxel-independent variational Bayesian inference (VI) [40] is the default inference method. Spatial variational Bayes (SVB), which regularizes flow estimates across neighboring voxels, is available as an alternative [41,42]. Fits that are numerically invalid or fit the data worse than the signal’s temporal mean can be replaced voxelwise by oSVD when fallback is enabled; these substitutions are identified in a fallback mask. Full forward-model, prior, inference, and validation details are reported in Supplementary Methods and Sections A-D.

The default method is oSVD, as it provides computationally efficient, non-parametric deconvolution while reducing sensitivity to AIF-tissue arrival-time differences relative to sSVD [18]. Reported rCBF is derived from the maximum of the recovered flow-scaled residue and is reported on a relative scale. Tmax is the time of the residue maximum for sSVD and Tikhonov, the signed residue-peak delay for oSVD, and the fitted AIF-to-tissue delay for CPI; TTP is the interval from the estimated bolus arrival used to define the baseline to peak tissue concentration.

#### (4) DCE Quantification

##### (4a) B1+ and T1 mapping

B1+ inhomogeneity is estimated from Actual Flip Angle Imaging [43] or a precomputed map when available; otherwise nominal flip angles are used. Native T1 maps are computed from variable-flip-angle (VFA) spoiled gradient-recalled echo (SPGR) acquisitions [44] by weighted linear least-squares (WLLS)-initialized nonlinear fitting of the Ernst equation in signal space, with standalone WLLS also available. Map-level QC records conditioning, B1+ status, non-finite voxels, and population-reference comparisons without replacing a numerically valid measured map. In the absence of a supplied T1 map, a uniform field-strength-dependent T1 is used when VFA data are unavailable, no valid map can be estimated, registration to DCE space is rejected, or fallback is explicitly requested. The VFA and B1+ pathways were evaluated against Quantitative Imaging Biomarkers Alliance (QIBA) T1 and B1-correction digital reference objects [24] (Supplementary Section M).

##### (4b) Signal-to-concentration conversion

DCE signal intensity is converted to gadolinium concentration by inversion of the spoiled gradient-echo signal equation using B1+-corrected flip angles [26,44]. Whole-blood AIF concentration is subsequently converted to plasma concentration used by the Tofts convention with arterial hematocrit (Hct), C_p_ = C_blood_ / (1 - Hct) [2]. The default arterial hematocrit of 0.42 follows the convention used for the Parker population AIF [45]. The volume transfer constant K^trans^ is therefore reported on the plasma-concentration convention used by the model without rescaling.

##### (4c) DCE AIF detection

DCE AIF detection uses an analogous atlas-to-whole-brain cascade, scoring candidates in the concentration domain according to peak concentration, contrast-to-noise ratio (CNR), bolus timing, washout fraction, bolus width, and upstroke slope. A second pass may retain the best arterial-timed single candidate when no multi-voxel pool is accepted, if it passes the same pooled checks. Failed detection falls back to the dose-scaled Parker population AIF [45]. The complete acceptance criteria and fallback behavior are reported in Supplementary Methods.

##### (4d) Pharmacokinetic model selection and fitting

Four pharmacokinetic models are implemented: standard Tofts [46], Extended Tofts [2], Patlak [47], and the two-compartment exchange model (2CXM) [48,49]. Predicted tissue concentration curves are calculated by analytic convolution of each model’s impulse response with a shape-preserving piecewise cubic Hermite interpolant of the plasma AIF [50]. The default estimator uses bounded least squares (BLS) within physiological parameter bounds: linear fitting for Patlak, a search over k_ep_ with linear fitting of the remaining parameters for Tofts and Extended Tofts, and trust-region fitting for 2CXM. Effective delay is estimated by pooling the fit error over candidate delays across in-plane neighbors to give a shared reference delay; a voxel-specific delay replaces it when it lowers BIC by at least 6 (Supplementary Methods). Optional VI, initialized from the retained BLS fit at its selected delay, produces Bayesian parameter estimates and parameter-wise posterior-standard-deviation maps [40]. With automatic model selection, eligible candidates are fitted by BLS and ranked voxelwise by the Bayesian information criterion (BIC) [51]. A model more complex than Patlak is retained only when it improves BIC by at least 6 units relative to a finite Patlak fit, corresponding to the conventional strong-evidence range [52]. While Patlak and Extended Tofts are always eligible, 2CXM eligibility is restricted to temporal resolutions of ≤2 s, motivated by the sampling sensitivity of plasma-flow estimation [53] (Supplementary Section J); 2CXM must also pass additional evidence checks against Extended Tofts (Supplementary Methods).

Standard Tofts remains available by explicit selection but is excluded from the automatic pool as its zero-plasma-volume assumption omits the vascular contribution expected in brain DCE.

#### (5) Cross-modality integration

When same-session DSC and DCE data are processed, PMRIPrep can construct a DCE v_p_ prior from NAWM-normalized DSC CBV using a fixed NAWM plasma-volume reference: μ(v_p_) = 0.010 × nCBV (Supplementary Methods and Section L). The prior is applied only during Bayesian refinement and is represented by a Gaussian distribution in the bounded fitting parameterization, using a nominal natural-space standard deviation of max[0.60 × μ(v_p_), 0.001]. Default least-squares fitting does not use the DSC-derived prior. The prior is skipped when DSC tissue CNR or nCBV support is inadequate, leakage correction is disabled or cannot construct an eligible reference curve, cross-modal registration QC is unacceptable, the whole-brain median of the DSC-derived v_p_ prior falls outside the prespecified screening range (0.005-0.030), or the prior is disabled by configuration.

DSC-DCE concordance is assessed after separate within-subject normalization of v_p_ and CBV by their NAWM medians, followed by matched Gaussian smoothing (σ = 1 mm) over valid voxels. Whole-brain and, when sufficient jointly valid lesion voxels are available, lesion-restricted concordance is reported. Concordance is labeled independent only when none of the retained voxelwise DCE fits used the DSC-informed prior; otherwise, it is labeled prior-informed. Prior-informed agreement is not presented as independent validation.

#### (6) Quality control

PMRIPrep generates a unified per-session HTML report following the visual-reporting approach established by fMRIPrep [14]. It includes brain-mask and registration assessment, tumor overlays, motion summaries, AIF curves, parameter maps, fit and leakage diagnostics, and DSC-DCE concordance when available (Supplementary Fig. S2).

Quantitative maps are not censored by the DCE QC mask. The mask identifies voxels with a retained converged fit, R² greater than 0, and finite parameters satisfying the model-specific checks and identification criteria detailed in Supplementary Methods. Its component rejection counts and thresholds are recorded in the QC JSON sidecar.

When available, it is used to generate explicitly labeled QC-filtered companions for regional statistics and fractional volumes from pharmacokinetic maps. The QC mask is not applied to model-free DCE statistics, which are independent of the retained pharmacokinetic fit.

### Evaluation data

#### DSC deconvolution accuracy

All four deconvolution methods were tested on a common signal-domain reference generated using an independently implemented forward model. An analytic gamma-variate AIF was convolved by adaptive quadrature with exponential, triangular, and plug-flow box residues before point sampling. The main comparison fixed full CBV at 4% and varied CBF from 10 to 70 mL/100 g/min, corresponding to MTT values from 24 to 3.43 s, without tissue delay. It included noiseless, SNR-100, SNR-50, and SNR-20 Rician magnitude-signal conditions with 200 realizations per noisy cell. CPI received the original magnitude signal, while the SVD methods received the corresponding concentration curves from PMRIPrep’s signal conversion. A common AIF/tissue scale set the exponential-residue, 4%-CBV, 60-mL/100-g/min reference to a 40% peak tissue signal drop, leaving the true perfusion values unchanged. A separate fixed-flow stress matrix crossed five residue families, MTT values of 5 and 16 s, and delays of 0 and 3 s under the same calibrated observation and noise conditions. Separate generators using the same calibration tested signed leakage, spatial inference, and transport dispersion.

#### External DSC digital reference object

External DSC validation used the GBM-DSC-MRI digital reference object (DRO) from The Cancer Imaging Archive [22,23] at 1.5 T and 3 T, comparing NAWM-normalized AUC-CBV (nCBV) against known ground truth across no-leakage, uncorrected-leakage, and BSW-corrected-leakage conditions.

#### DCE pharmacokinetic model accuracy

Algorithm-level accuracy was evaluated for Patlak, Tofts, Extended Tofts, and 2CXM on controlled concentration-time curves across 0–10% noise using the default BLS estimator with effective-delay estimation. The QIBA DCE-MRI DROs [24] separately evaluated Standard-Tofts recovery under normal and reduced cardiac output, Extended-Tofts comparability, reduced-cardiac-output sensitivity, and VFA T1/B1+ mapping (Supplementary Sections F, G and M).

#### DSC-informed DCE prior

Internally generated paired-modality experiments compared least-squares fitting, VI with each model’s default priors and VI with the DSC-informed prior across 20 paired random seeds. Concentration-level experiments used model-specific Extended-Tofts, Patlak and 2CXM truth, temporal resolutions of 2, 4 and 7.5 s, and 5% and 10% noise. Prior centers in these experiments were set from true v_p_, with or without controlled random and systematic errors. Signal-level DSC recovery was tested with Extended-Tofts DCE at 4-s and explicit 2CXM at 2-s temporal resolution, both with 5% noise, and NAWM calibration with Extended Tofts. Calibration included spatially matched prior means and randomly reassigned values. Both arms used the known AIF and shared the BLS model, selected delay and starting estimates. Simulated prior effects were calculated only from voxels with successful VI fits in both arms; fallback counts were reported separately. Numerical generation and fitting details are given in Supplementary Section L.

#### DCE synthetic phantoms

End-to-end pipeline validation was performed on the International Society for Magnetic Resonance in Medicine (ISMRM) Open Science Initiative for Perfusion Imaging (OSIPI) DCE digital reference objects [13]: 2 synthetic subjects with 2 sessions each, processed from BIDS input through quantitative parameter maps using BIC-based automatic model selection and automatic AIF detection.

#### OSIPI clinical test-retest cohort

Reproducibility was assessed using the OSIPI clinical DCE test-retest dataset [13]: eight subjects with recurrent glioblastoma selected from the RIDER Neuro MRI collection and scanned twice on the same 1.5 T scanner, 2 days apart. Each session was processed with default settings; all parameters were determined automatically. Following the OSIPI-DCE challenge methodology, mean K^trans^ was computed within the supplied tumor masks for each visit.

#### External-site clinical cohort

Eleven post-treatment glioblastoma cases were recruited. The cohort was not part of the initial development dataset and was used to demonstrate the complete workflow on external-site clinical data, with no prospective sample-size calculation or randomized allocation. This study was approved by the institutional review board of the University of Pennsylvania (protocol # 829645; date of approval 7 June 2023) and was compliant with the Health Insurance Portability and Accountability Act.

The MRI protocol was acquired on a Siemens MAGNETOM Vida 3 T scanner. The imaging protocol included a DSC sequence (repetition time/echo time (TR/TE) = 1500/35 ms, flip angle 60°, 120 volumes) and a DCE sequence (3D SPGR, TR/TE = 3.23/1.12 ms in nine cases and 5.09/1.57 ms in two, flip angle 21–25°, temporal resolution 7.1–7.5 s, 45 volumes). As variable flip angle (VFA) or B1+ data were not available, DCE modeling used a uniform T1 of 1200 ms.

In addition to the default least-squares analysis, VI was applied to the BLS fits with the DSC-informed v_p_ prior either enabled or skipped, using the same model, delay and starting estimates in both arms. The two VI analyses used identical DCE and DSC preprocessing outputs and differed only in whether the DSC-informed prior replaced the model’s standard v_p_ prior during Bayesian refinement. A separate case-level comparison used an enhancing-tumor mask to compare PMRIPrep maps with Siemens Spectra.

### Computational reproducibility

All analyses used PMRIPrep 1.0.0rc6.

### Statistical analysis

Simulation performance was summarized as signed bias or absolute error relative to known ground truth, with positive signed bias denoting overestimation. Repeated simulations were summarized using medians with interquartile ranges or 95th percentiles across realizations, as stated for each result. OSIPI accuracy and repeatability scores followed the published challenge definitions [13], using all finite tumor-ROI voxels; QC-filtered analyses were secondary. Repeatability was assessed using within-subject coefficients of variation, repeatability coefficients and Bland-Altman analysis. The primary calculations used the population standard deviation of each pair, as in the challenge; results using the sample standard deviation are reported in Supplementary Section I. Confidence intervals were estimated by bootstrap resampling of subjects for repeatability and paired simulation replicates for the prior experiments.

Pearson and Spearman correlations summarized association, and Lin’s concordance correlation coefficient summarized cross-software agreement. Paired clinical prior comparisons were summarized within subjects and then across subjects using the median and range. Simulated prior effects were summarized as mean paired percentage changes in RMSE. Clinical analyses were descriptive, and voxelwise correlations were not treated as independent subject-level observations. Statistical definitions and analysis-specific details are provided in Supplementary Methods and Sections I and L.

## Data availability

The OSIPI-DCE challenge datasets used for synthetic and test-retest evaluation are available from the ISMRM OSIPI consortium [13]. The GBM-DSC-MRI-DRO used for external DSC validation is available from The Cancer Imaging Archive [23]. The QIBA DCE-MRI digital reference objects are available from Zenodo [24]. Internally generated digital reference objects and their executable tests are distributed with the source code. The external-site clinical data that support the findings of this study are available from the corresponding author upon reasonable request. These data are not publicly available due to privacy and ethical restrictions.

## Code availability

PMRIPrep is freely available under the Apache License, Version 2.0. Source code, documentation, and containerized images are available at https://github.com/arulseh/pmriprep.

## Acknowledgements

Supported by Ministry of Health, Czech Republic - conceptual development of research organization (FNMH, 00064203), IG204302, IG244301, IG244302.

