## Supplementary_Information for "PMRIPrep: automated preprocessing and quantification of DSC and DCE MRI"

### Supplementary Materials

This supplement provides additional methods for PMRIPrep's anatomical preprocessing, tumor segmentation, arterial input function and bolus-arrival detection, dynamic susceptibility contrast (DSC) and dynamic contrast-enhanced (DCE) analysis, cross-modal integration, quality control, and derivative generation. Validation results cover DSC deconvolution and leakage correction, DCE pharmacokinetic recovery, OSIPI accuracy and repeatability, model selection, effective-delay estimation, T1 and B1+ estimation, the DSC-informed DCE prior, and a clinical cross-software comparison. PMRIPrep analyses and outputs were generated using version 1.0.0rc6.

#### Supplementary Methods

##### Anatomical preprocessing

PMRIPrep requires at least one precontrast or postcontrast T1w image. When both are available, the anatomical reference is selected according to contrast state, spatial resolution, and voxel anisotropy, with preference for precontrast T1w. If only one T1w image is available, it is used. Postcontrast-only processing is supported but does not include tissue segmentation.

The selected T1w image is reoriented to RAS and field-of-view cropped. Optional denoising (off by default) uses ANTs DenoiseImage with a Rician noise model [1]. Next, intensity nonuniformity is corrected using N4BiasFieldCorrection [2]. Axes with submillimeter voxel spacing are resampled to 1 mm using cubic interpolation, while voxel dimensions ≥1 mm are retained. Brain extraction is performed on the corrected reference image with HD-BET by default [3]. ANTs brain extraction using the TemplateFlow OASIS30ANTs template and brain-probability mask is available as an alternative and is used when HD-BET is unavailable.

Three-class white matter, gray matter, and cerebrospinal fluid segmentation is performed on precontrast T1w data using FSL FAST [4]. When precontrast T1w is the anatomical reference, it is segmented directly after preprocessing and brain extraction. When postcontrast T1w is selected as the reference, the precontrast image independently undergoes reorientation, cropping, optional denoising, and bias-field correction; it is then rigidly registered to the postcontrast reference, masked using the reference-space brain mask, and used for segmentation if its geometric-mean voxel size is ≤2 mm and its voxel-anisotropy ratio is ≤2.5.

Secondary T1w, T2w, and FLAIR images are processed only when required for tumor segmentation, selected as the registration reference, requested as a perfusion-space underlay, or needed to construct a requested T1w subtraction image. These images undergo reorientation, optional denoising, and bias-field correction and are rigidly registered to the anatomical reference using FSL FLIRT [5]. Normalized mutual information is used for postcontrast T1w registration and normalized correlation for T2w and FLAIR. The anatomical brain mask is then applied to secondary images in reference space. The transforms and brain mask are also applied to reoriented images that have not undergone bias correction or intensity normalization, producing reference-space inputs for HD-GLIO (when applicable). When T1w subtraction is requested, both acquisitions are available, and the metadata checks identify no acquisition mismatch, the bias-corrected precontrast image is subtracted from the bias-corrected postcontrast image in the selected anatomical reference space, and the brain mask is applied.

The brain-extracted anatomical reference is also rigidly aligned to the TemplateFlow MNI152NLin6Asym template using FSL FLIRT with normalized mutual information. This provides the transform needed to transfer atlases into native anatomical and perfusion spaces.

#### Tumor segmentation and NAWM reference construction

When precontrast and postcontrast T1w, T2w, and FLAIR images are available, PMRIPrep by default uses HD-GLIO [6] to generate enhancing-tumor and non-enhancing-tumor masks; these masks are also combined into a whole-tumor mask. User-supplied lesion and NAWM masks are also supported. An automatic NAWM mask is created from supratentorial voxels with a FAST white-matter probability of at least 0.90. Tumor compartment is determined from whole-tumor overlap with transformed supratentorial and cerebellar atlas masks. For supratentorial tumors, laterality is measured relative to the native-T1w brain-symmetry plane: tumors meeting the 60% laterality criterion use the contralateral hemisphere, while bilateral tumors use the hemisphere with lower tumor burden. Infratentorial tumors use both supratentorial hemispheres. If the brain-symmetry plane cannot be estimated, world $x = 0$ is used with a QC warning; if the transformed supratentorial atlas is unavailable, the candidate mask is retained for QC but withheld from normalization. Voxels within 20 mm of the whole-tumor mask are excluded, and the remaining mask is eroded by a physical distance corresponding to two times the smallest voxel dimension. If fewer than 500 voxels remain, the erosion is reduced stepwise. A NAWM mask that remains below 500 voxels is retained but labeled invalid in QC.

#### DSC and DCE preprocessing and motion handling

DSC and DCE data are reoriented to RAS and, by default, motion-corrected by rigid ANTs registration to the baseline mean, using mutual information and linear resampling [7,8]. Framewise displacement is calculated from the estimated motion parameters (sum of absolute frame-to-frame changes, with rotations converted to displacement at a 50-mm radius) and recorded in QC. Native perfusion-space brain masks are generated from a pre-bolus DSC mean or early precontrast DCE mean using HD-BET by default. The anatomical reference is rigidly registered to each native perfusion reference with six-degree-of-freedom FSL FLIRT. DCE registration uses normalized mutual information and retains header alignment unless a fitted transform improves foreground-overlap Dice by more than 0.01. For DSC, the brain-extracted T1w intensity histogram is first matched to the DSC reference; normalized correlation is then used for a dual-path coarse search and normalized mutual information for refinement. Alternative initializations are evaluated using foreground-overlap Dice. Tissue, tumor, and NAWM masks originating in anatomical space and arterial atlas masks originating in standard space are then transformed to the corresponding perfusion space. Dynamic data and quantitative maps remain in their native acquisition spaces.

#### Arterial input function (AIF) detection algorithm

Arterial input function specification is a central step in quantitative DSC and DCE analysis [9]. PMRIPrep automates AIF detection for both modalities using a multi-level cascade that adapts its search space to the available data. The default workflow requires no manual AIF selection; an external AIF can instead be supplied explicitly. A native-space search mask can also be supplied; it replaces the atlas tiers and bounds every candidate search, without a whole-brain fallback.

The cascade proceeds through three levels of decreasing spatial specificity:

1. **Atlas-constrained search:** candidate voxels restricted to arterial territory masks registered to native space, with progressively relaxed physiological criteria.
2. **Whole-brain search:** if atlas-based detection fails, the search expands to the entire brain with relaxed criteria.
3. **Fallback:** if no voxel-based AIF can be found, DCE uses the Parker population AIF, and DSC produces only AIF-independent outputs.

##### *Arterial atlas construction*

The atlas-constrained search uses arterial territory masks derived from the Montreal Neurological Institute (MNI)-space Vessel Occurrence Probability Atlas [10], a population-averaged time-of-flight magnetic resonance angiography (MRA) dataset.

Source data and thresholding

The vesselProbabilities.nii volume from the atlas (0.5 mm isotropic, natively in MNI space) was thresholded at 20% vessel occurrence probability and binarized. This empirical cutoff was selected by visual inspection to retain recognizable vascular structures while limiting nonvascular inclusion. The resulting binary mask was manually edited to remove venous and other non-arterial structures, retaining only the major arterial territories.

Dilation, resampling, and mask hierarchy

The cleaned binary mask was morphologically processed to create smooth, dilated territorial coverage. The processed masks were resampled from 0.5 mm to a 2 mm MNI152 grid (91 × 109 × 91). The resulting tiered cascade provides increasing spatial coverage. In the mask identifiers below, MCA and ACA denote the middle and anterior cerebral artery territories, respectively:

1. **MCA_ACA:** the base arterial mask with contiguous territorial coverage.
2. **MCA_ACA_dil:** one additional dilation, accommodating registration imprecision between MNI and native space.
3. **full_mask:** the full MRA arterial mask including the basilar artery and posterior circulation.
4. **full_mask_dil:** an additionally dilated full-arterial mask, used only when the expected atlas files are unavailable.

DCE searches MCA_ACA, MCA_ACA_dil and full_mask in that order; DSC searches MCA_ACA_dil and full_mask.

The bundled masks are stored in RAS orientation on the TemplateFlow MNI152NLin6Asym resolution-2 grid. At runtime, the corresponding transform chain maps them into native perfusion space.

*Common detection steps*

Search mask preparation

Before candidate search, a spatial mask is constructed to exclude regions likely to contain venous signal, partial-volume contamination, or lesion-affected tissue:

1. **Brain-edge exclusion.** To reduce superficial-vessel and boundary partial-volume contamination, DSC retains voxels at least 8 mm from the slice-wise, hole-filled outer brain boundary. This in-plane criterion is applied to all DSC search tiers; if it empties a tier, the original tier region is restored. DCE atlas-constrained tiers use two iterations of 3D binary brain-mask erosion, reverting to the uneroded mask if fewer than 100 voxels remain. The DCE whole-brain

fallback instead retains voxels at least 12 mm from the brain boundary and similarly omits this depth restriction if fewer than 100 voxels remain.

2. **Venous-sinus exclusion.** Whole-brain DSC and DCE search excludes a posterior midline band defined from the brain in-plane symmetry axis. The band covers the central 30% and posterior 60% of the brain, excluding deep midline veins while preserving anterior midline arteries. If the symmetry axis cannot be estimated, DSC aligns the same band with the image axes, and DCE excludes a strip covering the central 20% of the image width along its full anterior–posterior extent. This exclusion is omitted for atlas-constrained tiers and user-supplied search masks.

3. **Slab edge exclusion (DCE).** Voxels within 15 mm of the superior slab boundary are excluded in whole-brain search and within 4 mm in atlas-constrained search. To reject inflow artifacts from unsaturated spins entering the excitation volume, the inferior exclusion is the larger of 20 mm and four slice thicknesses, capped at 35% of the slab depth.

4. **Edge slice exclusion (DSC).** The first and last axial slices are excluded.

5. **Lesion exclusion.** If a lesion mask is provided, lesion voxels are removed from the search mask. A peri-lesional buffer (15 mm by default, configurable with a 10-mm minimum) further excludes tissue where tumor-related displacement, compression, or angiogenesis may disrupt the vasculature.

<u>Tissue time-to-peak reference</u>

A tissue time-to-peak (TTP) reference is computed to cross-validate candidate timing. Parenchymal voxels are identified as those with peak concentration in the 25th-75th percentile range, favoring parenchymal tissue over low- and high-enhancement extremes. The median TTP of these voxels serves as a tissue reference. Arterial candidates should peak before tissue; candidates peaking substantially after tissue are penalized or rejected as likely venous.

<u>Baseline detection</u>

Both modality-specific pathways below use a shared four-phase bolus-arrival detector. First, max-slope backtracking and second-derivative peak detection identify a derivative-based onset anchor; when the temporal resolution ($\Delta t$, the time between the starts of successive dynamic volumes; BIDS RepetitionTime) is 4 s or coarser, only the max-slope method is used. Second, amplitude z-score, gradient z-score, and cumulative-sum tests can move the estimate earlier only when at least two agree within 1–3 frames, the baseline window contains at least eight samples, and the signal change exceeds 1.5%; two or more tests lagging the anchor by 1–3 frames instead confirm it and raise confidence. Third, a weighted median combines estimates when additional curves are available. Finally, confidence is classified as high, medium, or low from method agreement and the strength of the derivative onset. DSC and DCE differ in the curves supplied to this detector and in how its estimate is reconciled with configured baseline settings and subsequent arterial information.

For DSC, the configured pre-bolus interval (five frames by default) is used only to bootstrap motion correction and the initial signal-to-concentration conversion. A preliminary arrival estimate is then obtained within the brain mask from the 99th-percentile concentration curve, with the 90th-percentile and mean curves contributing to the weighted consensus. This ensemble is used when more than 100 brain voxels are available; otherwise the bootstrap value is retained. The preliminary estimate is constrained to be no earlier than frame $\max(5, \lfloor 10\ \mathrm{s}/\Delta t \rfloor)$, and frames from the beginning of the series through the sample immediately before the estimated arrival define the baseline. After a measured AIF has been selected, or when an external AIF is supplied, the detector is applied again to the nonnegative part of that arterial curve with a two-frame minimum. This arterial estimate replaces the preliminary estimate when the preliminary estimate did not exceed the configured pre-bolus interval and the arterial estimate does, or when the arterial estimate has high confidence and differs by more than five frames; estimates outside 2 through half the series length are rejected. Otherwise the preliminary estimate is retained. The final baseline frame count, the preliminary and arterial estimates, the selected source, and detection confidence are recorded in QC provenance. When no AIF is accepted, the preliminary estimate is retained.

For DCE, automatic detection uses the 95th-percentile raw-signal curve within the brain mask when more than 100 voxels are available, and the mean curve for smaller masks or the mean of nonzero voxels when no mask is available. Automatic arrival detection imposes a minimum of five baseline frames. The detected arrival frame defines the number of precontrast baseline frames. When a baseline-frame count of at least five is configured, it defines the S0 window, but bolus arrival is still estimated for timing and QC. The configured window is shortened if necessary so that at least one subsequent volume remains and it ends before the detected arrival. If detection, configuration, or acquisition length provides fewer than the required five baseline frames, up to the first five volumes are used as the baseline, limited to all but the final volume and not extending beyond the retained bolus frame. The original detected arrival and fallback reason are recorded, baseline QC is marked poor, and a warning states that the resulting S0 estimate and quantitative pharmacokinetic maps are unreliable.

##### Hemisphere and tumor laterality

Both modalities record the left–right coverage of the selected AIF voxels relative to the brain-symmetry plane, or world x = 0 when the plane cannot be estimated. When a lesion mask is available, tumor laterality is classified relative to the same plane, with the enhancing-tumor mask preferred when available. Contralateral and ipsilateral candidate counts are reported for unilateral tumors. Lesion and peri-lesional exclusion is applied independently of laterality.

##### Candidate averaging and smoothing

Each selected DSC or DCE candidate contributes its voxel concentration curve to the pooled AIF. When multiple candidates are selected, DSC uses weights proportional to $1/[c_i - \min_j(c_j) + 0.1]$, normalized to sum to one, where $c_i$ is the candidate cost, and DCE uses weights proportional to candidate scores. If the DSC multi-candidate pools are rejected, the single-best fallback evaluates the lowest-cost voxel.

DSC applies second-order Savitzky-Golay smoothing with a Δt-adaptive odd window targeting approximately 7.5 s when the curve contains at least five samples. DCE uses the weighted measured curve at the acquired temporal resolution. Negative samples are retained in measured AIFs to avoid biasing the integral; DCE quality-control (QC) provenance reports their number, fraction, and minimum both overall and after the peak.

A gamma-variate model [11] is fitted to the first pass of the selected DSC AIF and scaled to its first-pass area. The fitted curve and its goodness of fit are reported for QC only; the measured AIF is used for deconvolution and as the denominator of the AUC-based rCBV.

##### Averaged-pool acceptance

After averaging, DSC candidate pools are checked for a dominant, correctly timed first-pass peak (global maximum within 40 s of baseline, with no later value above the first-pass peak), agreement with gamma-variate peak behavior [11] (fitted peak within 12% of the measured peak), and adequate return toward baseline (baseline-return ratio of at least −3%, when calculable). DCE pools are checked after plasma conversion and amplitude calibration. During Pass 1, a DCE pool is rejected when its late-tail-to-peak ratio exceeds 0.35, its full width at half maximum (FWHM) exceeds 22 s, its late tail has more than two secondary maxima, its concentration 30 s after the peak (at least four frames) exceeds 50% of the peak, its peak exceeds three times (five times at permissive stringency) the dose-scaled expected maximum, or calibration would require more than threefold upward scaling. Up to eightfold scaling is accepted only in the base MCA/ACA mask, when the shape checks pass and the pool contains at least ten candidates across at least two slices; the resulting AIF is labeled reduced confidence.

Failed pools are trimmed as described for each modality below. Per-tier decisions and pool sizes are recorded in QC provenance.

*DSC-specific detection*

Candidate identification

Candidates are identified by percentile thresholding on peak concentration (ΔR2*): all voxels above the 90th percentile of peak concentration within the search mask are retained.

Candidate metrics

The first-pass concentration peak is located in a nominal 3–40 s post-baseline window. Eleven candidate metrics combine that peak with baseline, post-peak and tail summaries:

| Metric | Definition |
|---|---|
| TTP | Time to peak within first-pass window (s) |
| BRR | Baseline return ratio: tail/peak × 100 (%) |
| PBR | Peak-to-baseline ratio: peak/baseline |
| SPR | Secondary peak ratio: max post-first-pass/peak × 100 (%) |
| PMI | Post-peak monotonicity: fraction of decreasing points in 30 s post-peak (%) |
| FWHM | Full width at half maximum (s) |
| PH | Peak height from signal: 100 × (S0 - $S_{min}$) / S0 (%) |
| PSR | Percent signal recovery: 100 × ($S_{tail}$ - $S_{min}$) / (S0 - $S_{min}$) (%) |
| S0 | Mean baseline signal intensity |
| Peak ΔR2* | Peak concentration in ΔR2* units ($s^{-1}$) |
| Tail fraction (late) | Mean ΔR2* over the last 30 s / first-pass peak |

Cost-function scoring

Candidates are scored by a composite cost function combining hard rejection criteria and continuous scoring. Three nested strictness levels (strict, relaxed, permissive) define progressively looser thresholds:

**Hard rejection** (any of the following eliminates a candidate):

- TTP outside the predefined acceptance window (strict: 5–40 s post-baseline, relaxed: 3–40 s, permissive: 2–45 s)
- BRR outside level range (strict: 5–30%, relaxed: 0–40%, permissive: −10–50%)
- PBR below level threshold (strict: 3.0, relaxed/permissive: 2.0)
- SPR exceeding level threshold (strict: 70%, relaxed: 85%, permissive: 100%)

- PMI below level threshold (strict: 60%, relaxed: 55%, permissive: 40%)
- FWHM outside range: maximum (strict: 10 s, relaxed: 12 s, permissive: 18 s); minimum (strict: 3.0 s, relaxed: 2.5 s, permissive: 2.0 s)
- S0 above ceiling ratio of median brain S0 (strict: 0.40, relaxed: 0.60, permissive: 0.80)
- PH below level threshold (strict: 25%, relaxed: 15%, permissive: 12%)
- PSR outside range: ceiling (strict: 110%, relaxed: 120%, permissive: 130%) and floor (strict: 40%, relaxed: 30%, permissive: 20%)
- Peak ΔR2* outside artifact bounds, scaled with field strength and effective Gd dose (3 T full-dose floor: 5 $s^{-1}$, clamped to a 3 $s^{-1}$ noise floor; 3 T full-dose ceiling: 120 $s^{-1}$)
- TTP beyond 80% of the scan duration (all levels)
- S0 below 0.10 of median brain S0 (all levels; heuristic low-baseline-intensity exclusion)

**Continuous scoring** (lower cost = better candidate): The continuous score is a weighted sum of TTP (1.0), PH (0.5), BRR (2.0), PSR (0.3), PMI (0.5), and peak ΔR2* (3.0, dominant weight; log-scale centered at 10 $s^{-1}$, capped at 30 $s^{-1}$ at or below the standard effective dose and scaled upward for larger effective doses). Peak ΔR2* provides a first-pass susceptibility signal related to intravascular contrast concentration and is less dependent on the arbitrary signal-intensity scale than baseline S0. S0 is used only for hard rejection (partial-volume gating), not continuous scoring.

<u>Tissue TTP cross-validation</u>

After scoring, each candidate's TTP is compared to the tissue reference: candidates peaking more than 2 s after tissue TTP are rejected as venous; those peaking 0–2 s after tissue take a large cost penalty, and those peaking within 2 s before tissue take a small one. During whole-brain tiers, candidates falling outside the arterial atlas take an additional penalty. These thresholds are expressed in seconds and applied to the sampled peak times.

<u>Post-scoring kinetic filters</u>

Two kinetic-coherence filters are applied sequentially to remove outlier candidates before averaging:

**Peak-spread gate.** The gate computes the 20th-percentile peak time across the candidate pool and rejects candidates whose peak time exceeds this reference by more than $\lceil 3 \text{ s} / \Delta t \rceil$ frames (3 s at Δt = 1.5 s, 4 s at Δt = 2 s).

**Tail/peak ratio gate.** Candidates with a late tail fraction (mean ΔR2* over the last 30 seconds divided by first-pass peak) exceeding 15% are rejected.

<u>Cascade architecture</u>

The DSC cascade comprises up to six sequential search tiers, followed by two detected-AIF fallbacks:

**Phase 1: Sequential search tiers.** Detection proceeds through up to six search levels of decreasing stringency, stopping at the first level that passes an acceptance gate:

| Level | Search Region | Criteria | Candidates retained* |
|---|---|---|---|
| Atlas strict | Dilated MCA/ACA atlas (MCA_ACA_dil) | Strict | 100% |
| Atlas relaxed | Dilated MCA/ACA atlas (MCA_ACA_dil) | Relaxed | 60% |
| Atlas permissive | Dilated MCA/ACA atlas (MCA_ACA_dil) | Permissive | 40% |
| Full atlas permissive | Full MRA atlas (incl. posterior circulation) | Permissive | 40% |
| Brain relaxed | Whole brain | Relaxed | 60% |
| Brain permissive | Whole brain | Permissive | 40% |

*Fraction of the requested number of candidates.

At each tier, the top-scoring candidates are tested using the averaged-pool criteria described above. A rejected pool is re-evaluated after removing the candidate with the lowest baseline-return ratio, or the highest-cost candidate when that ratio is unavailable, until the pool passes or fewer than three candidates remain.

**Phase 2: Post-cascade fallbacks.** If all six search tiers fail the acceptance gate, two intermediate fallbacks are attempted: (i) an accumulated union of kinetic-filter survivors across all search tiers, and (ii) a single-best fallback selecting the lowest-cost voxel from the union pool. Cross-tier mixing can produce a viable averaged AIF when no single tier's candidates are sufficient.

**Phase 3: No accepted AIF.** If these fallbacks also fail, deconvolution is not performed. rCBF, MTT, Tmax, residue-integral rCBV and AUC-based rCBV are not produced; nCBV, PSR, TTP, CNR, temporal SNR (tSNR) and, when BSW correction is applied, K2 remain available. QC records the detection failure.

When a search mask is supplied, it replaces the atlas tiers; the full-atlas and whole-brain tiers are not used.

*DCE-specific detection*

<u>Candidate signal characteristics</u>

For each voxel in the search mask, nine characteristics are computed:

| Metric | Definition |
|---|---|
| Peak value | Maximum concentration post-baseline (mM) |
| Peak time | Time to peak (s) |
| Peak frame | Frame index of peak |
| Baseline | Mean concentration over first N frames |
| Enhancement | Peak minus baseline (mM) |
| Washout fraction | (peak - tail) / (peak - baseline), clipped [0, 1] |
| FWHM | Full width at half maximum (s), with sub-frame interpolation |
| FWHM (frames) | FWHM in frame units, computed from FWHM (s) / Δt |
| Upstroke slope | Maximum dC/dt during wash-in (mM/s) |

Candidate filters

Within each search region, candidate voxels are selected by a sequence of seven filters of decreasing stringency. Filter 1 is applied first, and later filters are used only while too few candidates pass (fewer than the requested number for filters 1–5, fewer than ten for filters 6 and 7):

| Filter | Criteria |
|---|---|
| 1 | High peak, arterial timing, rapid uptake, washout > 20%, narrow FWHM, steep upstroke |
| 2 | High peak, early arrival, rapid uptake, washout > 20%, narrow FWHM |
| 3 | High peak, early arrival, washout > 20%, narrow FWHM |
| 4 | High peak, early arrival, washout > 10%, narrow FWHM |
| 5 | Moderate peak, early arrival, washout > 10%, FWHM < 45 s |
| 6 | Moderate peak, peak before tissue, narrow FWHM |
| 7 | Moderate peak, peak before tissue |

High and moderate peaks exceed the 98th and 95th percentiles of peak concentration in the search region. Arterial timing requires the voxel to peak more than round(4 s/Δt) frames (at least one) before the tissue TTP reference; early arrival requires the peak within the first 40% of the acquisition. Rapid uptake is enhancement above the 90th percentile, and steep upstroke is a maximum upslope in the top quartile of high-peak voxels. Narrow FWHM is below 25 s, with a minimum threshold of two frames. At Δt > 5 s, early arrival replaces arterial timing in filter 1, the peak-before-tissue requirement is dropped from filters 6 and 7, and candidate scoring (below) shifts weight from timing toward FWHM and washout.

Individual candidate scoring

Candidates passing the filters are scored by a composite quality metric:

- **Peak concentration** (20.0% at Δt ≤ 5 s; 19.0% at Δt > 5 s): deviation from the expected dose-adjusted range
- **CNR** (15.0% at Δt ≤ 5 s; 14.3% at Δt > 5 s): contrast-to-noise ratio (threshold: 5.0 strict to 2.0 permissive)
- **Washout** (20.0% at Δt ≤ 5 s; 23.8% at Δt > 5 s): an asymmetric band centered on a predefined washout target (~78%), with a 0.13 lower half-width and 0.20 upper half-width
- **FWHM** (15.0% at Δt ≤ 5 s; 23.8% at Δt > 5 s): predefined target ~12 s
- **Upstroke slope** (15.0% at Δt ≤ 5 s; 14.3% at Δt > 5 s): steeper slopes receive higher scores
- **Timing** (15.0% at Δt ≤ 5 s; 4.8% at Δt > 5 s): earlier arrival receives a higher score

Cluster-based selection

For the whole-brain fallback, spatial clustering (26-connectivity labeling) identifies coherent arterial groups when more candidates than requested are available. Each cluster is scored by FWHM (30%), washout (30%), upstroke (15%), and size (25%), and the highest-scoring cluster is retained. Atlas-constrained tiers skip this clustering step, as the arterial atlas already supplies the spatial prior and a single connected-component requirement could discard valid arterial candidates in separate territories.

Cascade orchestration

Three ordered arterial masks and three physiological stringency levels are orchestrated with one whole-brain fallback via a two-pass strategy:

**Pass 1: Full cascade.** The cascade can attempt up to 10 tiers (3 arterial masks × 3 stringency levels, then whole-brain relaxed). A tier proceeds only when at least two candidates have score ≥ 0.5 and at least one peaks within 10 seconds of bolus arrival. The weighted, plasma-converted, amplitude-calibrated pool is then checked with the averaged-pool criteria described above. When a pool fails, the candidate contributing most to the failed check is removed and the pool is retested while at least two candidates remain. A pool that needs more than threefold scaling outside the base MCA/ACA mask, or lacks the spatial support required for higher scaling, ends that tier without further trimming. The first accepted tier ends the cascade. The emitted AIF averages up to the requested number of qualifying candidates (ten by default); if this subset differs from the accepted pool, it is checked once more. If it fails this check, the AIF is retained with reduced confidence and increased reported parameter uncertainty; the cascade is not restarted.

**Pass 2: Single-candidate fallback.** If no tier is accepted, the highest-scoring single candidate with arterial timing retained during Pass 1 is checked with the same averaged-pool criteria and, if accepted, used with reduced confidence.

**Population fallback.** If both passes fail, the Parker population AIF [12] is used. Cascade-exhaustion provenance is recorded in the JSON sidecar.

Venous filtering

Candidates are flagged as venous when they peak less than round(4 s/$\Delta t$) frames (at least one) before the tissue TTP reference at $\Delta t \leq 5$ s, or more than one frame after it at $\Delta t > 5$ s. Flagged candidates are removed whenever at least one non-venous candidate survives; if every candidate is flagged venous, the tier fails and the cascade continues.

Candidate quality threshold

Candidates that pass the filters but fail a scoring threshold, such as the CNR minimum, can still receive a lower fallback score when their peak lies within the expected range. Candidates whose composite quality score falls below 0.5 are excluded. At least two qualifying candidates form a detected pool; a single arterial-timed survivor is retained as a possible second-pass fallback, while a tier with no qualifying survivor is exhausted and the cascade continues.

Amplitude calibration

The detected AIF peak is calibrated with a one-sided lower bound by default. The bound is a whole-blood peak of 5.0 mM scaled by the administered gadolinium dose relative to 0.1 mmol/kg and converted to plasma. A curve with a lower peak is scaled up to the bound; curves at or above the bound are left unchanged. An explicit user-specified target peak replaces the automatic bound. Automatic peak calibration is not applied to population or external AIFs. The applied factor, required factor, and calibration mode are recorded as QC provenance.

Parker population AIF fallback

The Parker population AIF [12] consists of two Gaussian peaks (first pass and recirculation) plus a modulated exponential tail:

$$C_b(t) = \sum_i \left[A_i / (\sigma_i\sqrt{2\pi})\right] \exp[-(t - T_i)^2 / (2\sigma_i^2)] + \alpha \exp(-\beta t) / \{1 + \exp[-s(t - \tau)]\}.$$

The population AIF is positioned relative to the end of the detected precontrast baseline and scaled by the administered gadolinium dose relative to 0.1 mmol/kg. As concentration is represented in mM, no relaxivity scaling is applied.

Blood-to-plasma correction

The detected AIF represents whole-blood concentration. For pharmacokinetic modeling, it is divided by (1 - Hct), where Hct = 0.42 by default and is configurable. The Parker population AIF is also parameterized as whole-blood concentration, C_b(t), and receives the same conversion [12]. Both paths therefore provide plasma concentration, $C_p(t) = C\_b(t)/(1 - Hct)$, to the pharmacokinetic models [13]. External AIFs must be supplied as plasma concentration in mM and are not corrected again.

#### Signal conversion

*DSC-MRI*

Raw DSC signal intensity is converted to the change in transverse relaxation rate ($\Delta R2^*$) using:

$$\Delta R2^*(t) = -\ln[S(t) / S0] / TE,$$

where S0 is the mean signal over the configured pre-bolus interval and TE is the echo time read from the Brain Imaging Data Structure (BIDS) sidecar. Curves are later re-centered on the detected baseline, and CPI estimates S0 as a model parameter. The resulting $\Delta R2^*$ is expressed in $s^{-1}$ and is proportional to gadolinium concentration under the assumed linear relationship between concentration and susceptibility-induced relaxation-rate change [14]. Its scale nevertheless depends on acquisition and tissue factors, so it is not treated as an absolute concentration measurement. The signal ratio is lower-bounded to prevent log(0) but not upper-bounded: $S(t)/S0 > 1$ produces negative $\Delta R2^*$, preserving symmetric noise in post-bolus frames rather than introducing positive bias through half-rectification.

*DCE-MRI*

DCE signal intensity is converted to gadolinium concentration by inversion of the spoiled gradient-recalled echo (SPGR) signal equation [13,15,16]. For each voxel and timepoint: (1) baseline signal S0 is computed from the precontrast frames; (2) the longitudinal relaxation rate R1(t) is recovered from the signal ratio S(t)/S0 using the precontrast T1 map, the B1+-corrected flip angle, and the RF-excitation repetition time TR (BIDS RepetitionTimeExcitation); (3) tissue concentration is computed as

$$C_t(t) = [R1(t) - R1(0)] / r1,$$

where r1 is the longitudinal relaxivity per mmol Gd ($s^{-1}$ $mM^{-1}$). The mean concentration over the precontrast frames is then subtracted voxelwise, retaining signed baseline fluctuations. Hematocrit correction is applied only to the arterial input function, not to tissue concentration. Agent identity is read from BIDS ContrastBolusAgent, and r1 is selected from the supported table or supplied explicitly. Rohrer's bovine-plasma measurements at 37 °C [17] provide the 1.5/3 T defaults for Gd-DTPA (4.1/3.7), gadobutrol (5.2/5.0), gadoterate (3.6/3.5), and gadoteridol (4.1/3.7). Gadopiclenol uses a human-serum value of 12.8 measured at 1.41 T and a 3 T value of 11.6 extrapolated from nuclear magnetic relaxation dispersion profiles [18], with the 1.41 T value assigned to the 1.5 T lookup. The existing field bins select the 1.5 T entry at $B0 \leq 2.25$ T and the 3 T entry at $2.25 < B0 \leq 5$ T. No default is assigned above 5 T. Missing or unsupported agent/field combinations require an explicit finite positive r1; there is no generic-agent substitution. These in-vitro defaults are calibration assumptions rather than tissue-specific in-vivo measurements.

#### T1 mapping

T1 may be supplied as a map already aligned with the DCE series or estimated from variable flip angle (VFA) spoiled gradient-echo acquisitions using the Ernst signal equation [16]:

$S(\alpha) = S0 \times \sin(\alpha) \times (1 - E1) / [1 - E1 \times \cos(\alpha)]$,

where $E1 = \exp(-TR/T1)$. Following the DESPOT1 approach, weighted linear least squares (WLLS) estimates E1 and initializes the default nonlinear least-squares (NLLS) fit. A vectorized damped Gauss-Newton solver then fits the Ernst equation directly in signal space, parameterized by $R1 = 1/T1$ and using an analytical Jacobian. Signal-space fitting avoids the noise transformation introduced by the DESPOT1 linearization. Standalone weighted linear least-squares fitting is also available.

A spatially uniform T1 (by default 900, 1200 or 1700 ms at 1.5, 3 or 7 T) is used when T1 mapping is skipped, or when no usable VFA data or external map is available, VFA fitting produces no valid estimates, or registration QC rejects transfer of the VFA map to DCE space. QC compares tissue summaries with field-strength-specific reference intervals.

#### B1+ field mapping

Three B1+ correction sources are supported: (i) Actual Flip Angle Imaging (AFI) [19], using the short- and long-TR volumes of a two-volume AFI acquisition to estimate the actual-to-nominal flip-angle ratio; (ii) precomputed B1+ maps, encoded as percent of nominal flip angle, ratio, degrees or Siemens 0.1-degree units, with the encoding inferred from the BIDS suffix, Units field and value range or specified by the user; and (iii) no correction, represented by nominal flip angles.

B1+ ratio maps undergo mask-normalized Gaussian smoothing (default FWHM 8 mm), local outlier rejection, and field-strength-dependent clipping. AFI-derived maps are transferred to VFA or DCE space by registration of the AFI long-TR image. A precomputed map must be world-aligned with the DCE series; it is resampled to the DCE grid and, for VFA T1 mapping, transferred with the inverse of the VFA-to-DCE transform. Nominal flip angles are used when registration is rejected, and processing stops if a required transform is missing rather than applying an unregistered B1+ map.

#### DSC leakage correction

Boxerman–Schmainda–Weisskoff (BSW) leakage correction [20] estimates K1 and K2 independently in each voxel using the integral model:

$$C_{voxel}(t) = K1 \times \bar{C}(t) - K2 \times \int_0^t \bar{C}(\tau)\, d\tau,$$

where $\bar{C}(t)$ is a mean brain reference curve. Both K1 and K2 are estimated without sign constraints by ordinary least-squares regression over the complete baseline-centered dynamic series, and all finite K2 estimates are retained. Positive K2 represents a T1-dominant leakage contribution that reduces the observed ΔR2* curve and is added back during correction. Negative K2 retains the opposite-sign contribution associated with T2*-dominant extravasation, as well as possible residue-shape or transit mismatch relative to the reference curve.

Reference voxels are selected from motion-corrected raw signal. A voxel is retained when its mean signal over the final ten samples does not exceed its prebolus mean by more than one prebolus sample standard deviation. $\bar{C}(t)$ is the mean baseline-centered ΔR2* curve over the selected voxels, of which at least 250 are required for numerical stability. The corrected tissue curve is

$$C_{corrected}(t) = C_{tissue}(t) + K2 \times \int_0^t \bar{C}(\tau)\, d\tau.$$

If an eligible reference cannot be constructed, or leakage correction is disabled, the baseline-centered uncorrected series is used and K2 is not estimated.

Preload administration and BSW leakage correction are complementary strategies in consensus glioma protocols [21]. The administered preload, when supplied, and a prebolus baseline ΔR2* summary are recorded for QC.

#### DSC deconvolution methods

Perfusion is estimated from the tissue–AIF convolution relation

$C_{tissue}(t) = h(t) \otimes AIF(t)$, with $h(t) = F \times R(t)$,

where F is the relative flow coefficient ($s^{-1}$), R(t) is the tissue residue function, with R(0) = 1, and h(t) is the flow-scaled residue [14], so that F is the maximum of h(t). Reported rCBF is 60 × F ($min^{-1}$). Standard SVD and Tikhonov regularization discretize the convolution with the Sourbron product-integration (Volterra) scheme [22], oscillation-index SVD uses a block-circulant formulation, and CPI uses the continuous operator described below. The SVD-family methods retain signed residues. sSVD and Tikhonov integrate all N recovered samples by the trapezoidal rule; oSVD integrates its full 2N-sample circular period as the signed sample sum multiplied by the temporal resolution. Four methods are available:

(i) Standard SVD (sSVD) [23]: Singular value decomposition with threshold-based truncation (default: 0.2 of maximum singular value). Delay-sensitive; mismatch between AIF and tissue arrival times can bias CBF estimates.

(ii) Oscillation-index SVD (oSVD) [24]: Uses a block-circulant SVD formulation [24] for delay insensitivity, with adaptive regularization that selects the least-regularized threshold whose oscillation index falls below the published target of 0.095. This is the default method.

(iii) Tikhonov regularization [25,26]: penalized least squares on the full N-sample Volterra system, with a modified first-difference penalty that constrains the leading two residue samples and differences between later samples. L-curve selection [27] evaluates 100 log-spaced λ values from $10^{-3}$ to 10 times the largest singular value of the forward matrix. Using the implementation's curvature sign convention, it selects the strict interior curvature maximum nearest the upper λ bound, allowing either curvature sign, or the upper endpoint if no such maximum exists. Residue-integral MTT is unavailable when the signed area is non-positive or the residue peak is below the numerical tolerance; reported MTT is the separate AUC-CBV/CBF endpoint.

(iv) Control point interpolation (CPI) [28]: Bayesian non-parametric estimation of a continuous residue function using the signal-domain model described below.

*Control point interpolation*

CPI [28] fits the measured DSC magnitude signal directly. Before fitting, all tissue signals are scaled by a single factor to a median voxel pre-bolus baseline of 100, the mean of the S0 prior.

The intrinsic residue $R\theta(t)$ is a monotone piecewise cubic Hermite curve through movable control points. $R\theta(0) = 1$, and each later amplitude is the previous one multiplied by a positive ratio of at most one, so the residue is non-increasing and nonnegative; the final amplitude is estimated. Knot intervals are estimated on a log scale with a minimum of one third of a frame, and the last knot is fixed at the end of the acquisition. A further parameter scales the initial slope between zero and three times the first secant. The number of control points (5–12) is chosen per acquisition from the post-peak decay time of the mean tissue curve, within the limits set by the number of frames and the knot spacing, or is set by the user.

For flow F and signed AIF-to-tissue delay δ, the vascular ΔR2* is

$$C_v(t) = F \int C_a(\tau)\, R\theta(t - \delta - \tau)\, d\tau,$$

where $C_a$ is the input ΔR2* AIF, interpolated between samples with a shape-preserving cubic and convolved exactly over its polynomial segments. The predicted signal is

$$\hat{S}(t) = S0 \exp[-TE\, C_v(t)],$$

with S0 estimated jointly with F, the residue parameters and δ, and Gaussian noise of unknown precision. Up to three starting points are derived from a truncated circular deconvolution; each is fitted, and the valid fit with the highest variational free energy is kept.

With BSW leakage correction enabled, CPI jointly estimates a signed K2 using the BSW reference-curve term [20]. The voxelwise BSW estimate and its standard error define a Gaussian prior on K2, and uncertainty in the reference-region mean curve is propagated through additional nuisance parameters. Without leakage correction, these terms are omitted.

Optional dispersion convolves the AIF with a unit-area gamma kernel whose mode and rate are estimated [29]; it is off by default. In automatic mode, fits with and without dispersion are compared by the free energy of the voxel-independent VI fits and the higher is kept; with SVB, this choice is made before the spatial updates.

Priors are independent Gaussians on the fitted parameters: F, mean 0 (variance $10^6$), positive; S0, 100 ($10^6$), positive; log amplitude ratios, log 0.5 (0.67), at most zero; delay, 0 s (10 $s^2$); log knot intervals, log 8 s (0.40); initial-slope multiplier, 1 (1), within [0, 3]; and, with dispersion, log mode and log rate, log 2 (4), with the mode within 0.05–30 s and the rate within 0.05–20 $s^{-1}$. Delay and knot ranges follow from the acquisition duration. The priors regularize weakly identified parameters and therefore influence the estimates.

*Voxel-independent variational Bayesian inference*

CPI uses voxel-independent variational Bayesian inference (VI) [30] by default. The model is linearized around the current estimate, with an approximate Gaussian posterior for the parameters and a Gamma posterior for the noise precision. Updates are kept within the parameter bounds and backtracked when they do not increase the variational objective, for up to 100 iterations; a fit is labeled converged when the largest parameter change falls below $10^{-3}$. Estimates are posterior means, and the posterior covariance gives the rCBF standard deviation. Reported uncertainty is conditional on the input AIF and, when used, the BSW-derived K2 prior.

A fit is accepted when it is numerically valid, has finite positive rCBF, residue-integral rCBV and MTT, and has $R^2 \geq 0$, that is, it fits at least as well as the signal's temporal mean. Accepted fits that did not meet the convergence criterion are kept and flagged in QC. Rejected voxels are replaced by oSVD when fallback is enabled, using the BSW-corrected concentration curve (or the uncorrected curve without leakage correction), and are marked in a fallback mask. The rCBF, residue-integral rCBV and MTT maps may therefore contain oSVD values in marked voxels, where arrival delay and Tmax are left undefined. The primary AUC-based rCBV is computed independently of the deconvolution.

*Spatial variational Bayesian inference*

Spatial variational Bayes (SVB) is available as an alternative to VI, with the same model, priors and noise model. Following Penny et al. and Groves et al. [31,32], a six-neighbor M-prior with boundary correction replaces the independent prior on flow; the other parameters keep their priors, and voxels with rejected fits are excluded from the neighborhood. SVB starts from the VI solution and performs ten spatial updates, each accepted for a voxel only when it improves that voxel's objective given its neighbors. Its accuracy relative to VI is reported in Supplementary Section C.

*Perfusion parameters*

The primary rCBV is the indicator-dilution area-under-the-curve (AUC) ratio [20,23]:

$$rCBV = \left[\int_{BAT}^{end} C_{tissue}(t)\,dt\right] / \left[\int_{BAT}^{end} C_{AIF}(t)\,dt\right],$$

where BAT is the finalized baseline arrival reference described above, and integration continues to the end of the acquisition. $C_{tissue}$ is the baseline-centered tissue curve, corrected for leakage when BSW correction is enabled (see DSC leakage correction), and $C_{AIF}$ is the baseline-centered measured or user-supplied AIF. As the numerator and denominator share the same window, recirculation is included in both; its arterial and tissue contributions need not cancel, as the curves have different shapes. The other parameters are defined below; nCBV and nCBF require an NAWM reference.

| Parameter | Definition | Units |
| --- | --- | --- |
| rCBF | 60 × maximum of the flow-scaled residue h(t), with h(t) in $s^{-1}$ | Relative (1/min) |
| rCBV (AUC, primary) | AUC ratio above | Relative |

| Parameter | Definition | Units |
|---|---|---|
| rCBV (deconvolution) | Integral of h(t) | Relative |
| MTT | 60 × $rCBV_{AUC}$/rCBF for sSVD, oSVD and Tikhonov; integral of the fitted residue R(t) for CPI | Seconds |
| Tmax | Time of the recovered-residue maximum for sSVD and Tikhonov, signed residue-peak delay for oSVD, fitted AIF-to-tissue delay for CPI | Seconds |
| TTP | Time from the finalized baseline arrival reference to peak tissue concentration | Seconds |
| K2 | BSW leakage coefficient | $s^{-1}$ |
| nCBV | Tissue AUC/median tissue AUC in NAWM (equal to $rCBV_{AUC}$/median $rCBV_{AUC}$ in NAWM) | Dimensionless |
| nCBF | rCBF/median rCBF in NAWM | Dimensionless |

Residue-peak timing depends on sampling and regularization, so Tmax is not an absolute bolus-arrival time; in zero-delay simulations, oSVD Tmax was positive by approximately 1–3 s.

#### DCE pharmacokinetic analysis

##### *Effective-delay estimation*

Each model is fitted over AIF–tissue delays of 0–12 s in 0.1-s steps, with the delayed AIF evaluated by piecewise cubic Hermite interpolation (PCHIP) [33]. Delay evidence is pooled within an in-plane neighborhood of 2-mm radius, or the in-plane voxel diagonal if larger, including the center voxel. For each candidate delay, n·log(SSE), where n is the number of time points, is summed over neighbors with valid fits at every delay; the delay with the lowest sum is the shared reference delay. The reference is zero when fewer than two neighbors contribute or the summed objective varies by less than 6 across delays. A voxel-specific delay replaces the reference when it lowers BIC by at least 6, counting the delay as an additional parameter. Disabling delay estimation fixes the delay at zero. The fitted delay is an effective model alignment rather than a vascular transit time, and values at 12 s may reflect the search limit.

##### *Pharmacokinetic models*

(i) Standard Tofts model [34]: $C_t(t) = K^{trans} \times [C_p \otimes \exp(-k_{ep} \times t)]$, where $k_{ep} = K^{trans} / v_e$. Two free parameters: $K^{trans}$ (volume transfer constant, $min^{-1}$) and $v_e$ (extravascular extracellular volume fraction).

(ii) Extended Tofts model [13]: $C_t(t) = v_p \times C_p(t) + K^{trans} \times [C_p \otimes \exp(-k_{ep} \times t)]$. Three free parameters: $K^{trans}$, $v_e$, and $v_p$.

(iii) Patlak model [35]: $C_t(t) = v_p \times C_p(t) + K^{trans} \times \int_0^t C_p(\tau)\, d\tau$. Two free parameters, $K^{trans}$ and $v_p$.

(iv) Two-compartment exchange model (2CXM) [36,37]: A four-parameter model (plasma flow $F_p$, permeability-surface area product PS, $v_e$, $v_p$) from which $K^{trans} = E \times F_p / 100$, where $E = PS / (F_p + PS)$ and $F_p$ and PS are expressed in mL min$^{-1}$ 100 mL$^{-1}$.

The default inference uses bounded least squares at each delay: linear fitting for Patlak, rate profiling with constrained linear solutions for Tofts and Extended Tofts, and a trust-region solver for 2CXM. Validation tables label this default least-squares pathway BLS. Fits are bounded to 0–5 min$^{-1}$ for $K^{trans}$, 0–1 for $v_e$ and $v_p$ with $v_e + v_p \leq 1$, and 0–500 mL min$^{-1}$ 100 mL$^{-1}$ for $F_p$ and PS.

Fitting is restricted to brain voxels with finite concentration curves and positive enhancement and, when a tSNR threshold is configured, adequate baseline tSNR.

Automatic model selection compares eligible models voxelwise by the Bayesian information criterion (BIC) [38]. Patlak and Extended Tofts are always eligible; PMRIPrep restricts 2CXM eligibility to a temporal resolution (BIDS RepetitionTime, or the spacing of the supplied time axis) of 2 s or finer, motivated by the sampling sensitivity of plasma-flow estimation [39] (Section J). The converged model with the lowest BIC is selected, but an Extended Tofts or 2CXM fit replaces a finite Patlak fit only when its BIC is at least 6 lower. 2CXM is further admitted only when, at its selected delay, an Extended Tofts fit detects both $K^{trans}$ and $v_p$ (each more than two standard errors from its bounds) and 2CXM lowers BIC relative to that fit. Standard Tofts is available by explicit selection but excluded from automatic selection, as its zero-plasma-volume assumption can cause the vascular contribution to be absorbed into $K^{trans}$ [13]. Optional VI [30] refines the selected model at the selected delay and reports posterior uncertainty without changing either; the DSC-informed $v_p$ prior, when available, enters only this step. If VI fails numerically, does not converge or returns an invalid estimate, the BLS estimate at the same model and delay is retained.

After selection, the selected fit is checked for plausibility. Nonfinite fits, fits at the upper rate bounds, and fits with zero $K^{trans}$ (Tofts and Extended Tofts), zero $v_e$ (Tofts, Extended Tofts and 2CXM), or zero $F_p$ or PS (2CXM) are rejected; Patlak may retain $K^{trans} = 0$. A rejected fit leaves the voxel's estimates missing rather than falling back to another model.

*Uncertainty*

BLS uncertainty maps report standard errors from the fitted-model Jacobian and residual variance, and VI maps report marginal posterior standard deviations. Both are conditional on the applied delay, the selected model, the concentration curves and the AIF; they do not include uncertainty from model selection, baseline estimation, T1 or B1+ mapping, or signal-to-concentration conversion. Both are scaled by heuristic factors derived from a base factor B: B = 2 for an AIF accepted through the higher scaling allowance with at most sixfold scaling, flagged for venous timing, or whose final selected candidate subset fails its repeated AIF checks; B = 3 for more than sixfold scaling or the population AIF; otherwise B = 1. Uncertainties are multiplied by B for $K^{trans}$, $F_p$ and PS, by (B + 1)/2 for $v_p$, and are unchanged for $v_e$.

*DCE QC mask*

A binary DCE QC mask accompanies the parameter maps. Its criteria are applied to retained fits and are separate from the estimation bounds. A voxel passes when its fit converged and was retained, $0 < K^{trans} < 5\ min^{-1}$, $R^2 > 0$ (the fit outperforms the curve's temporal mean), $0 < k_{ep} \leq 10\ min^{-1}$ for reversible models, and the fitted volume fractions lie strictly inside their limits (tolerance $10^{-6}$), including $v_e + v_p < 1$ for Extended Tofts and 2CXM. Its primary parameters ($K^{trans}$ and $v_p$ for Patlak, $K^{trans}$ and $v_e$ for Tofts, and $K^{trans}$, $v_e$ and $v_p$ for Extended Tofts and 2CXM) must also be identified at the selected BLS fit, including when VI supplies the reported map: the normalized information matrix must have a reciprocal condition number of at least $10^{-10}$, so that no parameter combination is effectively undetermined; each parameter must lie more than two standard errors from its bounds; $v_e + v_p$ must lie more than two standard errors below 1; and its effective relative uncertainty, combining the standard error with its variation across the delay profile, must not exceed 50%. $F_p$ and PS identification is reported separately. A tSNR threshold can be added. The mask does not censor the maps. It defines separately labeled QC-filtered summaries of the pharmacokinetic maps, including regional distributions, NAWM references and fractional-volume summaries; model-free DCE statistics are not filtered. As near-zero values often fail identification, QC-filtered summaries can shift upward. The QC sidecar records the per-criterion rejection counts.

#### DSC-informed DCE prior

The DSC-informed $v_p$ prior is available only with Bayesian DCE fitting. Its spatially varying nominal natural-scale center is

$$\mu(v_p) = 0.010 \times nCBV,$$

where 0.010 is a fixed calibration assumption for NAWM plasma volume rather than a subject-specific estimate [40,41]. This scaling assumes equal microvascular hematocrit in the voxel and the NAWM reference. The prior is Gaussian in bounded log-odds $v_p$, with nominal natural-scale center $\mu(v_p)$ and scale $\max[0.60 \times \mu(v_p), 0.001]$; it permits departures from the DSC-derived center according to the relative prior and likelihood information. It replaces only the default $v_p$ prior, which is Gaussian in the same transform with natural-scale mean and SD equal to the midpoint and one quarter of the model's typical $v_p$ range: 0.001–0.04 for Extended Tofts and Patlak (mean 0.0205, SD 0.00975) and 0.01–0.10 for 2CXM (mean 0.055, SD 0.0225). The $K^{trans}$, $v_e$, $F_p$, PS and noise priors are unchanged. Section L characterizes these choices under DCE noise and errors in the prior. BIC model selection is done without the prior; it is applied only during Bayesian refinement of a selected model that estimates $v_p$, in voxels with a finite, positive prior center no greater than 0.2. Some retained fits in a prior-eligible analysis may therefore remain unconstrained.

The prior requires a leakage-corrected, NAWM-normalized DSC CBV map, finite DSC tissue CNR with a mean of at least 3 and at least 84% of brain voxels with CNR > 4, at most 1% negative values among finite brain nCBV values, and registration support for transfer (transformed-foreground Dice ≥ 0.90 for DSC, and Dice ≥ 0.80 with Pearson correlation ≥ 0.03 for DCE). These registration thresholds screen whether transfer is supported and were not calibrated as measures of anatomical registration accuracy. The CNR coverage threshold is scaled by the administered gadolinium dose relative to 0.1 mmol/kg. It is rejected when its whole-brain median lies outside 0.005–0.030, a check for gross scaling or normalization errors rather than a reference range for tissue $v_p$. The same criteria apply to explicitly and automatically selected models that estimate $v_p$. Users may disable it. QC records the decision, its reason, and the fraction of retained fits that used the prior.

##### *NAWM-normalized $v_p$ and CBV concordance*

DSC–DCE concordance is assessed after dividing DCE $v_p$ and DSC nCBV by their medians in the same DCE-space NAWM mask and smoothing both with matched Gaussian kernels ($\sigma$ = 1 mm). Smoothing is normalized by each map's finite-data mask. At least 500 jointly valid NAWM voxels are required for the median normalization. Spearman and Pearson correlations are computed over jointly valid positive voxels in the whole brain and, when available, the lesion, each also requiring at least 500 voxels. The comparison is labeled independent only when no retained DCE fit used the DSC-informed prior; otherwise it is labeled prior-informed and is not treated as independent cross-modal validation.

#### Output summary

Outputs are written as BIDS-Derivatives. Anatomical derivatives include the processed structural references, brain mask, tissue probability maps, transforms and, when available, tumor labels and the NAWM mask; an accepted VFA-derived T1 map is published in DCE space, whereas external and uniform T1 maps and B1+ maps are used without being republished. DSC derivatives include rCBF, AUC-based and residue-integral rCBV, MTT, Tmax, TTP, nCBF, nCBV, tSNR, CNR, PSR and, with BSW correction, a signed K2 map; CPI adds fit $R^2$, a fallback mask and rCBF uncertainty. DCE derivatives include the applicable pharmacokinetic maps ($K^{trans}$, $v_e$, $k_{ep}$, $v_p$, and for 2CXM, $F_p$, PS and extraction fraction), model-free enhancement and timing maps (initial area under the curve (IAUC) and signal enhancement ratio (SER) when the acquisition covers their time windows), the selected-delay map, model-selection and fit-quality maps, the DCE QC mask, and uncertainty maps. JSON sidecars record regional summaries, with separately labeled QC-filtered summaries for pharmacokinetic maps, together with units, QC and provenance; an HTML report summarizes each subject. Optional smoothing produces additional derivatives rather than replacing the unsmoothed maps.

#### Supplementary validation results

The validation experiments tested numerical recovery, external comparability, repeatability, availability, and sensitivity to acquisition and modeling assumptions. The reference objects answer different questions and are not pooled into one performance estimate.

Unless stated otherwise, signed bias is relative to ground truth and positive values denote overestimation. Each table identifies its distribution summary; the DSC recovery tables generally report medians and 95th percentiles. Absolute log error is |log(estimate/truth)|. The S1 tables report absolute percentage error for comparison with conventional DSC recovery results. The S2–S4 tables use absolute log error to summarize multiplicative error; for small deviations, an absolute log error of 0.1 corresponds to approximately 10% relative error. For test–retest pairs, the conventional subject-level coefficient of variation (CV) uses the sample standard deviation of the two measurements. The corresponding calculation using the population standard deviation, matching the published OSIPI methodology, is also reported. Repeatability coefficients are 2.77 times the corresponding pooled within-subject CV. Correlations are Pearson correlations unless identified otherwise.

##### A. DSC deconvolution recovery in internal simulations

Sections A–D use internally generated simulations to assess implementation recovery and sensitivity; they are not independent external validation. The forward generator used continuous arterial-input and residue functions, with tissue curves computed by numerical integration independent of each fitting method's discretization, followed by Rician magnitude noise. Full-integral CBV denotes the volume obtained from the complete simulated residue, whereas within-acquisition AUC-CBV integrates from bolus arrival through the final acquired frame. Tables S1a-3 and S1b-3 compare AUC-CBV with full-integral CBV and therefore include acquisition-window truncation; Table S2-1 compares AUC-CBV with a no-leakage truth integrated over the same acquisition window. The comparison used the CBV, CBF, and MTT ranges of Mouridsen et al. [42]: full-integral CBV was fixed at 4% and CBF varied from 10 to 70 mL/100 g/min, giving MTT values from 24 to 3.43 s. Exponential, triangular, and box residues were evaluated without delay. A common AIF and tissue-curve normalization set the exponential-residue 4%-CBV, 60-mL/100-g/min reference to the published 40% tissue signal-drop convention [42] without changing perfusion truth. Each noisy condition used 200 realizations; each method therefore contributed 4,200 observations per noisy SNR and 21 noiseless observations.

Residue-integral MTT was the primary model-recovery endpoint. PMRIPrep's default reported MTT is AUC-CBV/CBF for the SVD methods and fitted-residue MTT for CPI.

Tables S1a-1 and S1a-2 report the pooled distribution across all flows and residue families and the residue-specific SNR-50 comparison underlying Figure 2, respectively. Values are median / 95th-percentile absolute percentage error among finite estimates. CBF and reported MTT were finite for every observation. Tikhonov residue MTT was finite in 10,648/12,621 observations; 1,973 estimates were unavailable (462 at SNR 100, 621 at SNR 50 and 890 at SNR 20). The other methods had complete residue-MTT availability. All 12,621 CPI values came from CPI rather than oSVD fallback: 12,424 met the solver convergence criterion, and 197 did not but passed PMRIPrep's CPI fit-acceptance checks.

##### Fixed-4%-CBV DSC recovery

**Supplementary Table S1a-1 | Pooled fixed-4%-CBV recovery by noise condition.**

| Method | SNR | CBF error (%) | Residue-MTT error (%) | Reported-MTT error (%) | Finite residue MTT / total |
|---|---|---|---|---|---|
| sSVD | Noiseless | 9.9 / 25.1 | 9.5 / 21.6 | 10.1 / 23.4 | 21/21 |
| sSVD | 100 | 11.0 / 26.7 | 11.1 / 25.5 | 11.1 / 27.5 | 4,200/4,200 |
| sSVD | 50 | 12.9 / 32.0 | 13.0 / 31.3 | 13.1 / 32.3 | 4,200/4,200 |
| sSVD | 20 | 17.4 / 58.3 | 20.5 / 53.4 | 20.0 / 53.4 | 4,200/4,200 |
| oSVD | Noiseless | 13.6 / 32.6 | 12.9 / 24.6 | 12.9 / 24.6 | 21/21 |
| oSVD | 100 | 16.6 / 40.2 | 16.6 / 38.5 | 16.6 / 38.5 | 4,200/4,200 |
| oSVD | 50 | 18.9 / 47.7 | 19.0 / 52.3 | 19.0 / 52.2 | 4,200/4,200 |
| oSVD | 20 | 22.8 / 70.2 | 25.6 / 76.4 | 25.6 / 76.3 | 4,200/4,200 |
| Tikhonov | Noiseless | 20.9 / 45.9 | 17.9 / 31.5 | 17.4 / 32.4 | 21/21 |
| Tikhonov | 100 | 14.2 / 27.3 | 40.3 / 184.1 | 14.0 / 32.9 | 3,738/4,200 |
| Tikhonov | 50 | 15.1 / 32.4 | 45.5 / 242.4 | 15.4 / 46.4 | 3,579/4,200 |
| Tikhonov | 20 | 19.9 / 44.6 | 64.7 / 349.3 | 23.3 / 88.8 | 3,310/4,200 |
| CPI | Noiseless | 17.4 / 171.8 | 14.7 / 63.2 | 14.7 / 63.2 | 21/21 |
| CPI | 100 | 7.6 / 72.2 | 10.2 / 41.2 | 10.2 / 41.2 | 4,200/4,200 |
| CPI | 50 | 13.1 / 82.1 | 17.5 / 49.2 | 17.5 / 49.2 | 4,200/4,200 |
| CPI | 20 | 21.6 / 85.7 | 23.6 / 86.4 | 23.6 / 86.4 | 4,200/4,200 |

The higher noiseless CPI CBF error than at SNR 100 was concentrated in the short-MTT exponential cases and does not indicate a benefit from added noise. The continuous generator and CPI's sampled forward model differ slightly; without noise, CPI accommodated that difference by trading higher CBF against shorter MTT and small positive delays. At SNR 100, the parameter priors constrained this trade-off.

**Supplementary Table S1a-2 | Residue-specific recovery at SNR 50.**

| Method | Residue | CBF error (%) | Residue-MTT error (%) | Reported-MTT error (%) | Finite residue MTT / total |
|---|---|---|---|---|---|
| sSVD | Exponential | 13.0 / 28.0 | 13.2 / 37.2 | 14.5 / 39.7 | 1,400/1,400 |
| sSVD | Triangle | 6.2 / 23.6 | 7.3 / 21.8 | 7.4 / 22.5 | 1,400/1,400 |
| sSVD | Box | 20.9 / 35.8 | 18.6 / 30.9 | 17.8 / 30.4 | 1,400/1,400 |
| oSVD | Exponential | 20.3 / 39.8 | 24.0 / 67.8 | 24.1 / 67.5 | 1,400/1,400 |
| oSVD | Triangle | 12.0 / 31.7 | 12.3 / 39.7 | 12.3 / 39.7 | 1,400/1,400 |
| oSVD | Box | 26.4 / 58.7 | 21.9 / 38.9 | 21.9 / 38.9 | 1,400/1,400 |
| Tikhonov | Exponential | 20.9 / 35.1 | 53.1 / 301.4 | 26.2 / 58.6 | 1,219/1,400 |
| Tikhonov | Triangle | 9.5 / 21.9 | 46.2 / 232.3 | 10.5 / 31.7 | 1,185/1,400 |
| Tikhonov | Box | 16.2 / 31.0 | 38.2 / 182.0 | 14.4 / 27.6 | 1,175/1,400 |
| CPI | Exponential | 13.2 / 30.2 | 26.1 / 57.1 | 26.1 / 57.1 | 1,400/1,400 |
| CPI | Triangle | 9.3 / 31.2 | 11.0 / 40.0 | 11.0 / 40.0 | 1,400/1,400 |
| CPI | Box | 36.5 / 101.2 | 20.4 / 45.8 | 20.4 / 45.8 | 1,400/1,400 |

In the pooled SNR-50 results in Table S1a-1, CPI and sSVD had similar median CBF error, but CPI had a larger 95th-percentile error, and the residue-specific breakdown showed a high-error CPI tail for box residues. sSVD had the lowest pooled residue-MTT error. Reported and residue-integral MTT errors were similar for sSVD and oSVD; Tikhonov reported MTT had lower error than its finite residue-integral estimates. At the lowest tested true CBF (10 mL/100 g/min), the median SNR-50 CBF estimates were 11.9, 12.4, 9.9, and 10.3 mL/100 g/min for sSVD, oSVD, Tikhonov, and CPI, respectively.

**Supplementary Table S1a-3 | Primary AUC-CBV recovery on the fixed-4%-CBV grid.**

| SNR | AUC-CBV error (%) | Median signed error (%) |
|---|---:|---:|
| Noiseless | 0.1 / 0.1 | -0.1 |
| 100 | 2.7 / 8.1 | -0.1 |
| 50 | 5.4 / 16.2 | -0.3 |
| 20 | 13.8 / 40.1 | -0.1 |

The primary AUC-CBV calculation is common to all four deconvolution methods. It remained nearly unbiased when pooled across the fixed-4%-CBV grid; its error increased with magnitude noise.

A separate fixed-flow stress matrix held CBF at 60 mL/100 g/min while crossing five residue shapes (exponential [gamma shape 1], gamma shape 3, gamma shape 10, triangle, and box), MTT values of 5 and 16 s, delays of 0 and 3 s, and four noise conditions (noiseless and baseline-signal SNR 100, 50, and 20). The MTT and CBV truths therefore varied together: full-integral CBV was 5% at 5 s and 16% at 16 s. The 16% condition is a duration and model-misspecification stress rather than a conventional brain-tissue reference. Each method contributed 4,000 observations per noisy SNR and 20 noiseless observations. CBF and reported MTT were finite throughout. Tikhonov residue MTT was finite in 11,280/12,020 observations, with 155, 236 and 349 unavailable at SNR 100, 50 and 20, respectively; the other methods had complete residue-MTT availability. All 12,020 CPI outputs came from CPI rather than fallback: 11,793 met the solver convergence criterion, and 227 did not but passed PMRIPrep's CPI fit-acceptance checks. Reported MTT is AUC-CBV/CBF for the SVD methods and fitted-residue MTT for CPI.

##### Fixed-flow duration, delay, and residue-shape stress test

**Supplementary Table S1b-1 | Pooled fixed-flow duration-and-delay recovery by noise condition.**

| Method | SNR | CBF error (%) | Residue-MTT error (%) | Reported-MTT error (%) | Finite residue MTT / total |
|---|---|---|---|---|---|
| sSVD | Noiseless | 12.3 / 28.0 | 11.3 / 26.8 | 11.1 / 24.9 | 20/20 |
| sSVD | 100 | 12.0 / 31.9 | 11.2 / 39.0 | 11.1 / 36.0 | 4,000/4,000 |
| sSVD | 50 | 12.1 / 31.5 | 11.8 / 40.0 | 11.6 / 38.2 | 4,000/4,000 |
| sSVD | 20 | 15.4 / 38.8 | 15.4 / 52.0 | 15.3 / 49.0 | 4,000/4,000 |
| oSVD | Noiseless | 18.7 / 24.5 | 15.8 / 19.7 | 15.8 / 19.7 | 20/20 |
| oSVD | 100 | 14.6 / 40.4 | 13.7 / 31.1 | 13.7 / 31.1 | 4,000/4,000 |
| oSVD | 50 | 17.0 / 50.0 | 16.1 / 38.8 | 16.1 / 38.7 | 4,000/4,000 |
| oSVD | 20 | 21.4 / 77.4 | 21.7 / 53.1 | 21.6 / 52.8 | 4,000/4,000 |
| Tikhonov | Noiseless | 21.4 / 55.6 | 25.7 / 49.8 | 25.3 / 48.7 | 20/20 |
| Tikhonov | 100 | 11.4 / 31.4 | 21.6 / 117.2 | 10.8 / 45.1 | 3,845/4,000 |
| Tikhonov | 50 | 12.9 / 38.6 | 24.9 / 148.2 | 12.5 / 62.4 | 3,764/4,000 |
| Tikhonov | 20 | 16.9 / 45.9 | 29.6 / 207.8 | 16.8 / 89.3 | 3,651/4,000 |
| CPI | Noiseless | 29.0 / 89.5 | 22.3 / 47.0 | 22.3 / 47.0 | 20/20 |
| CPI | 100 | 6.7 / 38.6 | 6.1 / 27.3 | 6.1 / 27.3 | 4,000/4,000 |
| CPI | 50 | 8.9 / 66.0 | 8.5 / 39.7 | 8.5 / 39.7 | 4,000/4,000 |
| CPI | 20 | 15.1 / 77.2 | 14.0 / 49.3 | 14.0 / 49.3 | 4,000/4,000 |

Values are pooled across the five residue families and are median / 95th-percentile absolute percentage error.

**Supplementary Table S1b-2 | MTT-by-delay recovery at SNR 50.**

| Method | MTT (s) | Delay (s) | CBF error (%) | Residue-MTT error (%) | Reported-MTT error (%) | Finite residue MTT / total |
|---|---|---|---|---|---|---|
| sSVD | 5 | 0 | 12.5 / 30.9 | 12.6 / 26.8 | 12.5 / 27.0 | 1,000/1,000 |
| sSVD | 5 | 3 | 16.0 / 36.9 | 19.6 / 65.7 | 17.7 / 62.7 | 1,000/1,000 |
| sSVD | 16 | 0 | 11.9 / 22.9 | 11.0 / 18.6 | 10.7 / 18.6 | 1,000/1,000 |
| sSVD | 16 | 3 | 10.4 / 21.9 | 10.1 / 18.8 | 10.2 / 18.8 | 1,000/1,000 |
| oSVD | 5 | 0 | 14.5 / 39.8 | 15.5 / 41.2 | 15.5 / 41.6 | 1,000/1,000 |
| oSVD | 5 | 3 | 15.4 / 41.7 | 16.0 / 43.1 | 15.8 / 43.1 | 1,000/1,000 |
| oSVD | 16 | 0 | 20.1 / 59.9 | 16.7 / 37.2 | 16.7 / 37.1 | 1,000/1,000 |
| oSVD | 16 | 3 | 19.3 / 53.3 | 16.6 / 35.5 | 16.5 / 35.5 | 1,000/1,000 |
| Tikhonov | 5 | 0 | 12.2 / 26.8 | 62.9 / 219.4 | 12.6 / 36.7 | 811/1,000 |
| Tikhonov | 5 | 3 | 26.5 / 46.7 | 52.7 / 162.7 | 35.0 / 90.1 | 956/1,000 |
| Tikhonov | 16 | 0 | 12.4 / 25.3 | 18.1 / 60.3 | 10.9 / 20.6 | 997/1,000 |
| Tikhonov | 16 | 3 | 8.0 / 22.1 | 11.8 / 38.3 | 7.8 / 27.6 | 1,000/1,000 |
| CPI | 5 | 0 | 17.5 / 76.2 | 17.7 / 43.8 | 17.7 / 43.8 | 1,000/1,000 |
| CPI | 5 | 3 | 19.2 / 82.0 | 18.7 / 44.9 | 18.7 / 44.9 | 1,000/1,000 |
| CPI | 16 | 0 | 4.5 / 19.4 | 4.4 / 15.9 | 4.4 / 15.9 | 1,000/1,000 |
| CPI | 16 | 3 | 4.7 / 17.4 | 4.3 / 15.4 | 4.3 / 15.4 | 1,000/1,000 |

The MTT-by-delay breakdown at SNR 50 showed that delay substantially increased short-MTT CBF and reported-MTT error for sSVD and Tikhonov. Tikhonov residue-MTT summaries use the finite counts shown in each row. CPI was more accurate in the long-MTT strata but retained broad short-MTT CBF tails. Each row pools five residue families and 200 realizations per family.

**Supplementary Table S1b-3 | Primary AUC-CBV recovery in the duration-and-delay stress test.**

| SNR | AUC-CBV error (%) | Median signed error (%) |
|---|---:|---:|
| Noiseless | 0.1 / 0.2 | -0.1 |
| 100 | 1.3 / 5.4 | -0.1 |
| 50 | 2.5 / 11.5 | -0.2 |
| 20 | 6.2 / 27.4 | -0.2 |

AUC-CBV was again common to all methods. Noiseless AUC-CBV error was negligible, but noise increased error more strongly at MTT 5 s than at 16 s.

#### B. CPI leakage and primary AUC-CBV validation

Leakage was simulated either with the BSW model (positive or negative K2) or with a separate two-compartment exchange model. The simulations were evaluated at baseline-signal SNR 50 and 20 with matched and prolonged tissue transit. Each stratum contained 50 independently noised observations. Primary AUC-CBV was evaluated separately from CPI's fitted-residue parameters. With matched tissue and reference transit times, BSW-corrected AUC-CBV had little bias without leakage or with BSW-form leakage; exchange leakage retained approximately −12% bias. With 16-s tissue MTT against the 5-s reference, it remained approximately 51% low even without leakage, while uncorrected within-acquisition AUC-CBV was nearly unbiased. This isolates the reference-transit limitation from signal noise. In the exchange-model simulations, the leakage waveform differed from the BSW leakage term used in fitting.

BSW+ and BSW− denote positive and negative K2 values generated with the BSW leakage model.

**Supplementary Table S2-1 | Primary AUC-CBV recovery with leakage.**

| Leakage | MTT (s) | SNR | Corrected AUC-CBV error | Uncorrected AUC-CBV error | Corrected bias (%) |
|---|---|---|---|---|---|
| None | 5 | 50 | 0.016 / 0.057 | 0.043 / 0.136 | -0.7 |
| None | 5 | 20 | 0.050 / 0.117 | 0.140 / 0.380 | -0.5 |
| None | 16 | 50 | 0.715 / 0.756 | 0.018 / 0.047 | -51.1 |
| None | 16 | 20 | 0.704 / 0.793 | 0.039 / 0.111 | -50.5 |
| BSW+ | 5 | 50 | 0.021 / 0.063 | 0.708 / 0.918 | +0.2 |
| BSW+ | 5 | 20 | 0.044 / 0.128 | 0.691 / 1.179 | -1.5 |
| BSW+ | 16 | 50 | 0.709 / 0.750 | 0.314 / 0.364 | -50.8 |
| BSW+ | 16 | 20 | 0.730 / 0.822 | 0.333 / 0.492 | -51.8 |
| BSW− | 5 | 50 | 0.019 / 0.067 | 0.411 / 0.476 | -0.2 |
| BSW− | 5 | 20 | 0.037 / 0.130 | 0.432 / 0.530 | +0.8 |
| BSW− | 16 | 50 | 0.721 / 0.765 | 0.239 / 0.267 | -51.4 |
| BSW− | 16 | 20 | 0.720 / 0.819 | 0.239 / 0.324 | -51.3 |
| Exchange T1 | 5 | 50 | 0.126 / 0.162 | not finite | -11.8 |
| Exchange T1 | 5 | 20 | 0.128 / 0.247 | not finite | -12.0 |
| Exchange T1 | 16 | 50 | 0.690 / 0.721 | 2.823 / 3.460 | -49.8 |
| Exchange T1 | 16 | 20 | 0.690 / 0.787 | 2.817 / 4.962 | -49.8 |

| Leakage | MTT (s) | SNR | Corrected AUC-CBV error | Uncorrected AUC-CBV error | Corrected bias (%) |
|---|---|---|---|---|---|
| Exchange T2* | 5 | 50 | 0.119 / 0.180 | 0.543 / 0.592 | -11.3 |
| Exchange T2* | 5 | 20 | 0.134 / 0.284 | 0.548 / 0.700 | -12.5 |
| Exchange T2* | 16 | 50 | 0.731 / 0.772 | 0.164 / 0.216 | -51.9 |
| Exchange T2* | 16 | 20 | 0.747 / 0.864 | 0.172 / 0.254 | -52.6 |

The two error columns report median / 95th-percentile absolute log error against the no-leakage AUC-CBV truth integrated over the same acquisition window. Corrected bias is the median signed percentage error of the BSW-corrected estimate. Uncorrected AUC-CBV error is marked not finite when the uncorrected tissue area was nonpositive and its log error was therefore undefined. The BSW-corrected estimate was available in all 1,000 cases. Exchange T1 and Exchange T2* denote T1- and T2*-dominant exchange leakage.

**Supplementary Table S2-2 | CPI fitted-parameter recovery with leakage at SNR 50.**

| Leakage | MTT (s) | CBF error | MTT error | K2 target ($s^{-1}$) | Median K2 ($s^{-1}$) |
|---|---|---|---|---|---|
| None | 5 | 0.129 / 0.271 | 0.065 / 0.155 | 0 | +0.0002 |
| None | 16 | 0.077 / 0.150 | 0.051 / 0.121 | 0 | +0.0003 |
| BSW+ | 5 | 0.133 / 0.295 | 0.058 / 0.148 | +0.0050 | +0.0052 |
| BSW+ | 16 | 0.062 / 0.148 | 0.028 / 0.107 | +0.0084 | +0.0086 |
| BSW− | 5 | 0.119 / 0.255 | 0.051 / 0.142 | -0.0050 | -0.0047 |
| BSW− | 16 | 0.050 / 0.139 | 0.031 / 0.100 | -0.0084 | -0.0084 |
| Exchange T1 | 5 | 0.240 / 0.369 | 0.105 / 0.215 | +0.0188 | +0.0192 |
| Exchange T1 | 16 | 0.121 / 0.169 | 0.080 / 0.125 | +0.0137 | +0.0278 |
| Exchange T2* | 5 | 0.259 / 0.420 | 0.135 / 0.268 | -0.0083 | -0.0076 |
| Exchange T2* | 16 | 0.085 / 0.145 | 0.126 / 0.172 | -0.0113 | +0.0009 |

CBF and fitted-residue MTT values are median / 95th-percentile absolute log error. For the exchange simulations, the K2 target is the value that best represents the simulated leakage waveform under the BSW model; it is not the exchange-model permeability parameter. When leakage was generated with the BSW model, CPI recovered the correct K2 sign in every stratum. In the exchange-model simulations at 16-s MTT, the fitted K2 was approximately twice the target for T1-dominant leakage and had the wrong sign for T2*-dominant leakage. Thus, the BSW model did not accurately describe these simulated leakage curves when tissue transit was prolonged. The exchange conditions also retained larger CBF and MTT errors. All 500 fits returned CPI estimates; no oSVD fallback was used.

#### C. CPI spatial variational Bayesian inference

VI and SVB were compared on the same 20 independently noised fields (five per pattern/SNR combination), using ten spatial update iterations for SVB. All 2,560 outputs per method were available and none used fallback. Recovery and spatial behavior were nearly identical. Both showed positive regional CBF bias of approximately 13% at SNR 50 and approximately 15% in homogeneous fields at SNR 20; spatial regularization did not remove this bias. Median runtime per 8 × 8 × 2 field was 12–15 s for VI and 21–28 s for SVB.

##### CPI VI and SVB recovery and spatial behavior

**Supplementary Table S3-1 | CPI VI and SVB saved-map recovery.**

| Pattern | SNR | Method | CBF error | MTT error | CBV error |
|---|---|---|---|---|---|
| Homogeneous | 50 | VI | 0.113 / 0.248 | 0.058 / 0.187 | 0.067 / 0.120 |
| Homogeneous | 50 | SVB | 0.114 / 0.247 | 0.058 / 0.186 | 0.067 / 0.120 |
| Homogeneous | 20 | VI | 0.144 / 0.361 | 0.092 / 0.292 | 0.138 / 0.279 |
| Homogeneous | 20 | SVB | 0.144 / 0.355 | 0.090 / 0.290 | 0.138 / 0.279 |
| Piecewise | 50 | VI | 0.115 / 0.235 | 0.063 / 0.173 | 0.064 / 0.137 |
| Piecewise | 50 | SVB | 0.115 / 0.234 | 0.063 / 0.173 | 0.064 / 0.137 |
| Piecewise | 20 | VI | 0.131 / 0.377 | 0.083 / 0.282 | 0.140 / 0.273 |
| Piecewise | 20 | SVB | 0.130 / 0.369 | 0.084 / 0.276 | 0.140 / 0.273 |

Errors are median / 95th-percentile absolute log error across saved-map voxels.

**Supplementary Table S3-2 | CPI VI and SVB spatial behavior.**

| Pattern | SNR | Method | Bias (%) | Regional SD | Edge mixing | Lesion contrast | Posterior SD |
|---|---|---|---|---|---|---|---|
| Homogeneous | 50 | VI | 12.9 | 0.088 | — | — | 0.151 |
| Homogeneous | 50 | SVB | 12.9 | 0.088 | — | — | 0.151 |

| Pattern | SNR | Method | Bias (%) | Regional SD | Edge mixing | Lesion contrast | Posterior SD |
|---|---|---|---|---|---|---|---|
| Homogeneous | 20 | VI | 14.9 | 0.162 | — | — | 0.224 |
| Homogeneous | 20 | SVB | 14.8 | 0.160 | — | — | 0.223 |
| Piecewise | 50 | VI | 13.0 / 11.6 / 11.8 | 0.085 / 0.089 / 0.074 | -0.051 | 1.110 | 0.152 |
| Piecewise | 50 | SVB | 13.1 / 11.6 / 11.5 | 0.085 / 0.088 / 0.072 | -0.048 | 1.103 | 0.152 |
| Piecewise | 20 | VI | 14.8 / 13.9 / 8.9 | 0.184 / 0.126 / 0.129 | -0.004 | 0.996 | 0.221 |
| Piecewise | 20 | SVB | 14.9 / 13.8 / 8.1 | 0.181 / 0.123 / 0.121 | 0.001 | 0.979 | 0.219 |

Spatial summaries are medians across five fields per condition; posterior SD is the median across their 640 voxels. Piecewise-region values are ordered left / right / lesion. Regional and posterior SD are normalized by true CBF; edge mixing is normalized boundary displacement, and lesion contrast is the estimated-to-true contrast ratio.

#### D. CPI dispersion option

Simulated dispersion was absent, gamma-shaped, or lognormal, a shape the gamma model cannot match exactly. Fitting with gamma dispersion substantially improved CBF and MTT for noisy matched and misspecified dispersed inputs. At SNR 50, automatic selection preserved nearly all of the matched-gamma recovery gain, retaining gamma dispersion in 17/20 matched-gamma observations and the no-dispersion fit in 15/20 no-dispersion controls. In the single noiseless gamma-dispersion observation, the gamma-dispersion fit increased CBF and MTT absolute log error from approximately 0.38 to 0.72, and automatic selection chose that fit. The noiseless rows contain one observation per condition and therefore describe individual fits rather than an error distribution.

##### CPI dispersion recovery and automatic selection

##### Supplementary Table S4-1 | CPI CBF recovery with dispersion.

| Simulated dispersion | SNR | Fit without dispersion | Fit with gamma dispersion | Automatic |
|---|---|---|---|---|
| None | Noiseless | 0.136 / 0.136 | 0.138 / 0.138 | 0.136 / 0.136 |
| Gamma | Noiseless | 0.379 / 0.379 | 0.721 / 0.721 | 0.721 / 0.721 |

| Simulated dispersion | SNR | Fit without dispersion | Fit with gamma dispersion | Automatic |
|---|---|---|---|---|
| Lognormal | Noiseless | 0.336 / 0.336 | 0.449 / 0.449 | 0.449 / 0.449 |
| None | 50 | 0.140 / 0.220 | 0.236 / 0.371 | 0.189 / 0.371 |
| Gamma | 50 | 0.278 / 0.466 | 0.070 / 0.229 | 0.073 / 0.246 |
| Lognormal | 50 | 0.197 / 0.436 | 0.101 / 0.196 | 0.109 / 0.217 |
| None | 20 | 0.198 / 0.436 | 0.276 / 0.509 | 0.230 / 0.436 |
| Gamma | 20 | 0.203 / 0.441 | 0.096 / 0.354 | 0.085 / 0.354 |
| Lognormal | 20 | 0.097 / 0.398 | 0.082 / 0.314 | 0.083 / 0.318 |

**Supplementary Table S4-2 | CPI MTT recovery with dispersion.**

| Simulated dispersion | SNR | Fit without dispersion | Fit with gamma dispersion | Automatic |
|---|---|---|---|---|
| None | Noiseless | 0.134 / 0.134 | 0.136 / 0.136 | 0.134 / 0.134 |
| Gamma | Noiseless | 0.385 / 0.385 | 0.721 / 0.721 | 0.721 / 0.721 |
| Lognormal | Noiseless | 0.342 / 0.342 | 0.449 / 0.449 | 0.449 / 0.449 |
| None | 50 | 0.083 / 0.197 | 0.183 / 0.328 | 0.142 / 0.328 |
| Gamma | 50 | 0.371 / 0.523 | 0.101 / 0.220 | 0.104 / 0.230 |
| Lognormal | 50 | 0.257 / 0.493 | 0.083 / 0.224 | 0.105 / 0.264 |
| None | 20 | 0.152 / 0.301 | 0.186 / 0.373 | 0.140 / 0.301 |
| Gamma | 20 | 0.331 / 0.610 | 0.112 / 0.540 | 0.121 / 0.540 |
| Lognormal | 20 | 0.196 / 0.532 | 0.086 / 0.182 | 0.099 / 0.225 |

Errors are median / 95th-percentile absolute log error. MTT and CBV truth were computed from the undispersed residue. All 123 observations produced accepted CPI maps; none used oSVD fallback or lacked a saved value.

Fitted-residue CBV changed little between the no-dispersion, gamma-dispersion, and automatic fits. At SNR 50, median absolute log error ranged from 0.050 to 0.059 and the 95th percentile from 0.082 to 0.104; at SNR 20, the corresponding ranges were 0.093–0.143 and 0.204–0.224.

**Supplementary Table S4-3 | Automatic CPI dispersion selection.**

| Simulated dispersion | SNR | No dispersion selected | Gamma dispersion selected |
|---|---|---|---|
| None | Noiseless | 1 | 0 |
| Gamma | Noiseless | 0 | 1 |
| Lognormal | Noiseless | 0 | 1 |
| None | 50 | 15 | 5 |
| Gamma | 50 | 3 | 17 |

| Simulated dispersion | SNR | No dispersion selected | Gamma dispersion selected |
|---|---|---|---|
| Lognormal | 50 | 7 | 13 |
| None | 20 | 17 | 3 |
| Gamma | 20 | 6 | 14 |
| Lognormal | 20 | 9 | 11 |

The final two columns give the number of observations assigned to each fit by the automatic dispersion comparison.

#### E. DSC population-DRO validation

The Semmineh population DRO [43,44] holds tissue physiology fixed while acquisition settings vary across 360 simulated scans.

**Semmineh acquisition-grid recovery**

**Supplementary Table S5a-1 | Reported outputs in the Semmineh acquisition grid.**

| Method | nCBF bias (%) | Reported tumor MTT bias (%) | Reported NAWM MTT bias (%) |
|---|---|---|---|
| sSVD | -42.3 [-44.9, -39.6] | +101.4 [+91.9, +111.1] | +8.4 [+4.6, +15.4] |
| oSVD | -34.3 [-37.3, -31.3] | +34.7 [+25.7, +44.1] | -16.4 [-21.9, -10.2] |
| Tikhonov | -30.1 [-33.1, -23.9] | +19.9 [+12.6, +28.5] | -20.3 [-25.8, -12.9] |
| CPI | -1.1 [-19.3, +19.7] | -7.8 [-20.9, +11.9] | -11.7 [-19.1, -7.4] |

**Supplementary Table S5a-2 | Residue-integral MTT in the Semmineh acquisition grid.**

| Method | Tumor residue-MTT bias (%) | NAWM residue-MTT bias (%) | Finite tumor / NAWM estimates |
|---|---|---|---|
| sSVD | +82.8 [+73.4, +95.8] | -1.5 [-6.6, +5.7] | 360/360; 360/360 |
| oSVD | +30.1 [+22.0, +38.1] | -20.0 [-25.0, -14.0] | 360/360; 360/360 |
| Tikhonov | +2.0 [-33.6, +55.4] | -49.4 [-68.9, -35.5] | 312/360; 355/360 |
| CPI | -7.8 [-20.9, +11.9] | -11.7 [-19.1, -7.4] | 360/360; 360/360 |

For each simulated scan, signal was averaged within each supplied region of interest (ROI), converted once to concentration, and corrected with BSW. The three SVD methods used the same corrected curves, whereas CPI fitted the original magnitude signal with the same AIF and a BSW-informed K2 prior. For SVD methods, reported MTT is AUC-CBV/CBF, whereas residue-integral MTT is the integral of the deconvolved residue divided by CBF. CPI reports its fitted-residue MTT, so its values are identical in the two tables. Values are median [Q1, Q3] percentage bias across the finite estimates from 360 scans. Tikhonov residue MTT was unavailable in 48 tumor and 5 NAWM fits;

the other endpoints were finite throughout. BSW-corrected non-leaky tumor nCBV bias was +7.0% [+2.9%, +10.9%]. CPI produced accepted estimates in all 720 tumor and NAWM fits: 685 met the solver convergence criterion, and 35 did not but passed PMRIPrep's CPI fit-acceptance checks. No oSVD fallback was used. This experiment measures sensitivity to acquisition settings around fixed physiology.

**Supplementary Table S5b | Semmineh nCBV leakage-correction sensitivity.**

| Region and correction | Median [Q1, Q3] bias (%) | 5th–95th percentile bias (%) |
|---|---|---|
| non-leaky, uncorrected | -7.9 [-9.0, -6.0] | [-10.3, -4.2] |
| leaky, uncorrected | -10.1 [-103.6, +32.6] | [-416.1, +55.4] |
| leaky, BSW-K2 corrected | -22.3 [-40.6, -14.9] | [-96.9, -11.4] |

The analysis compared separate leaky and non-leaky tumor regions generated from the same tumor hemodynamic specification (mean CBV 6.1%, SD 2.8%) across 360 acquisition settings. nCBV was calculated as the area under the tumor concentration curve divided by the corresponding area under the supplied NAWM reference curve; the ratio of the specified mean tumor and NAWM CBV values was 1.74. BSW correction narrowed the broad uncorrected leakage distribution but did not restore quantitative nCBV accuracy. All rows include 360 acquisition settings.

The nCBV bias reported with S5a refers to the BSW-corrected non-leaky tumor-to-NAWM AUC ratio. S5a averages the magnitude signal within each ROI before concentration conversion, matching its regional fitting analysis. S5b instead uses the supplied voxelwise concentration field to compare uncorrected non-leaky tumor and corrected leaky tumor nCBV. The approximately 15-percentage-point difference between the S5a and S5b non-leaky estimates therefore reflects both the processing route and the correction policy: S5a applies BSW correction, whereas the non-leaky estimate in S5b is uncorrected.

#### F. QIBA v9 Standard-Tofts recovery

PMRIPrep processed the complete Quantitative Imaging Biomarkers Alliance (QIBA) v9 Standard-Tofts digital reference object [45,46], which contains 44 acquisition conditions and 30 combinations of $K^{trans}$ and $v_e$. The primary analysis used its supplied noiseless AIF. For DRO signal-scale settings S0 = 5,000 and 10,000, median $K^{trans}$ and $v_e$ biases were within 2% at 6-s temporal resolution. At 10-s temporal resolution, the timing of frames relative to the arterial peak shifted median $K^{trans}$ bias to approximately −4% to +6%, while median $v_e$ bias remained within 2.5%. Here, temporal resolution is distinct from the 5-ms RF-excitation TR used in the spoiled-GRE signal equation.

**Supplementary Table S6 | QIBA v9 Standard-Tofts recovery.**

| Δt (s) | Sampling offset (s) | DRO signal scale (S0) | Supplied AIF peak (mM) | $K^{trans}$ bias (%) | $v_e$ bias (%) |
|---|---|---|---|---|---|
| 6 | 0 | 5,000 | 9.11 | -0.7 [-1.7, +0.3] | -1.0 [-1.9, -0.4] |
| 6 | 3 | 5,000 | 9.39 | -0.3 [-1.3, +1.6] | -0.9 [-1.8, -0.4] |
| 6 | 0 | 10,000 | 9.12 | -0.5 [-0.9, +0.2] | -0.4 [-0.6, -0.2] |
| 6 | 3 | 10,000 | 9.39 | -0.2 [-0.6, +1.0] | -0.5 [-0.7, -0.4] |
| 10 | 0 | 5,000 | 7.51 | +5.1 [+2.6, +13.0] | -0.4 [-2.3, +0.5] |
| 10 | 5 | 5,000 | 9.39 | -3.7 [-4.2, -3.2] | -2.4 [-3.3, -1.1] |
| 10 | 0 | 10,000 | 7.51 | +5.7 [+3.5, +12.8] | -0.1 [-1.6, +0.9] |
| 10 | 5 | 10,000 | 9.39 | -3.4 [-3.8, -3.0] | -1.9 [-2.6, 0] |

Values are median [Q1, Q3] percentage bias across the 30 combinations of $K^{trans}$ and $v_e$. S0 denotes the DRO signal-scale parameter in arbitrary units, distinct from the measured precontrast signal. The table reports noise-free conditions with S0 = 5,000 or 10,000. Both sampling offsets are retained as they change which frame samples the AIF peak.

The DRO also includes lower signal-scale and added-noise conditions. In the eight noise-free conditions with S0 = 500 or 1,000, median $K^{trans}$ bias ranged from −17.1% to −5.4% and median $v_e$ bias from −16.3% to −9.4%. The 28 noisy conditions had baseline SNRs of only 1.3–2.1; with the supplied AIF, $K^{trans}$ bias ranged from −97.0% to −17.0% and the within-patch $K^{trans}$ interquartile range divided by the median ranged from 99% to 264%.

#### G. QIBA Tofts-family recovery

The QIBA v4 object was used to evaluate Extended-Tofts recovery, including $v_p$, under noise-free conditions [45,46]. Matched QIBA v9 normal-output and v14 reduced-output objects were used to evaluate Standard-Tofts recovery with their supplied AIFs.

**Supplementary Table S7-1 | QIBA v4 Extended-Tofts recovery.**

| Patches | Plasma AIF peak (mM) | $K^{trans}$ MAE ($min^{-1}$) | $v_e$ MAE | $v_p$ MAE |
|---|---|---|---|---|
| 108 | 9.823 | 0.00183 | 0.03187 | 0.00071 |

MAE is the mean absolute error across the 108 tissue patches.

PMRIPrep signal-to-concentration conversion was applied to the noise-free QIBA v4 object using its supplied AIF without amplitude adjustment, and each patch-mean concentration curve was fitted with the standalone bounded Extended-Tofts fitter, with delay fixed at zero. These results verify recovery under the conditions represented by this object; they do not establish that Extended Tofts is valid for every tissue or acquisition. The v4 analysis used a temporal resolution of 0.5 s.

**Supplementary Table S7-2 | Standard-Tofts recovery under normal and reduced cardiac output.**

| Object | Supplied plasma AIF peak (mM) | $K^{trans}$ MAE ($min^{-1}$) | $v_e$ MAE |
|---|---|---|---|
| QIBA v9 normal output | 9.952 | 0.01084 | 0.07293 |
| QIBA v14 reduced output | 10.970 | 0.01234 | 0.03928 |

The matched, noise-free objects at S0 = 1,000 were evaluated in the 24 patches with $v_e \geq 0.05$, whose kinetics were resolvable at 6-s temporal resolution. Each patch-mean concentration curve was fitted with the standalone bounded Standard-Tofts fitter, with delay fixed at zero. This comparison tests recovery with the supplied AIF under two cardiac-output conditions; it does not treat AIF peak height as a cardiac-output measurement.

#### H. DCE cross-model recovery

Internally generated concentration-time curves tested estimator recovery, model mismatch, and model selection; they test implementation behavior but do not provide independent external validation.

Experiments used the Parker population AIF [12], scaled to a 5.0-mM peak with arrival at 0.5 min. The principal Tofts-family grid comprised 80 samples at 4-s intervals. Ground-truth curves were evaluated analytically on a time grid 100 times finer and then sampled at the acquisition times; fitting used only the sampled data. This prevented the same coarse numerical approximation from being used to both generate and fit the data. Direct 2CXM recovery experiments used 160 samples at 2-s intervals, preserving the 320-s acquisition duration. A separate model-selection experiment sampled 2CXM-generated curves at 4-s intervals, where production selection compared Patlak with Extended Tofts.

The Tofts grid used the six QIBA v9 $K^{trans}$ values (0.01, 0.02, 0.05, 0.10, 0.20, and 0.35 $min^{-1}$) crossed with five $v_e$ values (0.01, 0.05, 0.10, 0.20, and 0.50). Tofts, Extended Tofts, and Patlak were fitted to these Tofts-generated curves to separate matched-model recovery from the effects of adding a vascular term or neglecting backflux. Separate Extended-Tofts curves covered $v_p$ = 0.01–0.08 under two extravascular conditions. Four physiological 2CXM conditions covered normal gray matter and tumorlike plasma flow ($F_p$ = 50–80 mL/100 mL/min), permeability–surface area product (PS = 4–30 mL/100 mL/min), $v_e$ = 0.15–0.35, and $v_p$ = 0.03–0.06. Model-selection experiments generated Patlak, Extended Tofts, and 2CXM curves independently at 2-, 2.5-, or 4-s temporal resolution and fitted every model eligible at the corresponding interval. Section J examines 2CXM plasma-flow recovery across temporal resolutions of 1–7.5 s.

Additive zero-mean Gaussian concentration noise was scaled to each curve's peak. The primary heatmaps used 0% and 5% noise; noise-response experiments sampled 0%, 1%, 2%, 3%, 5%, 7%, and 10%. Each noisy parameter combination used 200 Monte Carlo realizations and is summarized by its median and first and third quartiles. The zero-noise conditions are deterministic. For delay estimation, the 200 realizations of each combination formed a homogeneous 20×10 voxel plane at 1-mm spacing; zero-noise curves were fitted alone.

Patlak, Tofts, Extended Tofts, and 2CXM were evaluated with the same BLS fitting, effective-delay estimation and fit-acceptance criteria used by PMRIPrep. The S8 tables summarize the experiments; Figure 3a shows recovery across the Tofts grid.

**Supplementary Table S8a | Tofts-family cross-model recovery.**

| Fitted model | Noise | $K^{trans}$ bias, median / maximum (%) | $v_e$ error, median / maximum | Accepted fits |
|---|---|---|---|---|
| Tofts | 0% | 0.5 / 36.5 | 0.001 / 0.003 | 30/30 |
| Tofts | 5% | 0.3 / 34.8 | 0.001 / 0.056 | 5999/6000 |
| ET | 0% | 2.2 / 46.1 | 0.002 / 0.009 | 30/30 |
| ET | 5% | 2.0 / 79.0 | 0.004 / 0.024 | 5997/6000 |
| Patlak | 0% | 77.2 / 100.0 | — | 30/30 |
| Patlak | 5% | 77.2 / 100.0 | — | 6000/6000 |

For Tofts and Extended Tofts (ET), the data were generated from the Tofts model and therefore test matched or nested recovery. Patlak was fitted to the same reversible curves to quantify the consequence of neglecting backflux. For $K^{trans}$, the table gives the median and maximum of the absolute median percentage biases across the 30 parameter combinations. The $v_e$ columns give the corresponding median and maximum absolute errors.

At 5% noise, 24 of 30 Tofts cells had median $K^{trans}$ bias within ±0.5%. In the two fastest-washout cells ($k_{ep}$ = 20 and 35 $min^{-1}$), delay estimation gave median biases of +11.8% and +34.8%, with fitted delays of 0.3 and 0.5 s. Fixing delay at its true value of zero reduced these biases to +1.2% and −1.9%. Extended Tofts returned $v_p \leq 0.007$ but increasingly underestimated $K^{trans}$ at faster kinetics, by up to 79% at 5% noise, where its additional parameter was weakly identifiable. Patlak's large negative bias on these reversible curves is expected, as Patlak neglects backflux.

#### Physiological 2CXM recovery at 2-s temporal resolution

##### Supplementary Table S8b-1 | Ground truth for physiological 2CXM recovery at 2-s temporal resolution.

| Tissue condition | $F_p$ | PS | $v_e$ | $v_p$ | $K^{trans}$ ($min^{-1}$) |
|---|---|---|---|---|---|
| Gray matter | 50 | 4 | 0.15 | 0.03 | 0.0370 |
| Tumor | 60 | 12 | 0.3 | 0.04 | 0.1000 |
| High-flow tumor | 80 | 20 | 0.35 | 0.05 | 0.1600 |
| Tumor rim | 70 | 30 | 0.15 | 0.06 | 0.2100 |

##### Supplementary Table S8b-2 | Physiological 2CXM recovery at 2-s temporal resolution.

| Tissue condition | $K^{trans}$ bias (%) | PS bias (%) | $\Delta v_e$ | $\Delta v_p$ |
|---|---|---|---|---|
| Gray matter | -0.5 / -0.3 [-3,+2] | -0.5 / -0.3 | -0.001 [-0.005,+0.004] | 0 |
| Tumor | -0.5 / -0.3 [-2,+2] | -0.5 / -0.8 | -0.001 [-0.005,+0.002] | -0.001 |
| High-flow tumor | -0.5 / -0.2 [-2,+2] | -0.5 / -1.2 | -0.001 [-0.004,+0.002] | -0.001 |
| Tumor rim | -0.4 / +2.0 [-3,+6] | -0.6 / -0.5 | +0.002 [-0.004,+0.008] | -0.004 |

$F_p$ and PS are reported in mL/100 mL/min. The physiological 2CXM conditions were generated and fitted with 2CXM; all 800 fits at 5% noise were accepted. Bias entries report the noiseless result followed by the median at 5% noise. For $K^{trans}$ and $\Delta v_e$, brackets give the first and third quartiles at 5% noise. The $\Delta v_e$ and $\Delta v_p$ columns report signed differences. At 5% simulated concentration noise, median $K^{trans}$ and PS biases were within 2.0% in magnitude. A separate rapid-exchange condition, reported below, tested a less identifiable setting: Tofts fits remained stable and Extended Tofts $K^{trans}$ bias reached −13%, whereas 2CXM developed large errors and up to 25% rejected fits as noise increased.

##### Supplementary Table S8c | Noise sensitivity with adequately sampled kinetics ($k_{ep}$ = 1.43 $min^{-1}$; Tofts $K^{trans}$ = 0.2, $v_e$ = 0.14; 2CXM $F_p$/PS/$v_e$/$v_p$ = 60/30/0.14/0.03).

| Noise | Tofts | ET | 2CXM |
|---|---|---|---|
| 0% | -0.3 | -0.4 | -0.5 |
| 1% | -0.3 [-0.5,-0.1] | -0.8 [-1.3,-0.5] | +0.1 [-0.8,+0.8] |
| 2% | -0.2 [-0.6,+0.1] | -1.0 [-1.8,-0.5] | +0.5 [-1.4,+1.9] |
| 3% | -0.2 [-0.7,+0.4] | -1.2 [-2.3,-0.4] | +1.0 [-1.9,+2.8] |

| Noise | Tofts | ET | 2CXM |
|---|---|---|---|
| 5% | -0.1 [-1.0,+0.9] | -1.6 [-3.1,-0.2] | +1.9 [-3.0,+5.1] |
| 7% | +0.1 [-1.2,+1.3] | -1.9 [-4.0,-0.1] | +1.3 [-4.3,+6.1] |
| 10% | +0.3 [-1.5,+2.1] | -2.3 [-5.1,+0.3] | +0.3 [-8.2,+6.9] |

Values are $K^{trans}$ percentage errors, reported as median [Q1, Q3] across 200 realizations except for the deterministic 0% noise condition. ET, Extended Tofts.

**Supplementary Table S8d | Noise sensitivity with rapid exchange ($k_{ep}$ = 7 min⁻¹; Tofts $K^{trans}$ = 0.35, $v_e$ = 0.05; 2CXM $F_p$/PS/$v_e$/$v_p$ = 70/70/0.05/0.05).**

| Noise | Tofts median [Q1,Q3] | ET median [Q1,Q3] | 2CXM median | 2CXM accepted (%) | 2CXM rejected or >50% error (%) | 2CXM 5th–95th percentile |
|---|---|---|---|---|---|---|
| 0% | -0.2 | -0.2 | -2.2 | 100% | 0% | [-2, -2] |
| 1% | -0.2 [-0.8,+0.4] | -3.4 [-5.2,-1.5] | +8.8 | 100% | 1% | [-33, +35] |
| 2% | -0.2 [-1.4,+1.0] | -4.7 [-7.7,-2.1] | +10.9 | 86% | 25% | [-74, +37] |
| 3% | -0.2 [-1.9,+1.8] | -6.0 [-9.7,-2.6] | +5.4 | 85% | 34% | [-91, +44] |
| 5% | -0.2 [-2.8,+3.0] | -8.1 [-13.2,-2.4] | -12.5 | 78% | 48% | [-100, +40] |
| 7% | -0.1 [-3.8,+4.2] | -9.8 [-17.2,-2.2] | -26.2 | 79% | 53% | [-100, +50] |
| 10% | 0 [-5.5,+6.6] | -13.1 [-21.8,-2.3] | -42.3 | 75% | 60% | [-100, +41] |

All error columns report $K^{trans}$ percentage error. Central values are medians; the final column summarizes the accepted 2CXM fits. The wide 2CXM percentile ranges and failure percentages show the asymmetric high-error distributions.

#### I. OSIPI external DCE validation

PMRIPrep outputs were evaluated using the OSIPI-DCE Challenge definitions [47]. Accuracy was assessed in four synthetic sessions representing two synthetic subjects (P1 and P2) and two visits, and repeatability was assessed in eight clinical test–retest subjects. The synthetic tissue curves were generated with Extended Tofts. Scoring used the challenge-provided tumor ROIs. The primary analysis included every finite voxel in each challenge tumor ROI, matching the published challenge evaluation; a secondary analysis included only voxels passing the DCE QC criteria. Accuracy compares each synthetic-session ROI mean with its reference value, whereas repeatability compares the two clinical visit means. The OSIPI silver score is 100 times the product of the accuracy score and the $K^{trans}$ repeatability score.

*Synthetic accuracy and hematocrit-convention sensitivity*

**Supplementary Table S9a | OSIPI DCE accuracy and repeatability scores.**

| AIF hematocrit convention | Voxel set | Accuracy score | $K^{trans}$ repeatability score | OSIPI silver score |
|---|---|---|---|---|
| Default (Hct = 0.42) | All finite tumor-ROI voxels | 0.847 | 0.850 | 72.0 |
| Default (Hct = 0.42) | QC-passing tumor-ROI voxels | 0.861 | 0.892 | 76.8 |
| Hct = 0 sensitivity | All finite tumor-ROI voxels | 0.923 | NA | NA |
| Hct = 0 sensitivity | QC-passing tumor-ROI voxels | 0.907 | NA | NA |

The hematocrit-zero analysis included only the four synthetic sessions and therefore has no clinical repeatability or combined silver score. The QC-passing rows use each run's own DCE QC mask. The all-finite-voxel repeatability score fell within the range reported for the ten challenge submissions (0.64–0.86), and the accuracy score within the published range (0.54–0.92). Only scores using all finite tumor-ROI voxels are directly comparable with the published challenge submissions.

**Supplementary Table S9b | OSIPI synthetic-session accuracy using all finite tumor-ROI voxels.**

| Synthetic session | GT mean $K^{trans}$ | Recovered mean | Bias | N voxels | MAE | Pearson r |
|---|---:|---:|---:|---:|---:|---:|
| P1, visit 1 | 0.0111 | 0.0087 | −0.0024 | 298 | 0.0038 | 0.770 |
| P1, visit 2 | 0.0041 | 0.0032 | −0.0010 | 341 | 0.0017 | 0.843 |
| P2, visit 1 | 0.1024 | 0.0684 | −0.0340 | 13,511 | 0.0404 | 0.759 |
| P2, visit 2 | 0.0686 | 0.0447 | −0.0240 | 11,769 | 0.0264 | 0.864 |

$K^{trans}$ values, bias and MAE are in $min^{-1}$; bias is the recovered minus the ground-truth (GT) ROI mean. Bias was negative in all four sessions.

**Hematocrit convention.** For a measured AIF, PMRIPrep treats the arterial curve as whole-blood concentration and divides it by (1 − Hct), with Hct = 0.42 by default, to obtain plasma concentration. The OSIPI synthetic objects were generated from an AIF already expressed as plasma concentration (corrected with hematocrit 0.45 [47]); for these objects, the generating convention corresponds to Hct = 0 in PMRIPrep's arterial conversion. The default correction therefore enlarges the AIF relative to the generating convention and lowers $K^{trans}$. The Hct = 0 analysis tests this convention difference. Clinical arterial signal reflects whole blood, so the default convention remains the primary analysis.

**Supplementary Table S9c | Hematocrit sensitivity in shared QC-passing voxels.**

| Session | N voxels | GT | Default (Hct = 0.42) | Hct = 0 |
|---|---|---|---|---|
| P1, visit 1 | 228 | 0.0109 | 0.0088 | 0.0152 |
| P1, visit 2 | 204 | 0.0046 | 0.0037 | 0.0063 |
| P2, visit 1 | 7,415 | 0.0842 | 0.0583 | 0.0941 |
| P2, visit 2 | 6,162 | 0.0585 | 0.0375 | 0.0611 |

Values are ROI-mean $K^{trans}$ ($min^{-1}$) on the intersection of the default and Hct = 0 DCE QC masks, so both conventions are compared over identical voxels. Setting Hct = 0 moved both P2 estimates close to ground truth. The aggregate accuracy score increased, from 0.857 to 0.905 on shared QC-passing voxels and from 0.847 to 0.923 on all finite voxels. Hct = 0 is reported as a sensitivity analysis of AIF convention rather than as an alternative preferred result.

*Clinical test–retest repeatability*

The OSIPI-DCE Challenge clinical test–retest dataset comprised eight subjects with recurrent glioblastoma selected from the RIDER Neuro MRI collection and scanned twice on the same 1.5 T Siemens scanner, two days apart [47], with a temporal resolution of approximately 4.76 s. Processing used pipeline defaults: automatic AIF detection with population fallback, automatic BIC-based model selection (Patlak and Extended Tofts at this temporal resolution), effective-delay estimation, and BLS fitting. The OSIPI silver score uses $K^{trans}$ repeatability; $v_e$ and $v_p$ are reported as additional repeatability endpoints.

**Supplementary Table S9d | OSIPI clinical parameter repeatability using all finite tumor-ROI voxels.**

| Parameter | Score | Challenge wCV | Challenge %RC | Conventional wCV | Conventional %RC |
|---|---|---|---|---|---|
| $K^{trans}$ | 0.850 | 18.2% | 50.4% [36.0%, 62.3%] | 25.7% | 71.2% [50.8%, 88.1%] |
| $v_e$ | 0.866 | 16.6% | 46.1% [27.1%, 61.5%] | 23.5% | 65.1% [38.3%, 86.9%] |
| $v_p$ | 0.891 | 13.0% | 35.9% [23.0%, 48.8%] | 18.3% | 50.8% [32.5%, 69.0%] |

Score is the OSIPI repeatability score; wCV is the root-mean-square within-subject CV, and %RC is the repeatability coefficient (2.77 × wCV). The challenge calculation uses the population standard deviation of each two-visit pair; the conventional calculation uses the sample standard deviation. Brackets give 95% intervals from 10,000 subject-level bootstrap samples. With QC-passing voxels, $K^{trans}$ challenge %RC was 34.4% [24.7%, 42.4%] and conventional %RC was 48.7% [34.9%, 59.9%].

**Supplementary Table S9e | OSIPI clinical $K^{trans}$ test–retest results.**

| Subject | Visit 1 | Visit 2 | CV (%) | AIF, both visits |
|---|---|---|---|---|
| C1 | 0.0312 | 0.0550 | 39.0 | population |
| C2 | 0.0278 | 0.0202 | 22.5 | detected |
| C3 | 0.0168 | 0.0252 | 28.3 | detected |
| C4 | 0.0165 | 0.0140 | 11.9 | detected |
| C5 | 0.0184 | 0.0230 | 15.5 | detected |
| C6 | 0.0344 | 0.0212 | 33.4 | population |
| C7 | 0.0165 | 0.0257 | 30.9 | detected |
| C8 | 0.0194 | 0.0187 | 2.9 | detected |

Values are tumor-ROI mean $K^{trans}$ ($min^{-1}$) using all finite voxels. CV uses the sample standard deviation of the two visits.

**AIF consistency.** The AIF source matched between visits for all eight subjects. C1 and C6 used the Parker population AIF on both visits; the remaining six used automatically detected AIFs.

**Repeatability.** The median subject-level CV of tumor-ROI mean $K^{trans}$ was 25.4%, and the root-mean-square within-subject CV was 25.7% (18.2% with the challenge convention). The two subjects using the population AIF had the largest CVs (C1, 39.0%; C6, 33.4%); this descriptive association was not tested as a causal effect of AIF selection. Bland–Altman analysis of tumor-ROI mean $K^{trans}$ between visits (visit 1 minus visit 2, n = 8) gave a bias of −0.00274 $min^{-1}$ and limits of agreement of −0.0252 to +0.0197 $min^{-1}$.

Limits of agreement were calculated as the mean paired difference ±1.96 times the sample standard deviation of the paired differences.

#### J. 2CXM plasma-flow recovery

**Flow recovery by temporal resolution.** Internal simulations generated 2CXM curves without AIF–tissue delay over 5 minutes at temporal resolutions of 1–7.5 s with four bolus timings relative to the sampling grid, for eight conditions chosen to evaluate flow recovery, including normal-brain values [40]. Curves were computed on a fine grid using the Parker AIF [12] (first-pass full width at half maximum 8.4 s) or the same AIF dispersed by a unit-area exponential kernel (time constant 5.9 s; full width at half maximum 14 s). The fitted AIF had independent Gaussian noise with SD 1% of its peak. Tissue signal had Rician noise with the same SD in all conditions, chosen to give a concentration CNR of 15.3 for an intermediate-flow condition at 2-s temporal resolution [39]. Each condition and bolus timing had 200 realizations, arranged as 2×5 voxel patches at 1-mm spacing with the same tissue parameters, so that delay estimation pooled neighboring voxels as in PMRIPrep. Direct 2CXM fits used effective-delay estimation.

**Supplementary Table S10a | Plasma flow ($F_p$) recovery across temporal resolutions.**

| Δt (s) | 8.4-s AIF | 14-s AIF |
|---|---|---|
| 1 | +2 / 19 (6,400) | +5 / 28 (6,400) |
| 2 | +7 / 34 (6,400) | +7 / 52 (6,398) |
| 2.5 | +7 / 44 (6,399) | +9 / 77 (6,398) |
| 3 | +7 / 56 (6,400) | +13 / 113 (6,385) |
| 4 | +21 / 144 (6,397) | +24 / 152 (6,386) |
| 6 | +328 / 653 (6,176) | +85 / 444 (6,269) |
| 7.5 | +532 / 1014 (5,603) | +228 / 906 (5,905) |

Cells give the largest absolute median bias (sign retained) / largest 95th-percentile absolute error (%), over conditions and bolus timings. Results use explicit-2CXM estimates retained in production parameter maps, before QC masking; parentheses give the number retained from 6,400 attempted fits. With absolute bias ≤20% and 95th-percentile error ≤50% as limits, flow met these criteria through 1 s with the broader AIF and through 2.5 s with the Parker AIF (Table S10a).

Fitted to these 2CXM-generated curves, Extended Tofts overestimated $K^{trans}$ and underestimated $v_p$ when plasma transit was appreciable, consistent with its omission of plasma transit [37]; this comparison favors 2CXM by construction. In slow-backflux tissue, $v_e$ was unreliable with either model over the 5-minute acquisition [48]: Extended Tofts returned consistently low values, whereas many 2CXM estimates reached the $v_e$ + $v_p$ = 1 limit.

**Flow recovery with AIF–tissue delay.** The simulations were repeated at 2-, 4- and 7.5-s intervals with the tissue curves delayed by 2 or 5 s relative to the AIF (Table S10b).

**Supplementary Table S10b | Plasma flow ($F_p$) recovery with AIF–tissue delay.**

| Δt (s) | Delay (s) | 8.4-s AIF | 14-s AIF |
|---|---|---|---|
| 2 | 0 | +7 / 34 (6,400) | +7 / 52 (6,398) |
| 2 | 2 | +5 / 33 (6,400) | +8 / 57 (6,399) |

| Δt (s) | Delay (s) | 8.4-s AIF | 14-s AIF |
|---|---|---|---|
| 2 | 5 | +3 / 29 (6,400) | +7 / 70 (6,396) |
| 4 | 0 | +21 / 144 (6,397) | +24 / 152 (6,386) |
| 4 | 2 | +12 / 63 (6,400) | +12 / 128 (6,379) |
| 4 | 5 | +15 / 81 (6,390) | +15 / 149 (6,383) |
| 7.5 | 0 | +532 / 1014 (5,603) | +228 / 906 (5,905) |
| 7.5 | 2 | +379 / 981 (5,670) | +200 / 749 (6,004) |
| 7.5 | 5 | +148 / 448 (6,280) | +73 / 506 (6,197) |

Cells, retained counts and inclusion follow Table S10a; zero-delay rows repeat S10a.

With AIF–tissue delay, the Parker AIF still met both limits at 2 s, whereas 95th-percentile errors with the broader AIF rose from 52% without delay to 57% and 70% with 2- and 5-s delays. To separate delay-estimation error from sampling error, the fits were repeated with the delay fixed at its simulated value. At 7.5 s, the largest 95th-percentile errors fell from 448–981% (Table S10b) to 99–269%, but still exceeded the criteria for both AIFs and both delays. Rapid sampling is recommended for plasma-flow estimation [39,49].

The simulations assumed 2CXM tissue kinetics, a 5-minute scan duration, and independent AIF noise; noise level, scan duration, and tissue kinetics also affect recovery [48].

#### K. Effective-delay estimation

Effective-delay estimation was compared with a fixed zero delay for Extended Tofts tissue with no delay or a 5-s delay at 4- and 7.4-s temporal resolution. Estimation reduced error for the 5-s delay in most conditions but could fit a nonzero delay where none was present.

**Supplementary Table S11 | Effect of effective-delay estimation on $K^{trans}$ and $v_p$.**

| Δt (s) | Condition | Delay (s) | $K^{trans}$ MAE, fixed zero | $K^{trans}$ MAE, estimated | $v_p$ MAE, fixed zero | $v_p$ MAE, estimated | Median fitted delay (s) |
|---|---|---|---|---|---|---|---|
| 4 | Moderate | 0 | 0.0021 | 0.0021 | 0.0011 | 0.0012 | 0 |
| 4 | Moderate | 5 | 0.0210 | 0.0021 | 0.0212 | 0.0012 | 5.1 |
| 4 | Fast washout | 0 | 0.0210 | 0.0231 | 0.0018 | 0.0029 | 0.2 |
| 4 | Fast washout | 5 | 0.0388 | 0.0200 | 0.0100 | 0.0024 | 5.2 |
| 7.4 | Moderate | 0 | 0.0026 | 0.0020 | 0.0015 | 0.0031 | 1.4 |
| 7.4 | Moderate | 5 | 0.0120 | 0.0044 | 0.0150 | 0.0079 | 5.1 |

| Δt (s) | Condition | Delay (s) | $K^{trans}$ MAE, fixed zero | $K^{trans}$ MAE, estimated | $v_p$ MAE, fixed zero | $v_p$ MAE, estimated | Median fitted delay (s) |
|---|---|---|---|---|---|---|---|
| 7.4 | Fast washout | 0 | 0.1059 | 0.0967 | 0.0078 | 0.0268 | 3.4 |
| 7.4 | Fast washout | 5 | 0.0157 | 0.0298 | 0.0100 | 0.0086 | 6.2 |

Extended Tofts tissue (moderate: $K^{trans}$ = 0.10 $min^{-1}$, $v_e$ = 0.30, $v_p$ = 0.03; fast washout: $K^{trans}$ = 0.35 $min^{-1}$, $v_e$ = 0.10, $v_p$ = 0.01) was computed on a fine grid with the Parker AIF [12], delayed and sampled over 80 frames. The fitted AIF was noise-free. Gaussian noise was 5% of the tissue peak, with 49 realizations per row arranged as 7×7 voxel in-plane patches at 1-mm spacing, and models were selected automatically. MAE ($K^{trans}$ in $min^{-1}$) used all finite retained estimates without QC-mask filtering.

In the moderate condition, estimation reduced $K^{trans}$ MAE for the 5-s delay from 0.0210 to 0.0021 $min^{-1}$ at 4 s and from 0.0120 to 0.0044 $min^{-1}$ at 7.4 s, and $v_p$ errors also decreased. Without delay, it fitted a median delay of 1.4 s at 7.4 s and doubled $v_p$ MAE. In the fast-washout condition at 7.4 s, the fitted delay reached 3.4 s without delay, and with a 5-s delay $K^{trans}$ MAE increased. Additional explicit Extended Tofts simulations with $v_p$ = 0–0.03, four sample timings relative to the AIF and delays of 0–3 s showed the same pattern: fitted delays could exceed the true delay, particularly at coarse temporal resolution. Estimation reduced $K^{trans}$ error for nonzero true delays at $v_p$ = 0.02–0.03 but increased it at 4 s for $v_p$ = 0 with delays up to 1 s, and at 7.4 s for $v_p \leq 0.01$ at most delays. Delay estimation therefore improves kinetic recovery when timing mismatch is substantial but is limited at coarse temporal resolution; the fitted delay is an effective model alignment rather than a measurement of vascular timing.

#### L. DSC-informed Bayesian $v_p$ prior

We tested whether a DSC-informed $v_p$ prior improves DCE recovery, how errors in the prior affect that benefit, and how much the prior changes clinical estimates. Simulations compared VI with each model's default priors and VI with the DSC-informed $v_p$ prior, both using the same BLS model, selected delay and starting estimates. The DSC-informed prior SD was set to 60% of its center. Comparisons included only retained tumor voxels with successful VI fits in both arms; BLS fallback in either arm excluded the voxel from both. Results are mean within-seed percentage RMSE changes across 20 seeds, with 95% confidence intervals from 10,000 paired bootstrap resamples. Negative changes indicate lower error.

*L.1 Recovery across models and temporal resolutions*

Concentration curves used temporal resolutions of 2, 4 or 7.5 s over 360 s, bolus arrival at 79.8 s, zero true effective delay and a known plasma AIF. Gaussian noise SD was 5% or 10% of each curve's peak. Each seed included 60 tumor targets on a 6×10 voxel plane at 1-mm spacing, with 100 NAWM values for prior normalization. Tumor and NAWM regions were separated so their delay-estimation neighborhoods did not overlap.

Extended-Tofts truth spanned $K^{trans}$ = 0.03–0.30 min⁻¹, $v_e$ = 0.15–0.55 and $v_p$ = 0.005–0.050. Patlak used $K^{trans}$ = 0.003–0.030 min⁻¹ and the same $v_p$ range. The 2CXM simulations used the same $K^{trans}$, $v_e$ and $v_p$ values as the Extended Tofts simulations, with plasma flow of 40–80 mL/min/100 mL and PS calculated to preserve $K^{trans}$. Each model fitted its own kinetics, so these experiments do not rank the models. All three models used the fitter's PCHIP operator for curve generation.

**Supplementary Table S12a | Default-prior VI recovery relative to least-squares fitting.**

| Model | Temporal resolution (s) | DCE noise | $v_p$ RMSE change [95% CI] | $K^{trans}$ RMSE change [95% CI] | $v_e$ RMSE change [95% CI] |
|---|---|---|---|---|---|
| ET | 2 | 5% | -0.4% [-1.8, +1.1] | 0% [-1.6, +1.6] | -0.6% [-13.4, +11.3] |
| ET | 2 | 10% | -1.8% [-3.8, +0.3] | -0.7% [-3.2, +1.8] | -6.9% [-20.5, +6.7] |
| ET | 4 | 5% | -0.1% [-1.6, +1.4] | -1.0% [-2.0, +0.01] | -6.9% [-20.8, +5.7] |
| ET | 4 | 10% | -3.2% [-5.2, -1.1] | -2.8% [-4.6, -0.9] | -19.9% [-33.1, -6.8] |
| ET | 7.5 | 5% | -0.4% [-2.0, +1.4] | -1.0% [-2.9, +0.8] | -30.1% [-45.7, -14.6] |
| ET | 7.5 | 10% | -4.3% [-6.4, -1.9] | -3.7% [-6.5, -0.8] | -38.9% [-49.9, -27.5] |
| Patlak | 2 | 5% | -0.6% [-1.9, +0.9] | +2.1% [+0.1, +4.5] | — |
| Patlak | 2 | 10% | +1.1% [-1.5, +3.6] | +7.8% [+4.5, +11.9] | — |
| Patlak | 4 | 5% | +1.7% [+0.3, +3.1] | +4.5% [+1.9, +7.3] | — |
| Patlak | 4 | 10% | +5.5% [+3.0, +7.9] | +12.3% [+8.1, +16.7] | — |

| Model | Temporal resolution (s) | DCE noise | $v_p$ RMSE change [95% CI] | $K^{trans}$ RMSE change [95% CI] | $v_e$ RMSE change [95% CI] |
|---|---|---|---|---|---|
| Patlak | 7.5 | 5% | +3.2% [+1.3, +5.2] | +8.9% [+5.7, +12.0] | — |
| Patlak | 7.5 | 10% | +9.8% [+6.4, +13.3] | +20.7% [+16.3, +25.1] | — |
| 2CXM, all paired | 2 | 5% | -68.6% [-75.7, -60.4] | -63.2% [-72.6, -52.1] | -59.2% [-67.7, -49.4] |
| 2CXM, all paired | 2 | 10% | -78.9% [-82.1, -75.3] | -72.0% [-76.0, -67.5] | -66.4% [-70.5, -62.0] |
| 2CXM, all paired | 4 | 5% | -69.7% [-77.4, -60.4] | -53.8% [-64.9, -41.8] | -58.1% [-67.0, -47.5] |
| 2CXM, all paired | 4 | 10% | -77.5% [-81.5, -72.7] | -63.1% [-69.2, -56.4] | -64.1% [-70.7, -55.8] |
| 2CXM, all paired | 7.5 | 5% | -74.9% [-79.0, -70.3] | -60.7% [-68.5, -51.5] | -64.8% [-70.3, -58.6] |
| 2CXM, all paired | 7.5 | 10% | -79.4% [-82.5, -75.4] | -57.6% [-65.3, -47.9] | -66.5% [-73.1, -57.4] |
| 2CXM, well constrained | 2 | 5% | -30.3% [-40.9, -20.3] | -27.1% [-39.4, -15.6] | -8.8% [-21.9, +3.9] |
| 2CXM, well constrained | 2 | 10% | -35.8% [-47.6, -24.0] | -23.2% [-37.4, -8.1] | -7.2% [-19.3, +4.5] |
| 2CXM, well constrained | 4 | 5% | -31.6% [-43.8, -20.5] | -15.6% [-31.6, +1.1] | -7.3% [-21.4, +6.4] |
| 2CXM, well constrained | 4 | 10% | -26.0% [-38.3, -13.3] | -13.8% [-24.2, -4.5] | -6.2% [-19.3, +9.5] |
| 2CXM, well constrained | 7.5 | 5% | -17.4% [-28.5, -5.2] | -5.2% [-15.1, +4.2] | -10.0% [-22.3, +3.9] |
| 2CXM, well constrained | 7.5 | 10% | -16.6% [-32.5, +1.4] | -10.5% [-18.2, -2.7] | +1.3% [-12.4, +16.5] |

Changes are relative to the shared BLS fit. Voxels are those used for the unbiased-prior comparisons in Tables S12b–e: 1,199–1,200 per condition for Extended Tofts, 1,200 for Patlak, 869–1,062 for all paired 2CXM fits and 296–658 for well-constrained 2CXM fits. Well-constrained fits are those whose primary parameters passed the identification criteria of the DCE QC mask (Supplementary Methods) at the shared BLS fit. ET, Extended Tofts.

Default-prior VI changed Extended-Tofts $K^{trans}$ and $v_p$ RMSE little and reduced $v_e$ RMSE at coarser temporal resolution, most at 7.5 s. For explicit 2CXM, VI reduced error in all three parameters; where the shared BLS fit was well constrained, reductions were smaller and no $v_e$ change was detected. For Patlak, VI increased $K^{trans}$ RMSE at every temporal resolution and noise level, and $v_p$ RMSE at 4 and 7.5 s.

The comparisons below measure the effect of replacing the default $v_p$ prior with the DSC-informed prior; all other priors were unchanged. Tables S12b–e use four prior-center scenarios:

- **True $v_p$:** the prior mean equals the voxel's ground-truth $v_p$.
- **Unbiased, 20% CV:** random multiplicative errors with no imposed systematic bias and a 20% coefficient of variation.
- **Tumor 30% low, 20% CV:** the same random errors, with tumor values scaled to 70% of truth.
- **Tumor 30% high, 20% CV:** the same random errors, with tumor values scaled to 130% of truth.

The random errors were lognormal. Noisy values were normalized by their NAWM median and multiplied by 0.010; true NAWM $v_p$ ranged from 0.008 to 0.012. The 20% CV describes errors in prior centers; all four conditions retained the same broad prior width.

**Supplementary Table S12b | Extended-Tofts prior sensitivity across temporal resolutions.**

| Temporal resolution (s) | DCE noise | Prior scenario | $v_p$ RMSE change [95% CI] | $K^{trans}$ RMSE change [95% CI] | $v_e$ RMSE change [95% CI] |
|---|---|---|---|---|---|
| 2 | 5% | True $v_p$ | -1.1% [-2.4, +0.2] | -0.2% [-1.5, +1.0] | -0.4% [-1.0, +0.2] |
| 2 | 5% | Unbiased, 20% CV | -1.1% [-2.4, +0.2] | 0% [-1.3, +1.2] | -0.4% [-1.0, +0.3] |
| 2 | 5% | Tumor 30% low, 20% CV | -0.7% [-2.4, +0.8] | +0.1% [-1.6, +1.6] | 0% [-0.7, +0.7] |
| 2 | 5% | Tumor 30% high, 20% CV | -0.7% [-1.8, +0.5] | +0.1% [-1.2, +1.2] | -0.6% [-1.2, +0.1] |
| 2 | 10% | True $v_p$ | -3.0% [-4.8, -1.2] | -0.7% [-2.7, +1.2] | -0.5% [-1.2, +0.3] |
| 2 | 10% | Unbiased, 20% CV | -2.9% [-4.7, -1.1] | -0.4% [-2.4, +1.5] | -0.4% [-1.2, +0.4] |
| 2 | 10% | Tumor 30% low, 20% CV | -2.2% [-4.2, -0.2] | -0.3% [-2.7, +2.0] | +0.2% [-0.7, +1.2] |
| 2 | 10% | Tumor 30% high, 20% CV | -1.8% [-3.7, +0.04] | +0.1% [-2.2, +2.1] | -0.8% [-1.5, -0.1] |

| Temporal resolution (s) | DCE noise | Prior scenario | $v_p$ RMSE change [95% CI] | $K^{trans}$ RMSE change [95% CI] | $v_e$ RMSE change [95% CI] |
|---|---|---|---|---|---|
| 4 | 5% | True $v_p$ | -1.7% [-3.1, -0.2] | +0.3% [-1.0, +1.7] | -0.6% [-1.5, +0.3] |
| 4 | 5% | Unbiased, 20% CV | -1.5% [-2.9, +0.1] | +0.5% [-1.0, +2.0] | -0.7% [-1.7, +0.3] |
| 4 | 5% | Tumor 30% low, 20% CV | -2.4% [-3.8, -1.0] | +0.6% [-0.8, +2.1] | -0.3% [-1.4, +0.9] |
| 4 | 5% | Tumor 30% high, 20% CV | 0% [-1.4, +1.5] | +0.9% [-0.5, +2.3] | -0.9% [-1.8, -0.1] |
| 4 | 10% | True $v_p$ | -3.2% [-5.5, -0.6] | +0.4% [-1.9, +2.6] | -0.8% [-1.6, +0.002] |
| 4 | 10% | Unbiased, 20% CV | -2.7% [-5.2, +0.1] | +0.6% [-1.9, +2.9] | -0.9% [-1.8, +0.04] |
| 4 | 10% | Tumor 30% low, 20% CV | -4.5% [-6.5, -2.4] | +0.2% [-2.0, +2.3] | -0.1% [-1.2, +1.0] |
| 4 | 10% | Tumor 30% high, 20% CV | +0.5% [-1.8, +3.2] | +2.2% [-0.1, +4.5] | -1.3% [-2.1, -0.6] |
| 7.5 | 5% | True $v_p$ | -3.3% [-5.3, -1.6] | -0.5% [-2.1, +1.2] | -0.1% [-0.7, +0.5] |
| 7.5 | 5% | Unbiased, 20% CV | -2.7% [-4.6, -1.0] | +0.1% [-1.6, +1.8] | -0.1% [-0.7, +0.6] |
| 7.5 | 5% | Tumor 30% low, 20% CV | -3.8% [-5.6, -2.1] | -0.5% [-2.3, +1.4] | +0.4% [-0.3, +1.2] |
| 7.5 | 5% | Tumor 30% high, 20% CV | -1.8% [-3.8, -0.01] | +0.3% [-1.6, +2.2] | -0.4% [-1.0, +0.2] |
| 7.5 | 10% | True $v_p$ | -6.1% [-8.9, -3.4] | -0.1% [-2.8, +2.7] | -0.4% [-0.9, +0.1] |
| 7.5 | 10% | Unbiased, 20% CV | -4.9% [-7.7, -2.3] | +0.6% [-2.3, +3.5] | -0.3% [-0.9, +0.2] |
| 7.5 | 10% | Tumor 30% low, 20% CV | -7.3% [-9.7, -5.0] | -1.2% [-3.7, +1.4] | +0.4% [-0.2, +1.2] |
| 7.5 | 10% | Tumor 30% high, 20% CV | -2.4% [-5.5, +0.6] | +2.1% [-1.1, +5.4] | -0.9% [-1.5, -0.4] |

Each condition generated 1,200 tumor targets across 20 seeds; 1,199–1,200 had successful VI fits in both arms in each comparison. Each arm had at most one tumor fallback (0.08% of retained fits).

For Extended Tofts, the unbiased prior slightly reduced $v_p$ RMSE, most at 10% noise, without any change detected in $K^{trans}$ or $v_e$.

#### Supplementary Table S12c | Patlak prior sensitivity across temporal resolutions.

| Temporal resolution (s) | DCE noise | Prior scenario | $v_p$ RMSE change [95% CI] | $K^{trans}$ RMSE change [95% CI] |
|---|---|---|---|---|
| 2 | 5% | True $v_p$ | -0.4% [-1.0, +0.2] | -0.6% [-0.8, -0.4] |
| 2 | 5% | Unbiased, 20% CV | -0.4% [-1.0, +0.1] | -0.6% [-0.8, -0.4] |
| 2 | 5% | Tumor 30% low, 20% CV | -0.2% [-0.6, +0.1] | -0.4% [-0.6, -0.2] |
| 2 | 5% | Tumor 30% high, 20% CV | -0.5% [-1.1, +0.2] | -0.7% [-0.9, -0.5] |
| 2 | 10% | True $v_p$ | -2.0% [-3.1, -0.9] | -1.8% [-2.1, -1.4] |
| 2 | 10% | Unbiased, 20% CV | -2.1% [-3.1, -1.0] | -1.8% [-2.1, -1.3] |
| 2 | 10% | Tumor 30% low, 20% CV | -1.3% [-2.0, -0.5] | -1.1% [-1.4, -0.8] |
| 2 | 10% | Tumor 30% high, 20% CV | -2.4% [-3.6, -1.1] | -2.1% [-2.6, -1.7] |
| 4 | 5% | True $v_p$ | -1.1% [-1.7, -0.5] | -0.8% [-1.1, -0.5] |
| 4 | 5% | Unbiased, 20% CV | -1.1% [-1.7, -0.4] | -0.8% [-1.1, -0.5] |
| 4 | 5% | Tumor 30% low, 20% CV | -0.8% [-1.2, -0.4] | -0.6% [-0.8, -0.3] |
| 4 | 5% | Tumor 30% high, 20% CV | -1.2% [-1.9, -0.4] | -0.9% [-1.2, -0.5] |
| 4 | 10% | True $v_p$ | -3.9% [-4.9, -2.7] | -2.2% [-2.7, -1.8] |
| 4 | 10% | Unbiased, 20% CV | -3.8% [-5.0, -2.5] | -2.2% [-2.7, -1.8] |
| 4 | 10% | Tumor 30% low, 20% CV | -2.7% [-3.3, -1.9] | -1.6% [-2.0, -1.2] |
| 4 | 10% | Tumor 30% high, 20% CV | -4.4% [-5.6, -2.9] | -2.7% [-3.3, -2.1] |
| 7.5 | 5% | True $v_p$ | -3.1% [-3.9, -2.2] | -1.8% [-2.1, -1.4] |
| 7.5 | 5% | Unbiased, 20% CV | -3.0% [-3.8, -2.2] | -1.8% [-2.1, -1.4] |
| 7.5 | 5% | Tumor 30% low, 20% CV | -2.3% [-2.9, -1.8] | -1.2% [-1.5, -1.0] |
| 7.5 | 5% | Tumor 30% high, 20% CV | -3.6% [-4.6, -2.5] | -2.1% [-2.5, -1.7] |
| 7.5 | 10% | True $v_p$ | -8.7% [-10.2, -7.3] | -4.3% [-4.8, -3.8] |
| 7.5 | 10% | Unbiased, 20% CV | -8.5% [-10.0, -7.1] | -4.3% [-4.8, -3.8] |
| 7.5 | 10% | Tumor 30% low, 20% CV | -6.3% [-7.2, -5.3] | -2.9% [-3.2, -2.5] |
| 7.5 | 10% | Tumor 30% high, 20% CV | -10.1% [-11.8, -8.3] | -5.2% [-5.8, -4.6] |

All 1,200 tumor targets per condition contributed to each comparison, with no VI fallbacks.

For Patlak, the unbiased prior reduced $K^{trans}$ and $v_p$ RMSE, most at 7.5 s and 10% noise.

**Supplementary Table S12d | Explicit-2CXM prior sensitivity across temporal resolutions.**

| Temporal resolution (s) | DCE noise | Prior scenario | $v_p$ RMSE change [95% CI] | $K^{trans}$ RMSE change [95% CI] | $v_e$ RMSE change [95% CI] |
|---|---|---|---|---|---|
| 2 | 5% | True $v_p$ | -45.8% [-58.5, -25.1] | -36.6% [-46.1, -22.2] | -19.1% [-25.6, -10.6] |
| 2 | 5% | Unbiased, 20% CV | -41.2% [-58.1, -12.5] | -33.3% [-45.3, -13.8] | -16.8% [-25.4, -3.7] |
| 2 | 5% | Tumor 30% low, 20% CV | -49.9% [-56.1, -42.7] | -36.4% [-44.7, -26.3] | -20.0% [-24.3, -15.7] |
| 2 | 5% | Tumor 30% high, 20% CV | -23.4% [-48.5, +9.6] | -22.2% [-36.1, -4.2] | -12.2% [-22.5, +0.7] |
| 2 | 10% | True $v_p$ | -55.3% [-59.7, -50.3] | -37.8% [-42.4, -32.8] | -11.3% [-14.6, -8.3] |
| 2 | 10% | Unbiased, 20% CV | -50.3% [-58.6, -37.1] | -35.1% [-41.8, -25.8] | -10.4% [-14.2, -6.8] |
| 2 | 10% | Tumor 30% low, 20% CV | -56.4% [-60.8, -51.4] | -39.1% [-44.6, -32.4] | -10.6% [-13.9, -7.5] |
| 2 | 10% | Tumor 30% high, 20% CV | -43.9% [-51.3, -35.2] | -28.1% [-33.6, -22.2] | -9.3% [-13.1, -6.0] |
| 4 | 5% | True $v_p$ | -54.7% [-59.0, -50.3] | -38.6% [-43.8, -33.3] | -17.7% [-21.7, -13.9] |
| 4 | 5% | Unbiased, 20% CV | -43.9% [-54.8, -28.1] | -33.7% [-39.9, -27.4] | -13.1% [-19.6, -4.4] |
| 4 | 5% | Tumor 30% low, 20% CV | -51.0% [-56.5, -44.6] | -40.4% [-46.0, -34.5] | -16.5% [-20.2, -13.0] |
| 4 | 5% | Tumor 30% high, 20% CV | -35.4% [-50.6, -15.5] | -27.7% [-34.6, -20.3] | -10.7% [-18.4, -0.8] |
| 4 | 10% | True $v_p$ | -55.4% [-58.9, -52.0] | -35.0% [-39.1, -31.2] | -8.9% [-12.6, -5.7] |
| 4 | 10% | Unbiased, 20% CV | -44.7% [-53.8, -31.4] | -31.2% [-35.8, -26.8] | -6.6% [-11.1, -2.0] |
| 4 | 10% | Tumor 30% low, 20% CV | -57.8% [-60.3, -55.3] | -38.8% [-43.0, -34.7] | -8.6% [-12.5, -5.2] |
| 4 | 10% | Tumor 30% high, 20% CV | -31.8% [-44.5, -16.8] | -22.6% [-27.5, -17.6] | -5.2% [-9.8, -0.2] |

| Temporal resolution (s) | DCE noise | Prior scenario | $v_p$ RMSE change [95% CI] | $K^{trans}$ RMSE change [95% CI] | $v_e$ RMSE change [95% CI] |
|---|---|---|---|---|---|
| 7.5 | 5% | True $v_p$ | -54.0% [-58.4, -49.1] | -37.6% [-41.4, -33.9] | -14.2% [-18.6, -10.0] |
| 7.5 | 5% | Unbiased, 20% CV | -43.9% [-54.8, -27.4] | -33.7% [-38.1, -28.9] | -8.9% [-16.4, +1.9] |
| 7.5 | 5% | Tumor 30% low, 20% CV | -53.9% [-57.5, -50.2] | -40.2% [-44.7, -35.7] | -13.3% [-17.8, -9.0] |
| 7.5 | 5% | Tumor 30% high, 20% CV | -34.2% [-49.6, -13.8] | -26.1% [-32.6, -19.3] | -5.2% [-13.7, +6.7] |
| 7.5 | 10% | True $v_p$ | -55.3% [-59.0, -51.5] | -31.7% [-35.3, -28.0] | -7.5% [-11.1, -4.5] |
| 7.5 | 10% | Unbiased, 20% CV | -48.3% [-54.3, -40.5] | -29.2% [-32.6, -25.9] | -5.8% [-9.3, -2.3] |
| 7.5 | 10% | Tumor 30% low, 20% CV | -58.3% [-61.4, -55.1] | -35.1% [-39.6, -30.5] | -7.2% [-10.9, -3.8] |
| 7.5 | 10% | Tumor 30% high, 20% CV | -34.5% [-44.8, -22.0] | -22.0% [-26.8, -17.2] | -2.4% [-6.4, +2.3] |

The table includes all paired successful VI fits. BLS retained 869–1,065 of 1,200 generated targets per condition, of which 869–1,063 had successful VI fits in both arms per prior comparison. Each arm had 0–3 tumor fallbacks (at most 0.3% of retained fits).

For explicit 2CXM, the prior reduced $K^{trans}$ and $v_p$ RMSE substantially across all paired fits, least with tumor centers 30% high. These means were sensitive to isolated fits: removing a single voxel shifted mean $v_p$ results by up to 12.3 percentage points. In one such voxel at 2 s and 5% noise (true $v_p$ 0.038), default-prior VI gave 0.040, whereas VI with the prior set to true $v_p$ gave 0.195; this voxel was not well constrained and is not in Table S12e.

**Supplementary Table S12e | Explicit-2CXM prior sensitivity in well-constrained fits.**

| Temporal resolution (s) | DCE noise | Prior scenario | $v_p$ RMSE change [95% CI] | $K^{trans}$ RMSE change [95% CI] | $v_e$ RMSE change [95% CI] |
|---|---|---|---|---|---|
| 2 | 5% | True $v_p$ | -25.5% [-33.9, -17.9] | -19.1% [-27.9, -11.4] | -3.8% [-7.8, -0.7] |
| 2 | 5% | Unbiased, 20% CV | -26.6% [-34.7, -19.5] | -19.9% [-28.7, -12.3] | -3.9% [-8.0, -0.7] |
| 2 | 5% | Tumor 30% low, 20% CV | -30.5% [-39.9, -21.5] | -22.1% [-31.3, -13.9] | -3.9% [-8.3, -0.4] |

| Temporal resolution (s) | DCE noise | Prior scenario | $v_p$ RMSE change [95% CI] | $K^{trans}$ RMSE change [95% CI] | $v_e$ RMSE change [95% CI] |
|---|---|---|---|---|---|
| 2 | 5% | Tumor 30% high, 20% CV | -16.9% [-25.6, -9.2] | -9.0% [-19.5, +0.6] | -2.2% [-6.2, +0.9] |
| 2 | 10% | True $v_p$ | -22.7% [-31.8, -13.9] | -11.6% [-18.8, -4.6] | +1.1% [-1.5, +3.4] |
| 2 | 10% | Unbiased, 20% CV | -23.1% [-31.8, -15.2] | -11.5% [-18.6, -4.8] | +0.8% [-1.7, +2.9] |
| 2 | 10% | Tumor 30% low, 20% CV | -29.6% [-38.5, -20.8] | -14.1% [-21.7, -6.7] | +0.9% [-1.7, +3.5] |
| 2 | 10% | Tumor 30% high, 20% CV | -12.8% [-23.6, -1.8] | -4.0% [-12.1, +4.3] | +1.6% [-1.3, +4.4] |
| 4 | 5% | True $v_p$ | -17.1% [-24.3, -10.5] | -9.5% [-13.8, -5.5] | +1.0% [-1.3, +2.8] |
| 4 | 5% | Unbiased, 20% CV | -12.8% [-22.5, -1.6] | -6.8% [-12.3, +0.2] | +1.6% [-1.0, +3.8] |
| 4 | 5% | Tumor 30% low, 20% CV | -14.6% [-26.7, +1.8] | -12.1% [-18.5, -5.4] | +1.5% [-1.0, +3.6] |
| 4 | 5% | Tumor 30% high, 20% CV | -1.1% [-15.1, +16.6] | +0.8% [-7.2, +11.2] | +2.5% [-0.3, +5.4] |
| 4 | 10% | True $v_p$ | -17.3% [-24.3, -10.5] | -2.1% [-6.6, +2.7] | +2.9% [+1.1, +4.4] |
| 4 | 10% | Unbiased, 20% CV | -16.6% [-23.0, -10.7] | -1.0% [-5.6, +4.2] | +2.9% [+1.3, +4.5] |
| 4 | 10% | Tumor 30% low, 20% CV | -22.9% [-30.3, -15.4] | -2.3% [-9.0, +4.7] | +3.6% [+1.5, +5.5] |
| 4 | 10% | Tumor 30% high, 20% CV | -9.0% [-17.4, -1.2] | -0.4% [-4.4, +3.5] | +2.4% [+0.9, +3.8] |
| 7.5 | 5% | True $v_p$ | -19.0% [-24.8, -13.8] | -9.6% [-15.0, -4.2] | +1.2% [-1.8, +3.7] |
| 7.5 | 5% | Unbiased, 20% CV | -16.8% [-22.9, -11.2] | -7.3% [-13.3, -1.0] | +1.4% [-1.6, +3.9] |
| 7.5 | 5% | Tumor 30% low, 20% CV | -21.8% [-28.4, -14.5] | -10.2% [-17.3, -1.9] | +1.5% [-2.0, +4.5] |
| 7.5 | 5% | Tumor 30% high, 20% CV | -13.5% [-19.3, -8.6] | -7.4% [-12.2, -3.4] | +1.0% [-1.7, +3.3] |
| 7.5 | 10% | True $v_p$ | -16.9% [-23.7, -11.1] | -5.0% [-10.4, +0.2] | +2.9% [+0.6, +4.9] |
| 7.5 | 10% | Unbiased, 20% CV | -13.7% [-20.3, -7.5] | -3.6% [-8.5, +0.9] | +3.2% [+1.0, +5.1] |
| 7.5 | 10% | Tumor 30% low, 20% CV | -22.5% [-30.3, -14.9] | -6.3% [-14.2, +2.0] | +3.7% [+1.3, +5.8] |

| Temporal resolution (s) | DCE noise | Prior scenario | $v_p$ RMSE change [95% CI] | $K^{trans}$ RMSE change [95% CI] | $v_e$ RMSE change [95% CI] |
|---|---|---|---|---|---|
| 7.5 | 10% | Tumor 30% high, 20% CV | -7.7% [-15.8, -0.1] | -2.2% [-6.7, +1.6] | +2.3% [+0.1, +4.2] |

The table includes well-constrained paired successful VI fits: 296–658 of 1,200 generated targets per prior comparison.

Where the shared BLS fit was well constrained, reductions were smaller, and $K^{trans}$ benefit was clearest at 2 s. At 10% noise, $v_e$ RMSE increased slightly at 4 and 7.5 s. These experiments concern explicit 2CXM; automatic eligibility is limited to temporal resolutions of 2 s or finer.

*L.2 Signal-level recovery and calibration errors*

These experiments tested whether priors derived from simulated DSC signal, rather than set directly as in Tables S12b–e, reduced DCE error, and whether errors in the assumed NAWM $v_p$ of 0.010 increased it. DSC signal from 1,000 NAWM and 60 tumor voxels per seed was converted to concentration, BSW-corrected and used to estimate AUC-based CBV and the NAWM-normalized prior. DSC temporal resolution was 1.5 s and TE was 30 ms. DSC curves used exact convolution of a piecewise-linear AIF (0.01-s grid) with an exponential residue. Five conditions tested signal noise and leakage (Table S12f). All five used the same Extended-Tofts DCE curves, generated as in L.1 with 4-s temporal resolution and 5% noise but different random seeds. VI with priors set to true $v_p$ was fitted to the same curves for comparison.

**Supplementary Table S12f | Signal-level recovery of the DSC-informed DCE prior.**

| DSC condition | Prior / true $v_p$ | $v_p$ RMSE change [95% CI] | $K^{trans}$ RMSE change [95% CI] | $v_e$ RMSE change [95% CI] |
|---|---|---|---|---|
| Noiseless DSC | 0.917 | -1.8% [-3.5, -0.3] | -0.3% [-1.4, +0.9] | -0.8% [-1.4, -0.2] |
| DSC SNR 50 | 0.924 | -1.7% [-3.3, -0.2] | -0.2% [-1.2, +1.0] | -0.7% [-1.4, -0.1] |
| DSC SNR 20 | 0.920 | -1.4% [-3.0, +0.1] | +0.2% [-0.9, +1.4] | -0.7% [-1.4, -0.1] |
| SNR 50 + T1 leakage | 0.922 | -1.7% [-3.4, -0.2] | -0.2% [-1.2, +1.0] | -0.7% [-1.4, -0.1] |
| SNR 50 + positive tail | 0.930 | -1.8% [-3.3, -0.3] | -0.2% [-1.2, +1.0] | -0.8% [-1.4, -0.1] |

All 1,200 tumor targets had successful VI fits in both arms.

Without DSC noise, prior means were 0.92 of true $v_p$, as BSW correction attributed the longer tumor transit times (4–8 s versus 4–5.5 s in NAWM) to negative K2, reducing the estimated CBV (Section B).

The prior reduced $v_p$ and $v_e$ RMSE slightly, with no detected change in $K^{trans}$; only the $v_p$ interval at DSC SNR 20 included zero. Priors set to true $v_p$ reduced $v_p$ RMSE by 1.9% on the same curves; compared directly, $v_p$ RMSE with the DSC SNR 50 prior was 0.2% higher.

The calibration experiment varied true NAWM $v_p$ while retaining the fixed 0.010 conversion from nCBV to the prior mean, using DSC SNR 50 and the same 4-s DCE protocol. As a control, prior values were randomly reassigned among tumor voxels, preserving their distribution.

**Supplementary Table S12g | Absolute NAWM $v_p$ calibration sensitivity.**

| True NAWM $v_p$ center | Prior / true $v_p$ | Matched $v_p$ RMSE change [95% CI] | Reassigned $v_p$ RMSE change [95% CI] | Matched $K^{trans}$ RMSE change [95% CI] | Matched $v_e$ RMSE change [95% CI] |
|---|---|---|---|---|---|
| 0.007 | 1.319 | -1.1% [-2.6, +0.5] | +2.5% [+0.9, +3.9] | -0.1% [-1.1, +1.0] | -1.1% [-1.7, -0.5] |
| 0.010 | 0.924 | -1.7% [-3.4, -0.3] | +1.2% [+0.1, +2.3] | -0.2% [-1.2, +1.0] | -0.7% [-1.4, -0.1] |
| 0.015 | 0.616 | -1.0% [-2.8, +0.8] | -0.7% [-2.7, +1.0] | +0.5% [-0.9, +2.0] | -0.3% [-1.0, +0.4] |

Each comparison included 1,199–1,200 tumor targets with successful VI fits in both arms. The default-prior and matched-prior arms had no fallbacks; each reassigned arm had at most one (0.08% of retained fits). In Tables S12f/g, prior/true-$v_p$ ratios are means across seeds of the median ratio over all generated tumor targets; RMSE changes use paired successful VI fits.

A $v_p$ benefit was detected only at the 0.010 calibration value, and reassignment increased $v_p$ error at the two lower values. With either incorrect calibration, $v_p$ RMSE was about 1% higher than with priors set to true $v_p$, and no increase in error relative to default VI was detected.

Signal-derived priors were also tested with explicit 2CXM at 2-s temporal resolution and 5% noise, reusing the L.1 tumor curves, BLS fits, selected delays and VI fits with priors set to true $v_p$. DSC curves were generated as for Extended Tofts, with transit time $6000 \times v_p/F_p$ seconds, using true $F_p$ in mL/min/100 mL. The 1,000-voxel NAWM calibration pool kept the original 100 NAWM voxels and added 900 with independent parameters and noise. DSC signal was simulated at SNR 50 without leakage and with a fixed T1 leakage term (K2 = 0.05) that was not tied to the simulated permeability.

**Supplementary Table S12h | Signal-derived priors in well-constrained explicit-2CXM fits.**

| Prior | $v_p$ RMSE change [95% CI] | $K^{trans}$ RMSE change [95% CI] | $v_e$ RMSE change [95% CI] |
|---|---|---|---|
| True $v_p$ | -25.5% [-34.2, -18.0] | -19.1% [-27.8, -11.7] | -3.8% [-7.9, -0.7] |
| DSC SNR 50, no leakage | -27.3% [-36.5, -19.2] | -20.7% [-29.2, -13.3] | -3.9% [-7.9, -0.8] |
| DSC SNR 50, imposed leakage | -27.4% [-36.5, -19.2] | -20.7% [-29.2, -13.3] | -3.9% [-7.9, -0.8] |

Changes are relative to default VI at 2-s temporal resolution and 5% DCE noise, using 658 paired successful VI fits where the shared BLS fit was well constrained.

DSC prior centers had a relative RMSE of 14.7–14.8%. The mean across seeds of the median prior/true ratio was 0.98 overall, 1.12 for transit times below 2 s and 0.90 for longer transit times. BSW correction recovered the imposed leakage (median tumor K2 0.048), and results with and without leakage were similar. In well-constrained fits, no difference in $v_p$ RMSE was detected between signal-derived priors and priors set to true $v_p$ (−2.4% with and without leakage). Across all 1,061 fits that succeeded with all three priors, including poorly constrained fits, $v_p$ RMSE with signal-derived priors was higher by 5.4% without and 5.5% with imposed leakage; as in Table S12d, these results were sensitive to isolated values.

*L.3 Paired clinical analysis*

In 11 subjects, VI with the DSC prior enabled or skipped used the same preprocessed inputs, BLS model, delay and starting estimates. Clinical summaries included finite paired retained outputs, including BLS fallbacks; posterior-SD comparisons required successful VI fits in both arms.

**Supplementary Table S12i | Clinical use and fit quality of the DSC-informed DCE prior.**

| Metric | Subject median | Subject range |
|---|---:|---:|
| Retained fits using prior | 65.8% | 53.5% to 72.1% |
| VI-to-BLS fallback | 3.9% | 2.2% to 4.9% |
| Fits within 0.5 prior SD | 27.4% | 23.0% to 35.4% |
| Fits beyond 2 prior SD | 11.8% | 7.3% to 17.2% |
| Median $R^2$, prior skipped | 0.559 | 0.430 to 0.659 |
| Median $R^2$, DSC prior enabled | 0.558 | 0.428 to 0.658 |

| Metric | Subject median | Subject range |
|---|---|---|
| Paired change in median $R^2$ | -0.0015 | -0.0026 to 0.0016 |
| Median absolute difference from prior mean (prior SDs) | 0.845 | 0.715 to 0.943 |

VI-to-BLS fallback is the fraction of retained voxels; with the prior skipped, it was 3.2% (range, 1.3–4.3%). Prior-departure rows include fits that used the prior, with distances expressed in prior SDs.

**Supplementary Table S12j | Clinical regional parameter changes with the DSC-informed DCE prior.**

| Region | Parameter | Median value change | Median absolute voxel change | Median posterior-SD change |
|---|---|---|---|---|
| whole brain | $v_p$ | -3.3% | 0.000191 | -0.5% |
| whole brain | $K^{trans}$ | 1.5% | 3.37e-05 | 0.5% |
| whole brain | $v_e$ | -0.1% | 0 | -0.2% |
| enhancing tumor | $v_p$ | -1.6% | 0.000303 | 0.4% |
| enhancing tumor | $K^{trans}$ | 0.5% | 0.000176 | 0.4% |
| enhancing tumor | $v_e$ | -0.2% | 0.000417 | -0.3% |
| non-enhancing tumor | $v_p$ | -17.9% | 0.000899 | -4.6% |
| non-enhancing tumor | $K^{trans}$ | 4.4% | 0.000108 | 0.5% |
| non-enhancing tumor | $v_e$ | 0% | 0.000151 | -0.5% |
| NAWM | $v_p$ | -16.2% | 0.0008 | -1.3% |
| NAWM | $K^{trans}$ | 3.3% | 8.5e-05 | 0.9% |
| NAWM | $v_e$ | 4.5% | 0.000435 | 0% |

Entries are medians across subjects. Percentage changes compare regional medians with the prior enabled versus skipped; absolute voxel changes are in $min^{-1}$ for $K^{trans}$ and volume fractions for $v_p$ and $v_e$. Posterior-SD changes exclude BLS fallbacks.

The largest changes were $v_p$ decreases of 16.2% in NAWM and 17.9% in non-enhancing tumor; enhancing-tumor estimates changed little. Within the same DCE QC mask in both arms, these decreases were 9.3% and 4.5%. Without independent $v_p$

ground truth, these changes cannot establish improved accuracy. DSC–DCE agreement is also not independent evidence as DSC supplies the prior.

#### M. T1 estimation and B1+ correction

PMRIPrep's VFA T1 estimator and B1+ correction were evaluated using the QIBA T1 v03 and B1 Correction v13 objects [45,46]. T1 results are grouped by the nominal input ratio S0/σ, which is not the same as measured image SNR. Known B1+ offsets test whether a supplied correction map is applied correctly; they do not test estimation of a B1+ map from double-angle data.

##### QIBA T1 recovery and application of B1+ correction

**Supplementary Table S13-1 | QIBA T1 recovery by nominal input S0/σ.**

| Nominal input S0/σ | Median \|bias\| (%) | Intra-patch CV (%) | N patches |
|---|---|---|---|
| 5–10 | 59.9 | 200.1 | 8 |
| 10–30 | 56.1 | 243.5 | 40 |
| 30–100 | 24.7 | 104.9 | 32 |
| 100–300 | 4.7 | 26.4 | 88 |
| 300–1,000 | 1.4 | 9.2 | 48 |
| ≥1,000 | 0.2 | 1.8 | 120 |

The QIBA T1 DRO is a 15 (T1: 22–2,828 ms) × 7 (equilibrium signal S0: 500–50,000) patch grid at TR = 5 ms and six flip angles (3–35°), replicated at six magnitude-MR (Rician) noise levels. Recovered T1 was the median of the central 6×6 voxels in each patch; metrics aggregate over the DCE-relevant T1 window (250–2,828 ms). The primary axis is the nominal input S0-to-noise ratio, S0/σ. This is a dimensionless simulation control variable, not measured image SNR. Median absolute bias is reported because pooled signed summaries can obscure errors that vary with T1 and signal level.

**Supplementary Table S13-2 | QIBA T1 recovery by input noise.**

| σ (signal SD) | S0/σ range | Median \|bias\| (%) | CV (%) | Bias at 2,828 ms (%) | Bias at 250 ms (%) |
|---|---|---|---|---|---|
| 2 | 250–25,000 | 0.5 | 1.9 | 0.5 | 0 |
| 5 | 100–10,000 | 0.7 | 5.0 | -0.3 | 0 |
| 10 | 50–5,000 | 1.5 | 10.5 | -6.3 | 0 |
| 20 | 25–2,500 | 3.5 | 17.3 | -13.3 | 0 |
| 50 | 10–1,000 | 6.3 | 48.6 | -47.7 | 6.2 |
| 100 | 5–500 | 19.6 | 90.3 | -78.0 | -0.4 |

In the nominal S0/σ = 100–300 bin, the DCE-relevant T1 window is recovered with a median absolute bias of 4.7%, falling to 1.4% at S0/σ = 300–1,000 and 0.2% at S0/σ ≥ 1,000; below S0/σ ≈ 100 both bias and variance rise steeply. The degradation is concentrated at the longest T1 and lowest-S0/σ patches: even within the σ ≤ 20 regime the 2,828-ms corner reaches −13%, while bias at 250 ms remained near zero.

PMRIPrep uses WLLS initialization followed by nonlinear least-squares fitting in signal space.

**Supplementary Table S13-3 | QIBA B1+ correction.**

| Transmit offset | Uncorrected T1 (ms) | Uncorrected bias (%) | Corrected T1 (ms) | Corrected bias (%) |
|---|---|---|---|---|
| +10% | 1213.3 | +21.3 | 999.9 | -0.01 |
| +20% | 1448.5 | +44.8 | 1000.3 | +0.03 |
| -10% | 807.8 | -19.2 | 1000.3 | +0.03 |
| -20% | 636.1 | -36.4 | 1000.1 | +0.01 |

Both QIBA objects were generated with the same spoiled gradient-echo equation fitted by PMRIPrep. They therefore test the estimator under matched model assumptions and do not measure robustness to clinical model mismatch. The same WLLS-initialized NLLS estimator was applied directly to the phantom. Physiological T1-range and cerebrospinal fluid (CSF)/edge filters were omitted as they are not applicable to a phantom spanning 22–2,828 ms.

The B1-Correction object is a uniform phantom (true T1 = 1,000 ms) acquired under four transmit-field offsets. Its B1+ data use a double-angle method that PMRIPrep does not ingest, so the correction map was constructed from the known offset for each condition, independently of the fitted VFA signals. Without correction, the fits reproduced the expected flip-angle bias; supplying the known correction returned T1 to ground truth.

The uncorrected biases match the small-flip DESPOT1 prediction that T1 scales with the square of the flip-angle factor, $T1_{obs}/T1 = (1 + \delta)^2$ (+21/+45% for δ = +0.1/+0.2 and −19/−36% for δ = −0.1/−0.2). Supplying the exact actual-to-nominal flip-angle factor reduced these errors to <0.04%. This test confirms the sign and scaling of the correction; as the exact factor was supplied, it cannot establish the accuracy of a measured B1+ map or downstream $K^{trans}$.

#### N. Cross-software comparison: PMRIPrep and Siemens Spectra (Tissue4D)

We compared PMRIPrep and Spectra parametric maps voxelwise in one subject with glioblastoma acquired on a Siemens MAGNETOM Vida 3 T. Within enhancing tumor, NAWM-normalized CBV showed the strongest cross-software correspondence (Pearson $r = 0.993$; Spearman $\rho = 0.992$; Lin concordance correlation coefficient (CCC) = 0.991). The corresponding r, ρ, and CCC values were 0.846, 0.905, and 0.835 for $K^{trans}$ and 0.870, 0.827, and 0.700 for $v_p$. The median voxelwise Spectra/PMRIPrep $v_p$ ratio was 1.87.

For $v_e$ ($r = 0.820$, $\rho = 0.839$, and CCC = 0.788), fewer voxels were included in the comparison as $v_e$ is unavailable where automatic BIC selection retains Patlak. Spectra $v_e$ values appeared capped at 0.50, producing the horizontal accumulation in the scatter plot. Comparisons used finite, positive overlapping voxels without filtering by the PMRIPrep DCE QC mask.

PMRIPrep $K^{trans}$ contained vascular hotspots that were absent from the Spectra map. Spectra $v_p$ showed greater spatial variability, while the PMRIPrep $v_e$ map contained fewer positive voxels, as Patlak was selected in a subset of voxels. The undocumented Spectra fitting and masking settings preclude attributing these differences to a specific processing step. DCE correlations were lower in NAWM ($r = 0.31$–$0.69$), where parameter values were close to zero; NAWM nCBV agreement remained high ($r = 0.995$).

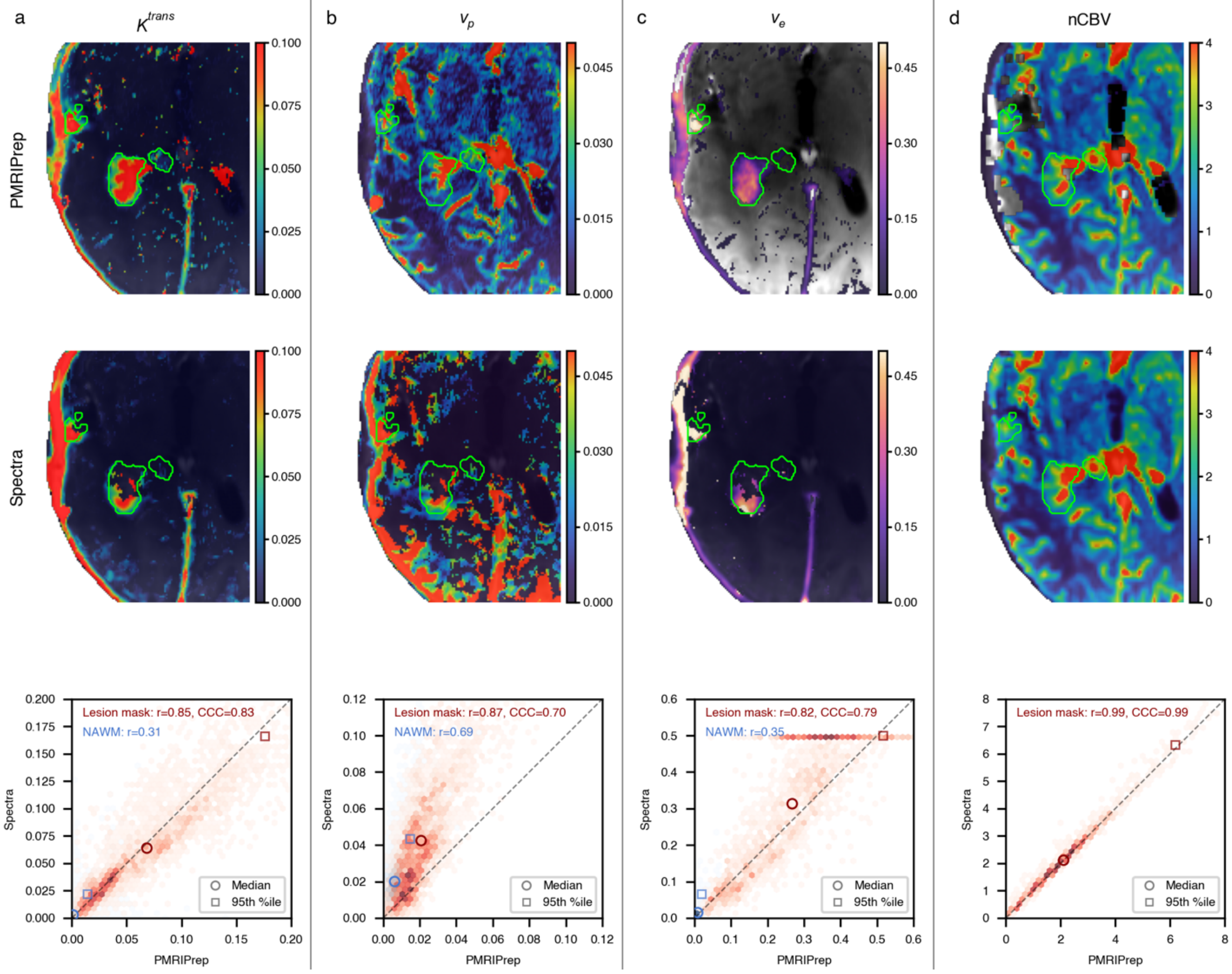


**Supplementary Figure S1 | Cross-software quantitative-map comparison.** Representative maps and voxelwise comparisons between PMRIPrep and Siemens Spectra (Tissue4D) in one glioblastoma subject. Columns show **a,** $K^{trans}$; **b,** $v_p$; **c,** $v_e$; and **d,** CBV normalized to normal-appearing white matter (nCBV). The upper row shows PMRIPrep maps, the middle row shows Spectra maps at matched axial locations, and the lower row shows voxelwise comparisons. Green contours delineate the enhancing-tumor ROI. DCE maps share the same native grid; nCBV maps from both packages were resampled to DCE space for display, and nCBV statistics were computed on the PMRIPrep DSC grid. The lower row compares positive finite values in enhancing tumor (red) and, for the DCE parameters, NAWM (blue), without filtering by the PMRIPrep DCE QC mask. Open circles and squares denote ROI medians and 95th percentiles, dashed lines indicate identity, and labels report the Pearson correlation coefficient (r), with the Lin CCC for enhancing tumor. Scatter plots use matched x and y limits and tick positions. Voxels beyond the displayed axis ranges remain included in the reported statistics. The horizontal accumulation in the Spectra $v_e$ comparison reflects its upper value of 0.50. Exact Spectra fitting, arterial input function, and masking settings were unavailable.

SELECT IMAGES FROM PMRIPREP QC REPORT

DESCRIPTION OF REPORT CONTENT

**a. Brain extraction**

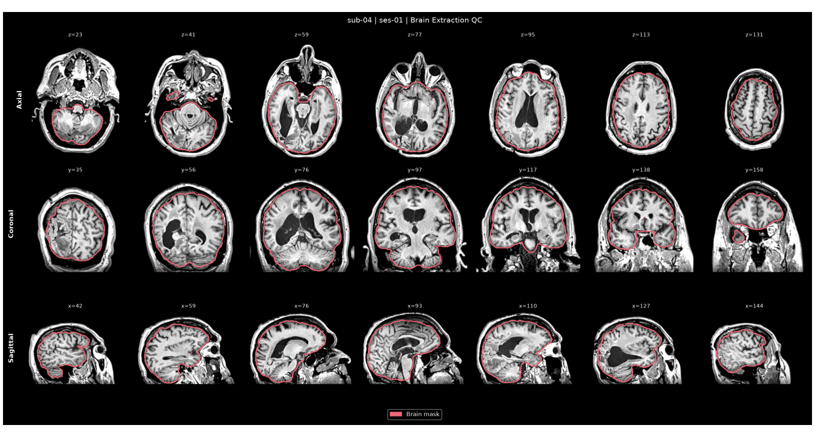

Brain extraction on T1w.

**b. Tumor segmentation**

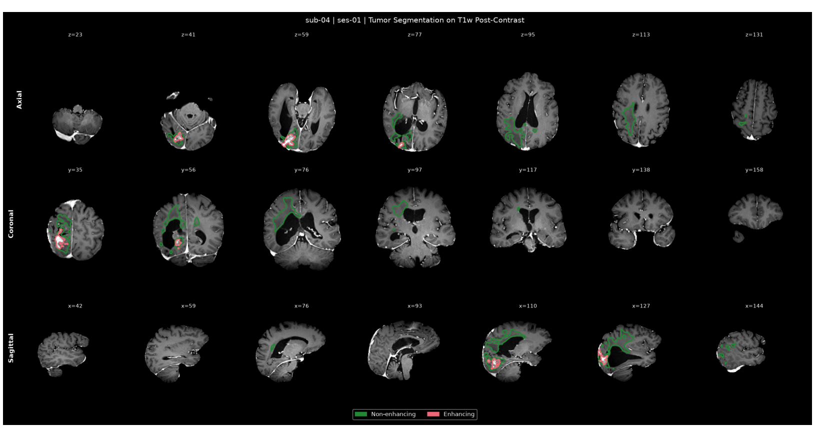

Automated tumor segmentation shown on post-contrast T1w. Green = non-enhancing tumor; red = enhancing tumor.

**c. NAWM reference ROI**

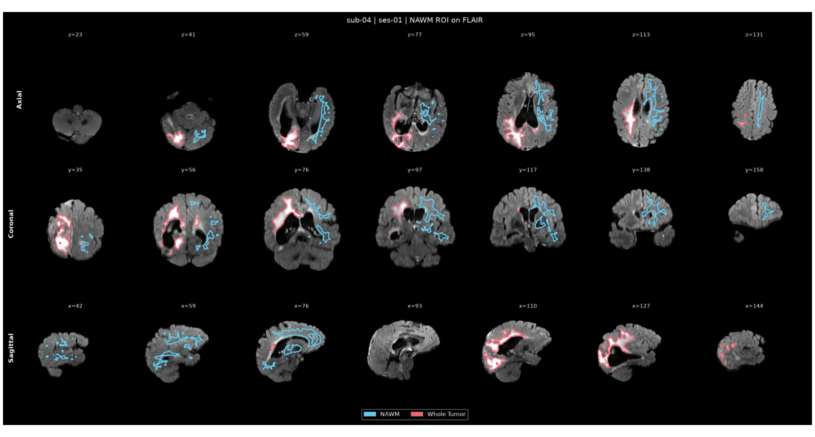

Normal-appearing white matter ROI (cyan) on FLAIR, placed contralateral to the tumor (red) when the tumor is lateralized. It provides the reference for nCBV and nCBF and, when qualified, anchors the DSC-informed DCE $v_p$ prior.

**d. DSC–T1w registration**

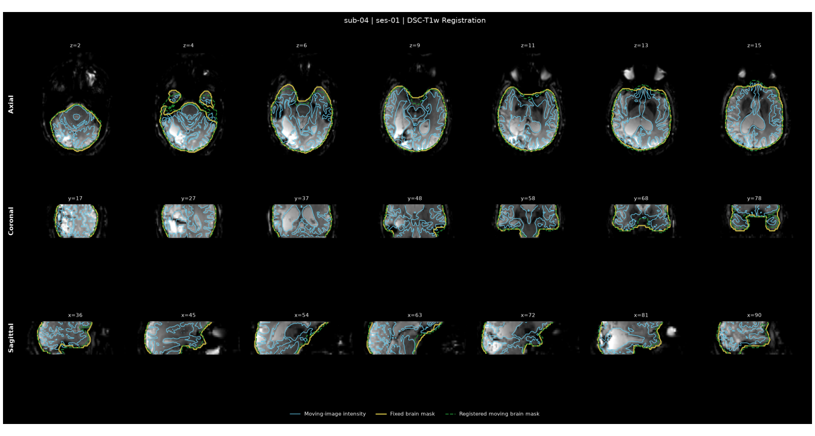

T1w anatomy resampled into DSC reference space using the inverse of the estimated DSC-to-T1w rigid transform (6 DOF). Contours show the native DSC brain mask and the resampled anatomical brain mask.

**e. DSC AIF detection**

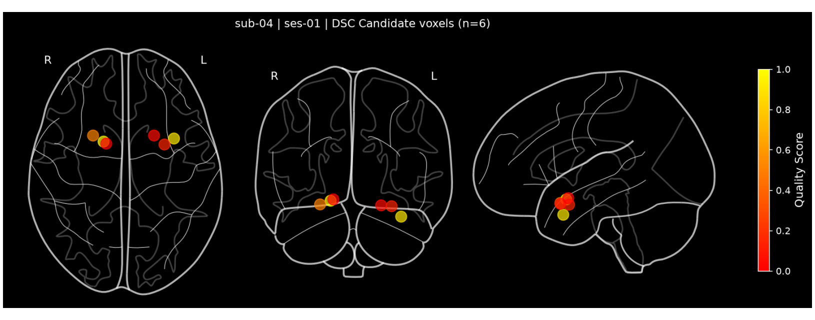


Glass-brain projection of AIF sources, color-coded by quality score (red = low, yellow = high).

**f. DSC leakage correction**

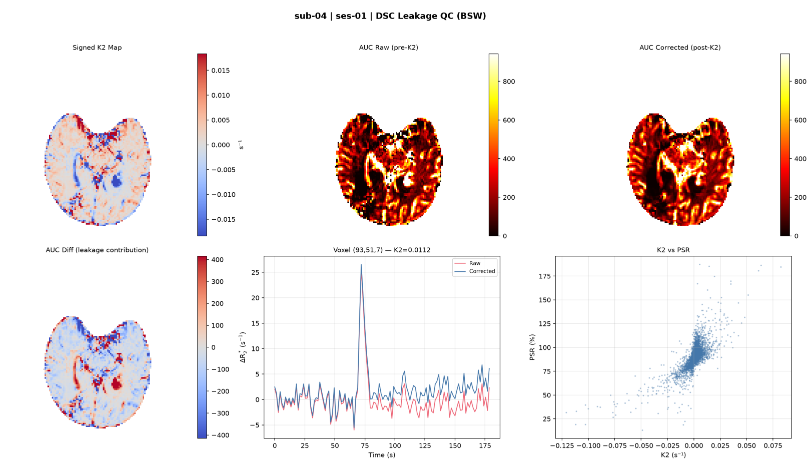


Boxerman–Schmainda–Weisskoff correction for contrast extravasation. Top row: $K_2$ map and tissue AUC before and after leakage correction. Bottom row: voxel-level difference, raw vs. corrected time course, and $K_2$ vs. PSR scatter.

**g. DSC representative voxel fit**

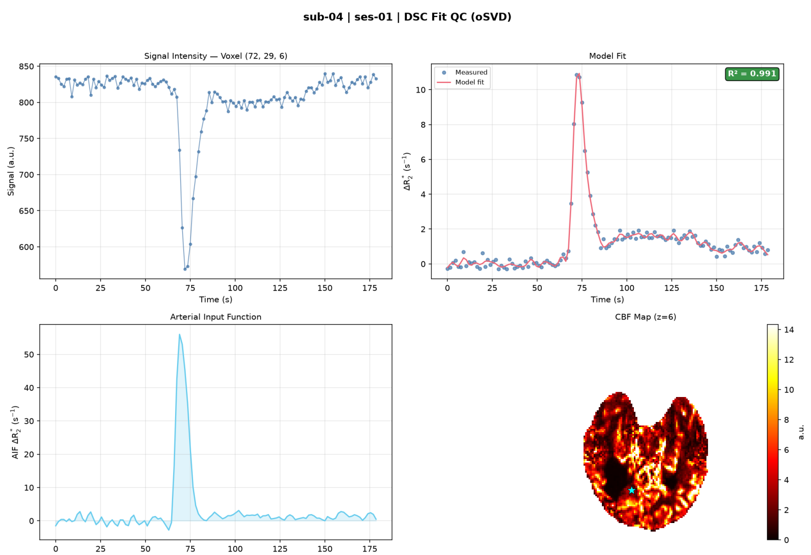


Representative voxel showing raw signal, concentration fit, AIF curve, and resulting rCBF map. Selection targets the median positive rCBF within enhancing tumor when available, with an eroded-brain fallback.

**h. DCE AIF detection**

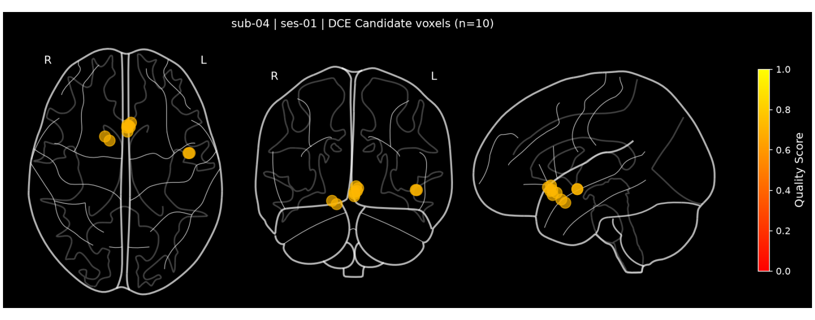


Glass-brain projection of DCE AIF sources, color-coded by quality score (red = low, yellow = high).

**i. DCE model selection**

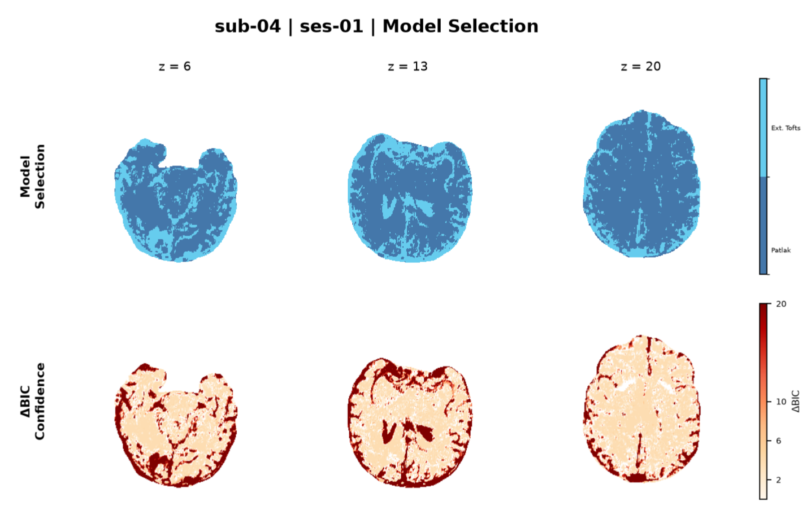


Top: per-voxel winning pharmacokinetic model. Bottom: ΔBIC confidence map. Regions with low ΔBIC indicate ambiguous model preference; high values indicate strong evidence for the selected model.

**j. DCE representative voxel fit**

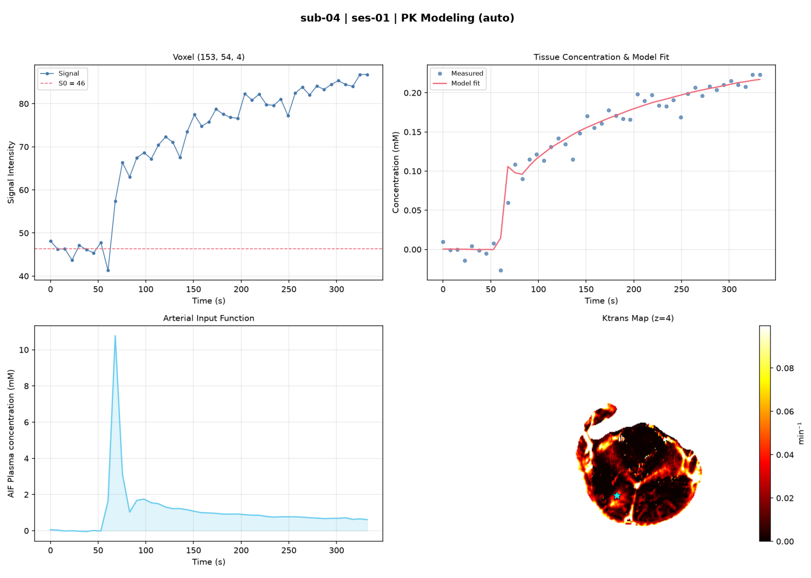


Representative voxel showing raw signal, concentration fit, AIF curve, and $K^{trans}$ map. Selection targets the median positive $K^{trans}$ within enhancing tumor when available, preferentially among voxels passing quantitative QC.

**Supplementary Figure S2 | Representative automated quality-control report images.** Selected QC figures from one clinical glioblastoma examination: **a,** brain extraction; **b,** tumor segmentation; **c,** normal-appearing white matter (NAWM) reference ROI; **d,** DSC-to-T1w registration; **e,** DSC arterial input function (AIF) detection; **f,** DSC leakage correction; **g,** representative DSC voxel fit; **h,** DCE AIF detection; **i,** DCE model selection; and **j,** representative DCE voxel fit. Images are shown at left and descriptions at right. PMRIPrep generates the applicable diagnostics for each subject-session according to the available modalities and executed workflow components.
